# Analyses of 37 composts revealed microbial taxa associated with disease suppressiveness

**DOI:** 10.1101/2025.01.17.633646

**Authors:** Anja Logo, Benedikt Boppré, Jacques Fuchs, Monika Maurhofer, Thomas Oberhänsli, Barbara Thürig, Franco Widmer, Johanna Mayerhofer, Pascale Flury

## Abstract

Compost is a valuable amendment for soil and potting substrate when it comes to suppressing soilborne pathogens. However, the effectiveness of different composts varies and can not yet be predicted. Microbial communities in compost play a key role in disease suppression, and therefore their composition or specific taxa may serve as indicators of suppressive composts. In this study, we investigated 37 composts from seven commercial compost producers to analyze the association of their bacterial and fungal communities with suppressive activity in three plant-pathogen systems: cress-*Globisporangium ultimum*, cucumber-*Globisporangium ultimum* and cucumber-*Rhizoctonia solani*. Our results underscore that compost suppressiveness is primarily pathogen-specific and, to a lesser extent, host-plant-specific. Suppressiveness was not correlated with physico-chemical properties, microbial activity, or the alpha-and beta-diversity of composts bacterial and fungal communities. Instead, microbial composition was largely shaped by producer-specific composting conditions and maturation processes, which were not necessarily linked to suppressive activity. A more nuanced comparison between the most and least suppressive composts revealed bacterial and few fungal taxa as potential indicators of suppressiveness for each plant-pathogen system. Notably, for *G. ultimum*-suppression, bacteria from the genera *Luteimonas*, *Sphingopyxis*, and *Algoriphagus* and for *R. solani* bacteria belonging to the phylum *Actinomycetota* emerged as promising candidates.

**Importance:** Soilborne diseases are a major yield-limiting factor in agricultural crop production world-wide, particularly in seedling cultivation. Their control remains a significant challenge and still largely relies on chemical fumigation of soils and steam sterilization of potting substrates. While chemical fumigants are increasingly criticized for their negative environmental impact, sterilization practices in general disrupt beneficial microbial com-munities, making substrates more susceptible to pathogen (re)-infestation. Amending soil or potting substrate with disease-suppressive compost offers a promising alternative. However, the targeted use of compost for plant protection is hindered by variable effectiveness and the lack of reliable tools to identify effective composts. This study provides a comprehensive abiotic and biotic characterization of compost, enabling a detailed anal-ysis of the properties associated with suppressiveness. The identification of bacterial and fungal taxa indicative of disease-suppressive composts lays the groundwork for targeted isolation of microorganisms and functional studies, with the ultimate aim of predicting and optimizing compost-mediated disease suppression.

## 1. Introduction

Soilborne diseases are a persistent challenge in agricultural crop production. Particularly, damping-off of seedlings, caused by fungi and fungus-like organisms such as *Rhizoctonia* spp. and *Globisporangium* spp., is a major yield constraint in both nurseries and fields (Lamichhane et al., 2017). Current disease management strategies, especially in organic agriculture, are limited and rely primarily on soil or substrate disinfestation to eradicate pathogens (Gullino et al., 2022). However, these practices can negatively impact (soil) microbial communities (Castellano-Hinojosa et al., 2022). With growing recognition of the essential role of beneficial microorganisms in agroecosystems (Hartmann and Six, 2023), there is an urgent need to develop more sustainable disease management approaches. One such strategy is the promotion of disease-suppressive condi-tions—environments where pathogens either fail to establish or persist, or are present but cause little to no damage to the host plant (Baker and Cook, 1974). The use of compost in soils or soiless substrates (e.g., peat) can promote disease suppressiveness and enhance overall plant health, offering a promising and environmentally sustainable strategy for managing soilborne diseases (Hoitink and Kuter, 1986; Noble and Coventry, 2005; Termorshuizen et al., 2006; De Corato, 2020; Edlinger et al., 2025).

Suppressive activity of composts has been reported against a broad range of plant pathogens including fungi, oomycetes, bacteria, nematodes, and viruses (Neher et al., 2022; Lutz et al., 2020). However, efficacy varies from compost to compost and is often specific to plant-pathogen interactions (Termorshuizen et al., 2006; Bonanomi et al., 2010; Pane et al., 2011). Feedstock composition, compost maturity and stability, as well as the composting process, have been shown to influence the development of disease-suppressive composts (Scheuerell and Mahaffee, 2005; Termorshuizen et al., 2006; Zmora-Nahum et al., 2008). However, despite decades of compost research, the key factors driving disease-suppressive activity remain poorly understood (Hadar and Papadopoulou, 2012; Bouchtaoui et al., 2024). Moreover, we still lack reliable indicators to predict the suppressive potential of a given compost (Bonanomi et al., 2010; Neher et al., 2022). To enable the more effective and targeted use of compost in crop protection, it is essential to deepen our understanding of the mechanisms behind disease suppression and to develop reliable diagnostic tools.

It is widely acknowledged that microorganisms play a crucial role in disease suppression by composts (Hoitink and Kuter, 1986; Bonanomi et al., 2010; Mehta et al., 2014; De Corato, 2020). This is supported by studies showing that most composts lose their suppressiveness when sterilized by heat treatment or gamma irradiation (Noble and Coventry, 2005; De Corato et al., 2019), and that suppressiveness can be restored by reintroducing compost-associated microorganisms (Nelson et al., 1983; Cao et al., 2018). Moreover, microbial recolonization during the cooling phase of the composting process is essential for the development of suppressive activity (Scheuerell and Mahaffee, 2005). Compost microorganisms can suppress soilborne pathogens directly via predation, parasitism, antibiosis or competition and indirectly via the plant through inducing resistance and improving plant nutrition (Mehta et al., 2014). However, mechanisms of action have only been studied for a few genera of compost microorganisms (Lutz et al., 2020), while for the majority of the diverse bacteria and fungi present in composts, the effects on plant pathogens and crop health, as well as their mechanisms of action, remain unknown.

Although not yet fully understood, the central role of compost microorganisms in disease suppression suggests they could serve as indicators for identifying disease-suppressive composts. Thus, we proposed an integrated approach to identify beneficial microorganisms and key microbial consortia in disease-suppressive composts (Lutz et al., 2020). This approach combines standardized bioassays to asses suppressive activity with metagenomics to characterize the microbial communities of composts. In a later step, promising microorganisms are isolated and their role in disease suppression investigated. Studies following a similar approach, though based on small compost sets (n*<*10), have identified various bacterial and fungal groups associated with disease-suppressive composts (Yu et al., 2015; De Corato et al., 2018, 2019; Scotti et al., 2020; Hernández-Lara et al., 2022). However, particularly bacterial compost communities show high taxonomic variability across compost (Wang et al., 2020) and thus, large composts sets are needed to identify robust patterns.

In a previous study (Mayerhofer et al., 2021a), we applied the integrated approach to 17 composts targeting the pathogen *Globisporangium ultimum*, a globally distributed soilborne pathogen with a wide host range (Lamichhane et al., 2017). The composts were tested in a peat-based substrate, which is generally considered conducive to pathogen development, making it a suitable baseline for assessing the suppressive potential of compost amendments (Hoitink and Boehm, 1999). Moreover, peat remains widely used in seedling production despite environmental concerns over carbon emissions from peat extraction and its near non-renewable nature (Kitir et al., 2018). We assessed compost’s suppressive activity using cress as a model plant, and characterized the composts’ bacterial communities through amplicon sequencing of the 16S rRNA (V3–V4) gene region. This revealed a significant association between the bacterial community structure and suppressive activity, and identified 75 bacterial amplicon sequence variants (ASVs) as indicative of the most suppressive composts.

Building on this foundation, we aimed to validate the robustness of these bacterial indicators, and expand upon the scope of previous work by identifying fungal indicators, utilizing an unprecedented dataset of 37 new compost samples. To consider possible plant–pathogen specificity of compost’s suppressive activity, we extended the analysis beyond the cress–*G. ultimum* model system to include two additional plant–pathogen systems. To compare *G. ultimum*-suppression between host plants, we tested suppression also in cucumber, an important horticultural crop. To allow comparison between pathogens, we tested suppressiveness in cucumber also against the fungal pathogen *Rhizoctonia solani*, which likely shows a different response to compost application than *G. ultimum* (Termorshuizen et al., 2006). Finally, we constructed an extensive dataset by integrating bacterial and fungal community profiling with assessments of 14 compost properties. This integration enabled us to comprehensively explore the relationships between compost properties, microbial communities, and disease-suppressive activity across multiple plant-pathogen systems, ultimately identifying indicators of disease-suppressive composts.

## 2. Materials and Methods

### 2.1. Selection and collection of composts

Recognizing that composts derived from different feedstocks harbor distinct microbial communities, this study focused on composts primarily made from green waste (EstrellaGonzález et al., 2020; Mayerhofer et al., 2021a; Hernández-Lara et al., 2022). Samples were collected from seven large-scale composting sites (A-G) in Switzerland during four collection periods (I-IV): May, June, and September 2022, and April 2023. At each time point, two composts of differing maturation stages were collected from sites A–E, when possible; additional samples were sourced from sites F and G as needed. Sites A, B and F compost in tabular windrows (up to 3.5 m high), while sites C, D, E and G used triangular windrows (up to 2 m high, 3-4 m wide). Composting was done on paved ground, except for site E, where composting took place along agricultural fields directly on soil. Composting processes and feedstocks are detailed in Tables S1 and S2. Three composts (Ex 1-3) were excluded from the data set due their negative effect on cress shoot growth in the absence of pathogen application (Figure S1), resulting in a final data set of 37 composts.

Three to five subsamples of each compost pile were pooled and sieved through a 10 mm mesh for bioassays and physicochemical analyses. For assessing maximum water holding capacity (WHC), microbial activity and microbial communities, composts were further sieved through a 2 mm mesh. Most analyses were initiated within one week of collection. Due to logistical constrains, samples used for microbial activity analysis and *R. solani* bioassays were stored at 5°C for two weeks.

### 2.2. Physicochemical properties and microbial activity of composts

Dry substance (DS) was determined by quantifying water loss after heating of 200 g of compost at 105°C for 24 h. To measure total nitrogen (N_tot_) and total organic carbon (C_org_), dried compost samples were milled with an impact rotor mill (Retsch, Germany) and sieved through a 1 mm mesh. Then, 500 mg each were analyzed using an elemental analyzer (Vario Max Cube, Elementar, Langenselbold, Germany) as described by Krause et al. (2022). The maximum WHC was assessed via capillary binding capacity using the reference method of Agroscope (2020). Nitrite (NO_2_^-^), nitrate (NO_3_^-^), ammonium (NH_4_^+^), phosphate (PO_4_^3-^) and pH were measured in a calcium chloride (CaCl_2_)-extract, while salinity and the content of soluble humic substances (OD_550_) were measured in a water-extract of the compost. Microbial activity was assessed using fluorescein diacetate (FDA) hydrolysis and basal respiration, each measured in three technical replicates (Ntougias et al., 2006; Alef and Nannipieri, 1995). Details on the methods are provided in the Supplementary Methods A and B. The total mineral nitrogen (N_min_) was calculated as the sum of NO_2_^-^, NO_3_^-^ and NH_4_^+^; the nitrification index (NO_3_^-^-N/N_min_) as NO_3_^-^ divided by N_min_; and the carbon-nitrogen ratio (C_org_/N_tot_) as C_org_ divided by N_tot_. The measured values for the physicochemical properties and the mean for the microbial activities are listed in Table S3.

### 2.3. Pot assays to assess disease suppression

A total of 37 composts were tested for their disease-suppressive activity in peat across three plant-pathogen systems: cress-*Globisporangium ultimum*, cucumber-*G. ultimum*, and *Rhizoctonia solani*-cucumber. The peat-compost mixtures are hereafter referred to as compost-substrates (1-37). Unamended peat served as the control in all experiments and is hereafter referred to as no-compost control. All substrates were tested at low and high pathogen concentrations, to account for variation in pathogen virulence, as well as without pathogen. With six replicate pots per treatment for cress and eight for cucumber, the total number of pots was 246 and 328, respectively.

More precisely, composts were mixed at a 1:4 ratio (v/v) with a peat-substrate (Einheitserde Typ 0, Einheitserdewerke Werkverband e.V., Sinntal-Altengronau, Germany) and fertilized with horn meal (2.30 g L^-1^ substrate, Biorga Hornmehl, Hauert, Grossaffoltern, Switzerland). The no-compost controls received also horn meal and were additionally supplemented with 1.94 g L^-1^ Granuphos (Landor, Muttenz, Switzerland) and 1.33 g L^-1^ Biorganic Kali-Magnesia (Hauert, Grossaffoltern, Switzerland) to compensate for the nutrients input from composts. All substrates were moistened with tap water (20%, v/v) and incubated at room temperature for one week prior to inoculation with pathogen.

Pathogens were cultured on Potato Dextrose Agar (PDA; Carl Roth, Karlsruhe, Ger-many) at 20°C for *G. ultimum* and at 24°C for *R. solani* (anastomosis group 4). For each experiment, pathogens were freshly cultured from long-term stocks. Long-term stocks consisted of mycelium-agar plugs in autoclaved ultra pure water (ELGA LabWater, Celle, Germany) stored at 20°C for *G. ultimum* and in 10% glycerin + 5% trehalose at-80°C for *R. solani*.

for inoculation with *G. ultimum*, three 5 mm-diameter agar plugs from a seven-dayold culture on PDA were transferred to a Petri dish containing 20 g sterilized organic millet seeds soaked in 10 mL autoclaved water. The cultures were incubated in the dark at 20°C for seven days. The colonized millet was then chopped, mixed with sand, and gently homogenized with a mortar. Immediately afterward, 20 g of the millet-sand-pathogen mix, containing either 0.45 g or 1.35 g of inoculum (representing low and high pathogen concentration, respectively), was added per L of compost-substrate or no-compost control. Control pots without pathogen received sand without inoculum. Substrates with and without pathogen were filled into pots with 6 cm (cress) or 9 cm diameter (cucumber). For the *R. solani* bioassays, the pathogen was cultured on PDA for three days. Three agar plugs (diameters: 0.8 or 1.4 cm for sets I–III; 0.6 or 0.8 cm for compost set VI) were placed into pots pre-filled to one-third with substrate, which was then topped up to the brim.

For the cress bioassay, each pot was sown with 0.6 g organic cress seeds (approximately 260 seeds, *Lepidium sativum*, Bigler Samen AG, Steffisburg, Switzerland) and moistened with a hand sprayer. For the cucumber bioassays, seeds (*Cucumis sativus*, cultivar “Chinese snake”, Bigler Samen AG, Steffisberg, Switzerland) were surface-sterilized in 0.1% (v/v) potassium hydrochloride solution for ten min, rinsed six times with sterile water, placed on 1.5% water agar, and incubated at 24°C in the dark for 46-52 hours. Each pot received five pre-germinated seeds, planted at a depth of 2 cm. All pots of a treatment were placed on plastic trays (cress: Licefa, Germany; cucumber: Anzuchtsbox M, Romberg, Germany) and covered for three days to maintain high humidity (cress trays: plastic bags; cucumber trays: transparent lids). After this period, cress pots were watered daily, while cucumber pots were watered every two to four days. Shoot biomass was harvested after 6, 11 and 13 days for the cress-*G. ultimum*, cucumber-*G. ultimum* and cucumber-*R. solani* system, respectively. All bioassays were performed in climate chambers (GroBanks, CLF Plant Climatics, Wertingen, Germany) at a day/night regime of 16 h/8 h and a light intensity of 130 *µ*mol m^-2^ s^-1^. The air humidity was maintained at 70% and the temperatures set to 20°C day/18°C night for *G. ultimum* and 23°C day/21°C night for *R. solani*.

Disease suppression for each substrate was calculated as relative biomass by dividing the shoot weight of plants grown with pathogen application by the mean shoot weight of plants grown without. Additionally, disease symptoms were visually assessed for each cress pot and cucumber plant using a disease scoring. For all three plant-pathogen systems the disease scoring correlated with relative shoot biomass (Figure S2). To compare disease suppression across test sets, one pathogen level was selected based on the relative biomass observed in the no-compost control (Figure S3). We aimed at least 50% reduction in plant biomass due to pathogen inoculation in the no-compost control and no significant difference among the four sets. This was accomplished by applying 0.45 g *G. ultimum*-millet mix L^-1^ substrate for cress-*G. ultimum*; for cucumber-*G. ultimum* 0.45 g L^-1^ was used in sets I and II, and 1.35 g L^-1^ in sets III and IV. For cucumber-*R. solani*, a plug size of 1.4 cm was used in set I, and 0.8 cm in sets II-IV.

### 2.4. Bacterial and fungal amplicon sequencing

For DNA extraction, 200 mg of 2 mm-sieved compost or peat samples were combined with 1 mL of NucleoMag lysis buffer and 1.5 g of beads (SiLibeads Type ZY S 0.4-0.6 mm, Sigmund Lindner, Germany) in 2 mL screw-cap tubes. The samples were prepared in four technical replicates and stored at-20°C until use. DNA was extracted using the “NucleoMag DNA Microbiome for DNA purification from soil, stool and biofilm” DNA extraction kit (Macherey Nagel, Düren, Germany) following the manufacturer’s instructions with some changes. All samples were homogenized using a TissueLyser (Qiagen, Netherlands) at 30 Hz for 4 min. To remove leftover ethanol the magnetic beads, were dried at 70°C for 30 min prior to elution of the DNA in 0.1 x TE buffer (1 mM Tris, 0.1 mM Na_2_EDTA, pH 8.0),. DNA extracts were stored at-20°C. DNA was quantified using Quant-iT PicoGreen dsDNA Assay kit (Invitrogen, Waltham, MA, USA) with a Cary Eclipse fluorescence spectrophotometer (Varian, Palo Alto, CA, USA). For PCR amplification DNA extracts were diluted to 5 ng *µ*L^-1^ using deionized and autoclaved H_2_O.

For the bacteria, the variable regions 3 and 4 of the 16S rRNA gene marker were amplified using the primer pair 341F (5’-CCTAYGGGDBGCWSCAG-3’) and 806R (5’-GGACTACNVGGGTHTCTAAT-3’). For the fungi, the internal transcribed spacer region 2 (ITS2) was amplified using the primer pair ITS3 (5’-CANCGATGAAGAAC GYRG-3’) and ITS4 (5’-CCTSCSCTTANTDATATGC-3’) modified by Tedersoo et al. (2015). PCR reactions were performed as described in Mayerhofer et al. (2021a) with the following modifications: 1 *µ*M of each primer was used, the PCR included 30 and 33 cycles for the 16S and ITS2 primer pair, respectively, and the annealing temperature was set to 56°C. A no-template control was included for each 96-well plate. PCR reactions were run in quadruplets and pooled following quality control on an Agarose gel. DNA concentration of PCR products was quantified as described for genomic DNA and diluted to 20 ng mL^-1^. Library preparation included tagging with TruSeq indices, normalizing read counts from an Illumina MiniSeq run, and sequencing on the Illumina NextSeq2000 platform. Paired-end sequencing was performed with 300 bp long reads (Illumina Inc., San Diego, CA, USA) at the Functional Genomics Center Zurich, University of Zurich / ETH Zurich (Switzerland).

### 2.5. Bioinformatics and quality control

Quality control, ASV inference and taxonomic classification of sequences was performed using a pipeline adapted from Frey et al. (2016) and Mayerhofer et al. (2021b). Briefly, raw sequences were trimmed by removing primer sequences using cutadapt (v.3.5, Martin (2011), one mismatch allowed). Implemented in vsearch (v.2.27.0, Rognes et al. (2016)), sequences were merged, quality filtered and dereplicated. To obtain amplicon sequence variant (ASV), sequences were de-noised with the UNOISE algorithm (Edgar and Flyvbjerg, 2015). Chimera removal was done with UCHIME (Edgar et al., 2011). Checking for the presence of ribosomal signatures was performed using Metaxa2 (v.2.2.3, Bengtsson-Palme et al. (2015)) for the bacteria and ITSx (v.1.1.3, Bengtsson-Palme et al. (2013)) for the fungi. All sequences without ribosomal signature were discarded. Finally, reads were mapped against the final set of identified ASVs using the algorithm “usearch global”. Taxonomic classification of ASVs was performed with the command “clas-sify.seqs” in MOTHUR (v.1.47, Schloss et al. (2009)). Bacterial sequences were classified using the Genome Taxonomy Database (GTDB, release SSU 214; Parks et al. (2021)), trimmed to the 16S gene as described by Mayerhofer et al. (2025, in review). Fungal sequences were classified using the UNITE database (version 9.0; Abarenkov et al. (2024)).

Non-bacterial sequences (e.g., archaea, chloroplasts, and mitochondria) and nonfungal sequences (e.g., protists and plants) were removed. For the fungi three samples were excluded as outliers since they had less than than 10,000 fungal sequences. To standardize read counts, all samples were rarefied to the lowest counts (35,928 bacterial; 10,211 fungal sequences) with 100 iterations using “rrarefy” from the *vegan* package. Analyses were performed on the 100 times sub-sampled ASV tables or their median counts. Number of sequences and ASVs for the different filtering steps are indicated in Table S4.

### 2.6. Statistical analyses

All analyses and visualizations were performed in R (v.4.3.1, R Core Team (2023)) using Rstudio (v.2023.06) and the *tidyverse* packages (Wickham et al., 2019). Statistical significance was set at *p <*0.05, with Benjamini–Hochberg correction applied for multiple testing when appropriate (Benjamini and Hochberg, 1995). Differences in suppressive activity among substrates was analyzed per compost set using one-way analysis of variance (ANOVA) followed by Tukey’s honestly significant difference (Tukey HSD) post hoc test (Lenth, 2024) or Kruskal-Wallis and pairwise Wilcoxon tests when normality (assessed via residual plots) was violated. The same approach was used to compare alpha diversity and other compost properties across substrates, composting sites, and sets. Pearson or Spearman correlation was used depending on Shapiro-Wilk normality tests; only correlations with *|coefficient| >*0.4 were considered significant. Principle Component Analysis (PCA) of compost properties was based on Euclidean distance. Strongly co-correlated compost properties (*|ρ| >* 0.8, Fig. S7) and those with missing data (basal respiration, compost age) were excluded. Microbial community analyses were performed using the *vegan* package (Oksanen et al., 2022). Alpha diversity measures of microbial communities were calculated using the“*specnumber*” and “*diversity*” functions on 100 times subsampled ASV-tables and then averaged. Bray-Curtis dissimilarities were computed using “*vegdist*” on the same sub-sampled data. Compositional differences in physicochemical properties and microbial communities were tested with multivariate permutation analysis of variance (PERMANOVA, Anderson et al. (2006)) using “*adonis2*”, and visualized using nonmetric multidimensional scaling (NMDS) using the “*metaMDS*” function.

Four statistical approaches were used to identify ASV associated with the nine most suppressive composts for each plant-pathogen system: Point Biserial Correlation and Indicator Species Analysis (Cáceres and Legendre, 2009), MaAsLin2 (Mallick et al., 2021), and ALDEx2 (Fernandes et al., 2014). The settings for each approach are provided in Supplementary Methods C. A consensus approach as suggested by Nearing et al. (2022) was applied, considering ASVs as “indicative” if they were significantly associated in at least three approaches. ChatGPT version 4o was used for code debugging.

## 3. Results

### 3.1. The disease-suppressive activity of composts was plant-pathogen-specific

To investigate the factors associated with disease suppression by composts, we evaluated 37 composts across three plant–pathogen systems: cress–*G. ultimum*, cucumber–*G. ultimum*, and cucumber–*R. solani*. Of these, 33 composts were significantly disease suppressive compared to the no-compost control in at least one system (Fig. 1A–C). Three composts (24, 34, 35) were effective in all three systems, while four (9, 10, 11, 12) showed no suppressive effect in any. The highest number of suppressive composts was found in the cress–*G. ultimum* system (Fig. 1D). Fewer composts were effective in the cucumber–*G. ultimum* system, with 14 overlapping between the two *G. ultimum* systems. Suppression of *G. ultimum* in cress and cucumber was significantly correlated (r = 0.63, *p <*0.001; Fig. S4). In cucumber–*R. solani* system, the least composts were suppressive and suppression was not correlated with *G. ultimum*-suppression (cress: r = –0.07, *p* = 0.662; cucumber: r = 0.03, *p* = 0.543).

**Figure 1:**
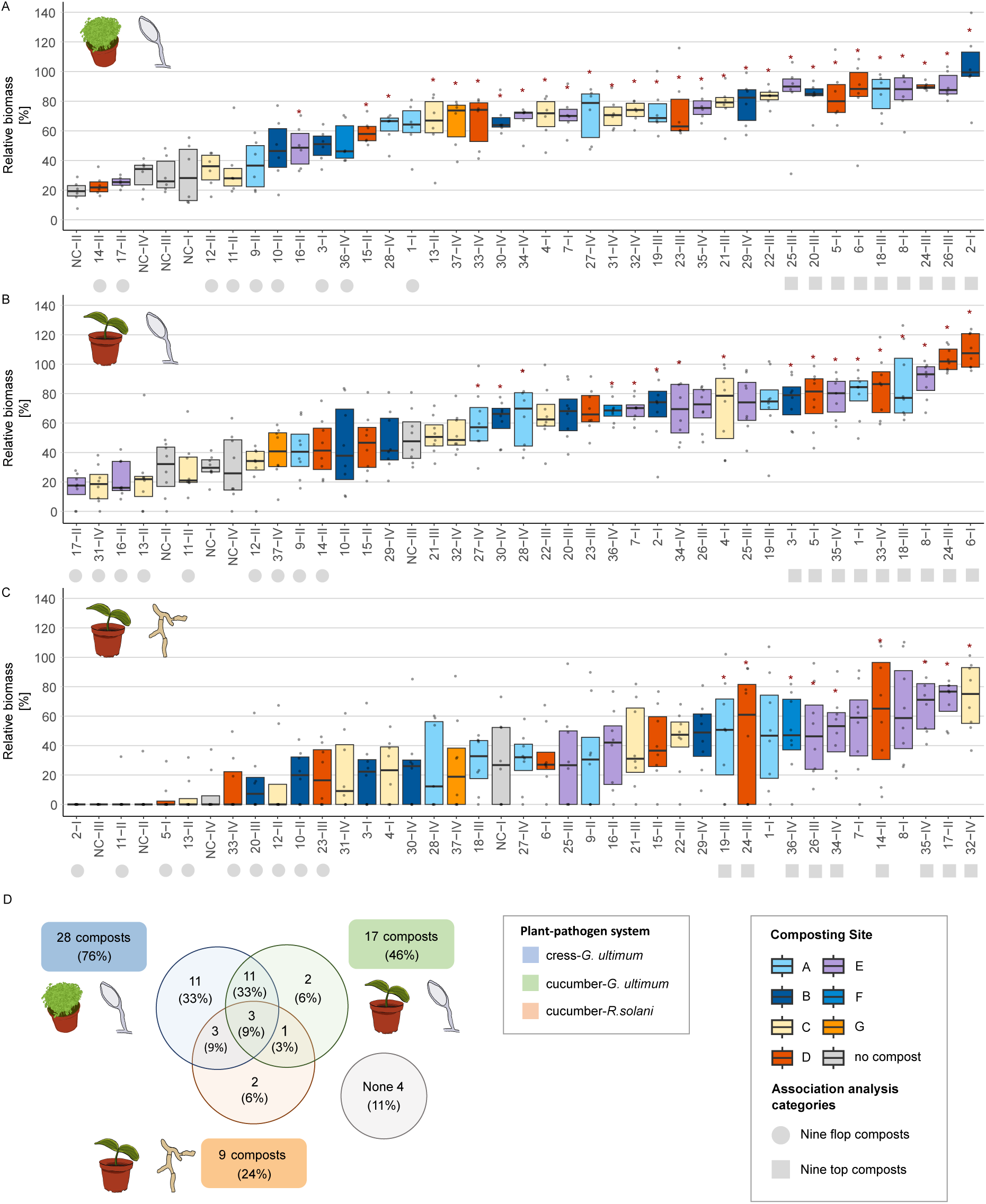
Disease suppression of composts differed among plant-pathogen systems. Thirty-seven composts were tested in three plant-pathogen systems: (A) cress-*Globisporangium ultimum*, (B) cucumber-*G. ultimum* and (C) cucumber-*Rhizoctonia solani*. Disease suppression is expressed as relative biomass, which is calculated by dividing the shoot biomass with pathogen application by the average shoot biomass without pathogen application for each substrate. The relative biomass of each replicate pot (n = 6 for cress and n = 8 for cucumber) is shown as a grey point; boxes represent the interquartile range, and dashes denote the median. Composts are identified by numbers followed by Roman numerals, which represent their respective compost sets (I-IV; I = May 2022, II = July 22, III = September 22, IV = May 2023). No-compost controls are labeled with “NC”. Substrate treatments are ordered by increasing means. Red asterisks indicate compost-substrates with significantly higher relative biomass compared to the no-compost control in the respective set (statistical test results in Table S5). The shape of the symbol following the substrate name highlights the nine most (circle) and least (square) suppressive composts in each system, used for associating single ASVs with the most suppressive composts. (D) The Venn diagram depicts the number of suppressive composts shared across plant-pathogen systems.

The composts originated from seven composting sites (A–G, Table S1), with at least one compost from each site showing suppressive activity in one or more plant-pathogen systems. Suppression did not significantly differ between sites, with two exceptions: composts from site D showed higher suppression in the cucumber–*G. ultimum* system compared to site C, and those from site E were more effective in the cucumber–*R. solani* system than those from site B (Fig. S5). These results indicate that disease suppression cannot be consistently attributed to the composting site.

The 37 composts were collected and tested in four sets of 8-10 composts at different time points. Each set included both suppressive and non-suppressive composts. While the mean suppressive activity of sets I, III and IV was similar, set II showed significantly reduces suppression in both *G. ultimum* systems (Fig. S5).

### 3.2. Physicochemical properties and microbial activities of composts were primarily associated with composting site and compost maturation

Mean disease suppression of the 37 composts in the three plant-pathogens systems showed no correlation with any of the 14 physicochemical compost properties, basal respiration, FDA hydrolysis or compost age (Spearman’s Rank sum test, *|ρ| >* 0.4, *p*-adj *<* 0.05, Table 1, scatter plots Fig. S8, S9, S10). We hypothesized that a combination of properties, rather than individual ones, might be associated with disease suppression. However, a Principle Component Analysis (PCA) based on nine compost properties indicated that this was not the case either (Table 1, Fig. S6). Although not linked to disease suppression, the PCA showed that composts cluster by composting site along the first principle component (PC1; PERMANOVA, R^2^ = 42.4%, Pseudo-F = 3.68, p = 0.001, Fig. 2A), and to a lesser extent by compost age along PC2 (R^2^ = 8.3%, Pseudo-F = 2.99, p = 0.014, Fig. 2B). PC1 was mainly driven by C_org_, FDA, PO_4_^3-^ and salinity, while PC2 reflected variation in pH, NH_4_^+^ and NO_3_^-^—the later typically associated with compost maturation. These patterns suggest that the compost properties reflect composting site and maturation stage rather than disease suppressive activity.

**Figure 2:**
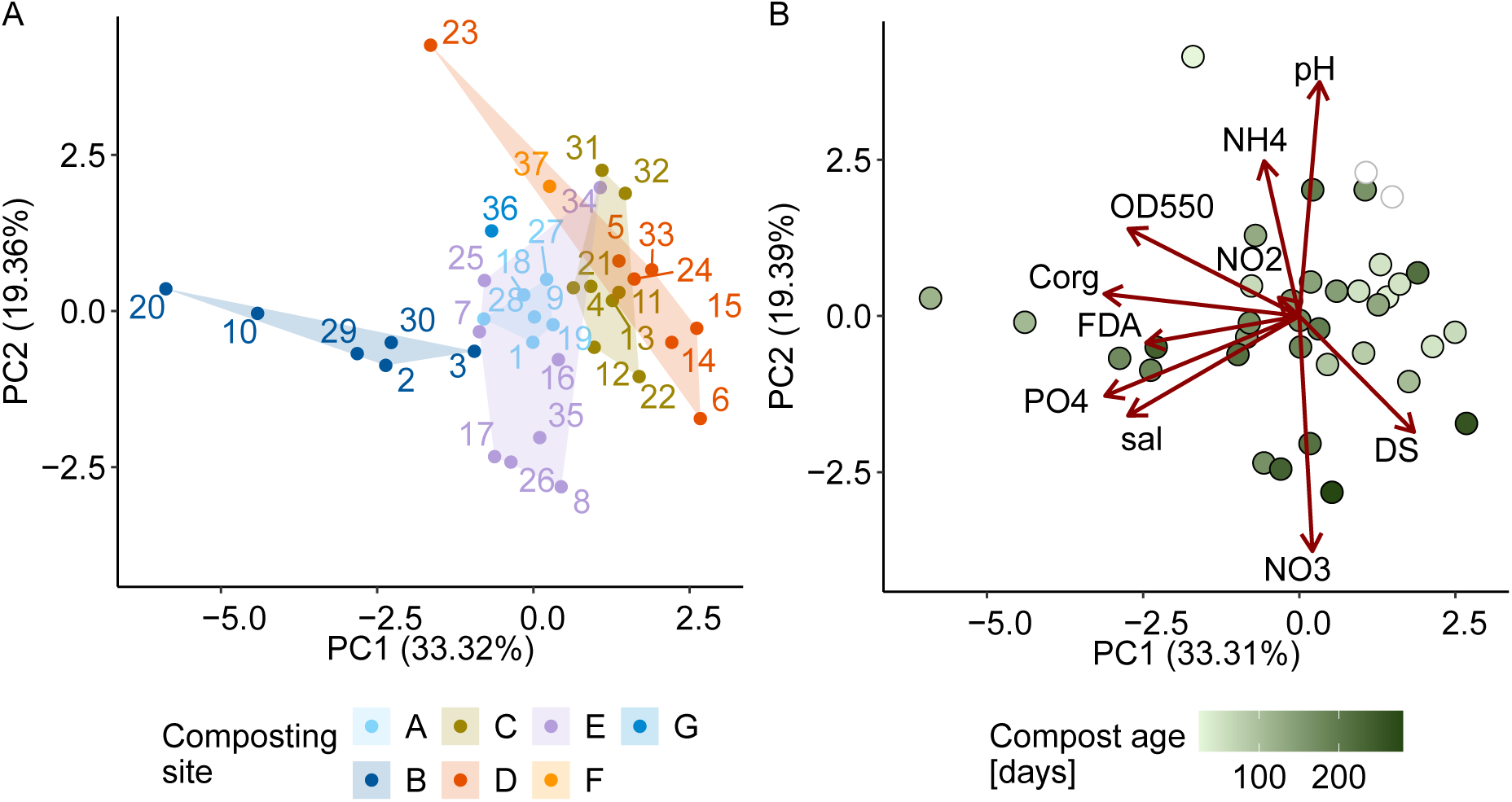
Compost properties were associated with the composting site and compost age. The Euclidean distance between composts was calculated based on nine selected compost properties (indicated with an asterisk in Table 1). The plots display the first and second principal component. Each compost is represented by a dot colored according to either its (A) composting site or (B) composting age. Two composts (31, 32) with unknown compost age are displayed in white. In A, the compost IDs (1-37) are indicated, and in B, the loading vectors of the PCA for all nine compost properties are shown in dark red.

**Table 1:** Correlation analyses of disease-suppressive activity of composts with physicochemical properties, microbial activity, and alpha- and beta-diversity measures. For univariate analysis, Spearman’s Rank sum correlation was used. For multivariate analysis, PERMANOVA was calculated based on Euclidean distance of compost properties and the Bray Curtis dissimilarity of bacterial and fungal ASV abundances. Compost properties included in the PCA in Figure 2 are marked with an asterisk. *P*-values for Spearman’s correlation were adjusted for multiple testing using the Benjamini-Hochberg correction for compost properties and microbial activities (n = 17) and alpha diversity measures (n = 3).

|  |  | cress-*G. ultimum* |  |  | cucumber-*G. ultimum* |  |  | cucumber-*R. solani* |  |  |
| --- | --- | --- | --- | --- | --- | --- | --- | --- | --- | --- |
| | | $\rho$ | *p* | *p*-adj | $\rho$ | *p* | *p*-adj | $\rho$ | *p* | *p*-adj |
| **Compost properties** | Compost age [days] | 0.21 | ns | ns | 0.37 | * | ns | 0.18 | ns | ns |
|  | DS* [%] | -0.08 | ns | ns | 0.04 | ns | ns | 0.32 | . | ns |
|  | Max. WHC<sup>a</sup> | 0.01 | ns | ns | -0.18 | ns | ns | -0.27 | . | ns |
|  | pH* | -0.03 | ns | ns | -0.20 | ns | ns | -0.17 | ns | ns |
|  | Salinity*<sup>b</sup> | 0.11 | ns | ns | 0.14 | ns | ns | 0.11 | ns | ns |
|  | OD<sub>550</sub>* | 0.12 | ns | ns | 0.05 | ns | ns | -0.19 | ns | ns |
|  | NO<sub>2</sub><sup>-</sup>-N<sup>c</sup> | -0.28 | . | ns | -0.24 | ns | ns | 0.14 | ns | ns |
|  | NO<sub>3</sub><sup>-</sup>-N*<sup>c</sup> | 0.10 | ns | ns | 0.12 | ns | ns | 0.30 | . | ns |
|  | NH<sub>4</sub><sup>+</sup>-N*<sup>c</sup> | -0.22 | ns | ns | -0.30 | . | ns | -0.10 | ns | ns |
|  | N<sub>min</sub><sup>c</sup> | 0.06 | ns | ns | 0.06 | ns | ns | 0.23 | ns | ns |
|  | NO<sub>3</sub><sup>-</sup>-N/N<sub>min</sub> | 0.20 | ns | ns | 0.37 | * | ns | 0.27 | ns | ns |
|  | PO<sub>4</sub><sup>3-</sup>-N*<sup>c</sup> | 0.09 | ns | ns | 0.06 | ns | ns | 0.05 | ns | ns |
|  | N<sub>tot</sub> [%] | 0.11 | ns | ns | 0.06 | ns | ns | 0.06 | ns | ns |
|  | C<sub>org</sub>* [%] | -0.03 | ns | ns | -0.07 | ns | ns | -0.10 | ns | ns |
|  | C<sub>org</sub>/N<sub>tot</sub> | -0.27 | ns | ns | -0.26 | ns | ns | -0.38 | * | ns |
| **Microbial activity** | Basal respiration<sup>d</sup> | 0.05 | ns | ns | 0.04 | ns | ns | -0.30 | . | ns |
|  | FDA hydrolysis*<sup>e</sup> | -0.16 | ns | ns | -0.45 | ** | ns | -0.01 | ns | ns |
| **Alpha diversity** | Bacterial richness | 0.49 | ** | ** | 0.31 | . | ns | 0.29 | . | ns |
|  | Bacterial Shannon evenness | 0.45 | ** | ** | 0.16 | ns | ns | 0.04 | ns | ns |
|  | Bacterial Shannon diversity | 0.55 | *** | ** | 0.24 | ns | ns | 0.18 | ns | ns |
|  | Fungal richness | 0.25 | ns | ns | 0.13 | ns | ns | 0.14 | ns | ns |
|  | Fungal Shannon evenness | 0.35 | * | . | 0.18 | ns | ns | 0.14 | ns | ns |
|  | Fungal Shannon diversity | 0.36 | * | . | 0.16 | ns | ns | 0.16 | ns | ns |
|  |  | F (R<sup>2</sup>) | *p* | *p*-adj | F (R<sup>2</sup>) | *p* | *p*-adj | F (R<sup>2</sup>) | *p* | *p*-adj |
| **Multivariate analysis** | PCA compost properties | 0.82 (2.3%) | ns | - | 1.01 (2.8%) | ns | - | 1.54 (4.0%) | ns | - |
|  | Bacterial community structure | 1.48 (4.0%) | ns | - | 0.92 (2.6%) | ns | - | 1.38 (3.8%) | ns | - |
|  | Fungal community structure | 1.36 (3.9%) | ns | - | 1.10 (3.1%) | ns | - | 1.16 (3.2%) | ns | - |

ns = not significant, "." *p* < 0.1, "\*" *p* < 0.05, "\*\*" *p* < 0.01, "\*\*\*" *p* < 0.001,
<sup>a</sup> [g H<sub>2</sub>O (g DS)<sup>-1</sup>], <sup>b</sup> [g KCl<sub>eq</sub> (kg DS)<sup>-1</sup>], KCl<sub>eq</sub> = Potassium chloride-equivalent, <sup>c</sup> [mg (kg DS)<sup>-1</sup>], <sup>d</sup> [mg CO<sub>2</sub>-C h<sup>-1</sup>], <sup>e</sup> [μg FDA (min g DS)<sup>-1</sup>]

### 3.3. Bacterial and fungal communities were linked with compost properties but not with disease-suppressive activity

We characterized the bacterial and fungal communities of the 37 composts and four no-compost controls using 16S (V3-V4) and ITS2 amplicon sequencing, identifying 19,129 bacterial and 2,309 fungal ASVs. Non-metric multidimensional scaling (NMDS) ordination revealed tight clustering of technical replicates (Fig. S11) and PERMANOVA confirmed strong differences among composts in both bacterial (R^2^ = 96.1%, Pseudo-F = 76.01, p = 0.001) and fungal community structure (R^2^ = 96.2% Pseudo-F = 76.14, p = 0.001). Only 0.8% of bacterial and 0.3% of fungal ASVs were shared across all composts, further emphasizing community distinctness across composts. Despite low ASV overlap, abundant taxa at the phylum, family, and genus level were common across composts (Fig. S12 and S13, Table S6). Bacterial communities were taxonomically diverse, while fungal communities were often dominated by a few genera. Bacterial and fungal community structure were significantly correlated (Mantel test, Pearson’s *r* = 0.62, *p* = 0.001).

All assessed alpha diversity measures (ASV richness, Shannon evenness, and Shannon diversity) differed significantly across composts for both bacteria and fungi (Fig. 3, Fig. S14, ANOVA results in Table S7). Bacterial richness ranged from 955 to 5,021 ASVs (mean: 2,691, SD: 936), and fungal richness from 35 to 401 ASVs (mean: 177.5, SD: 96.9). Bacterial and fungal alpha diversity measures were significantly correlated (*ρ* = 0.66-0.74, p *<*0.001).

**Figure 3:**
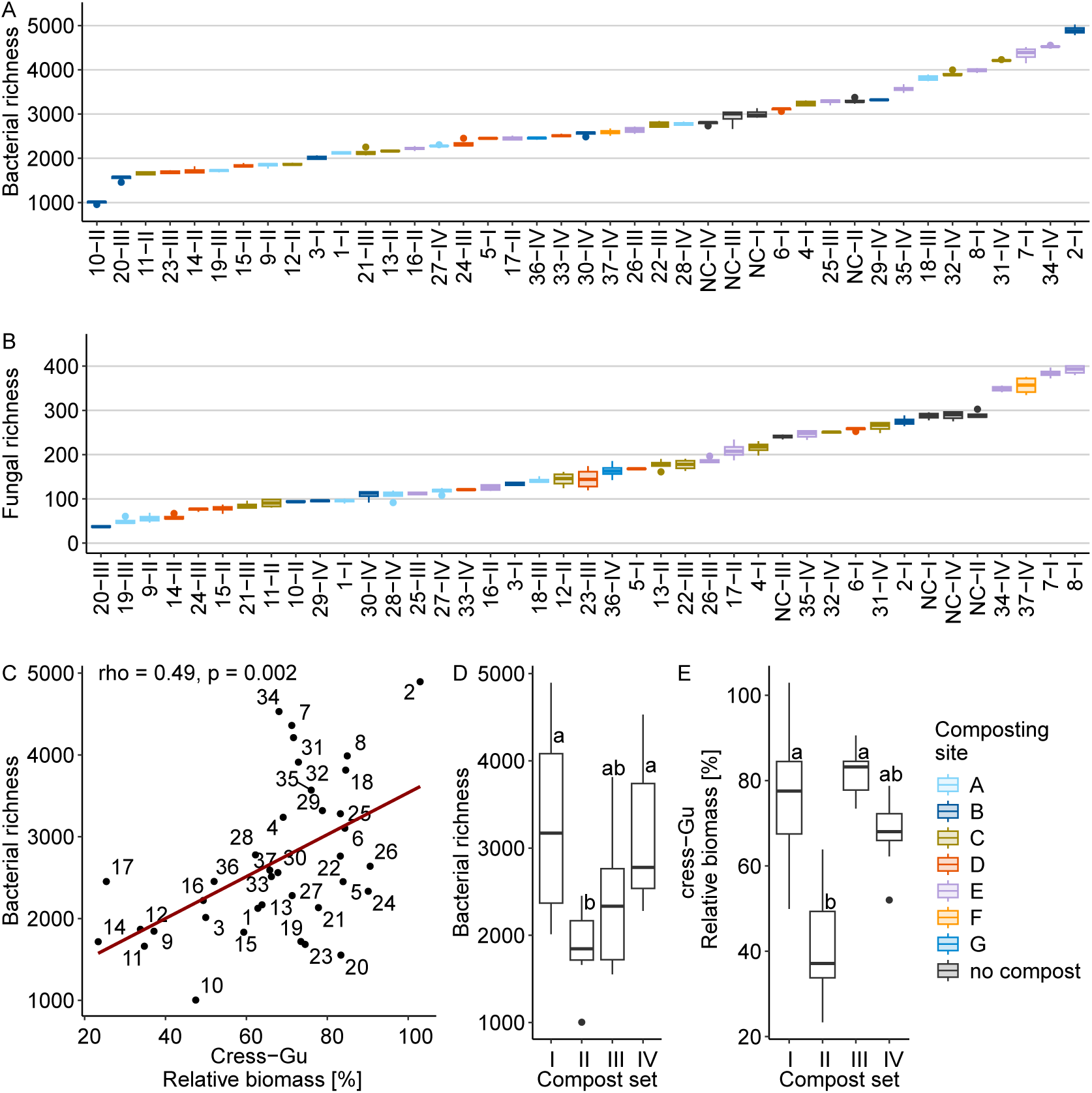
**Bacterial and fungal diversity differed among composts and sets but did not explain differences in disease suppression**. Boxplots display (A) bacterial and (B) fungal richness for the 37 composts (1-37) and the no-compost control (NC) of the four compost sets (I-IV) ordered by their means (n =4). (C) The scatter plot shows the relationship between bacterial richness and disease suppression in the cress-*G. ultimum* system. The linear trend line is depicted in dark red. Boxplots show differences among compost sets for (D) bacterial richness and (E) disease suppression in the cress-*G. ultimum* system. For all boxplots the boxes represent the interquartile range (IQR, 25^th^ to 75^th^ percentile), the horizontal line within the box indicates the median, the whiskers extend to 1.5 x IQR, and outliers (*>*1.5 x IQR) are shown as points.

All bacterial alpha diversity measures were positively correlated with *G. ultimum* suppression in cress, while fungal evenness and diversity showed a non-significant positive trend (Fig. 3C, Table 1). No such relationships were found for the cucumber systems. This correlation in cress was driven by compost from set II, which had low diversity and low suppressive activity. In contrast, set III showed high suppression despite low diversity (Fig. 3D and E). Removing set II from the analysis eliminated the observed correlations (p *>*0.05).

To explore potential drivers of alpha diversity, we assessed correlations with compost properties (Table S8). Bacterial richness increased with compost age and the nitrate-N/N_min_—both associated with compost maturation—while fungal richness showed similar, albeit not significant, trends.

Bacterial and fungal community structure, based on the mean ASV abundance of technical replicates, was not significantly associated with disease-suppressive activity in any of the three plant-pathogen systems (PERMANOVA, Table 1, Fig. S15). However, both bacterial and fungal communities were significantly associated with the composting site (Fig. 4A and B, Table S9), the compost age (Fig. 4C and D) and with most compost properties (Table S8). Bacterial community structures of the composts was most strongly associated with OD_550_, PO_4_^3-^ and FDA hydrolysis, while fungal community structure with N_tot_, salinity and NO_3_^-^. In summary, composting site and compost properties explained the fungal and bacterial community structure, but this was not linked to disease suppression.

**Figure 4:**
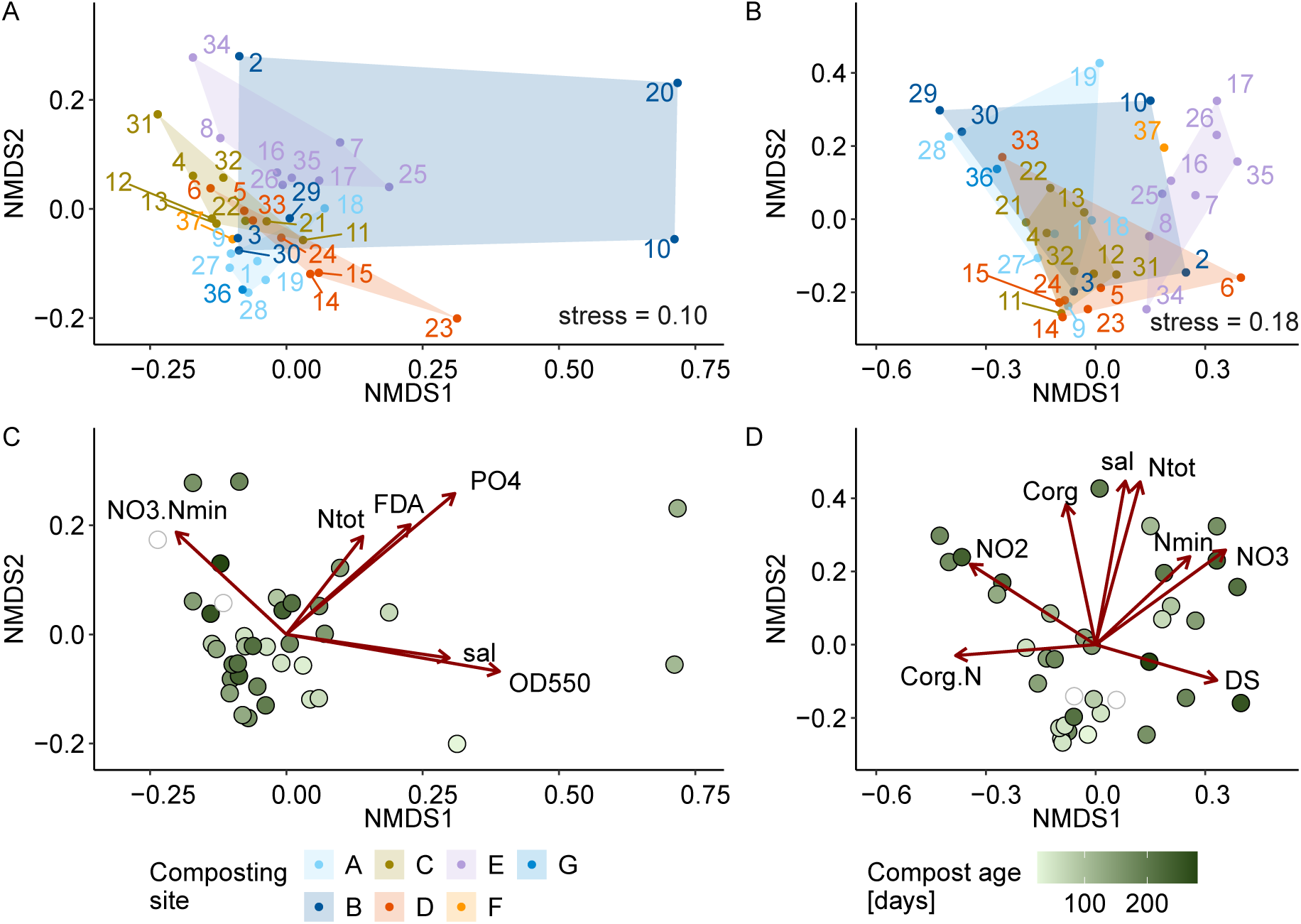
Bacterial and fungal community structure were linked to composting site, compost age, and compost properties. Bray-Curtis dissimilarities were calculated based on the mean ASV abundance of the four replicates. A NMDS plot is shown for (A, C) bacterial community structure and (B, D) fungal community structure colored by the (A, B) composting site and (C, D) composting age. The dark red arrows indicate compost properties significantly associated with community structure along the first two NMDS axes (PERMANOVA results are provided in Tables S8 and S9). Compost 20, which exhibited a distinct community structure, was considered an outlier and excluded from the fungal ordination (Fig. S11).

### 3.4. Potential indicator taxa for disease-suppressive composts were mostly bacteria and specific to the plant-pathogen systems

Bacterial and fungal community structures did not explain differences in disease suppression. However, specific taxa may still be associated with suppressive composts. To investigate this, we compared the abundance of individual bacterial and fungal ASVs between the nine most and nine least suppressive composts for each plant-pathogen system (selected composts shown in Fig. 1A–C). ASVs were included in the association analysis if their mean abundance was at least one read in six or more of the 18 selected composts. This was the case for 70, 85 and 88 fungal and 1,738, 1,827 and 1,664 bacterial ASVs, for the cress-*G. ultimum*, cucumber-*G. ultimum* and cucumber-*R. solani* system, respectively. ASVs were considered indicative if they were statistically significant in at least three of the four methods (Point Biserial Correlation, Indicator Species Analysis, MaAsLin2, ALDEx2). In total, 317 bacterial and 9 fungal ASVs were identified as indicative of the most suppressive composts in at least one system (number of ASVs for each method and overlaps shown in Fig. S16).

All indicative fungal ASVs were specific to individual plant-pathogen systems (Fig. 5A). Those associated with *G. ultimum*-suppression in cress and *R. solani*-suppression in cucumber belonged to the phylum *Ascomycota* but could not be classified at the genus level (Table S15). In the cucumber-*G. ultimum* system, three ASVs were identified as *Ascomycota*, including one as *Kernia* sp. and one as *Scedosporium* sp. Additionally, one ASV was classified to the phylum *Basidiomycota* (genus *Coperinellus)*, and one to the phylum *Mortierollomycota* (genus *Mortierella*). Compared to bacteria, fewer fungal ASVs were identified, and their association with disease suppression was weaker, as indicated by lower effect sizes in statistical tests (Table S15). Consequently, bacterial taxa appear more relevant for indicating disease suppression by composts, and further analyses focused on them.

**Figure 5:**
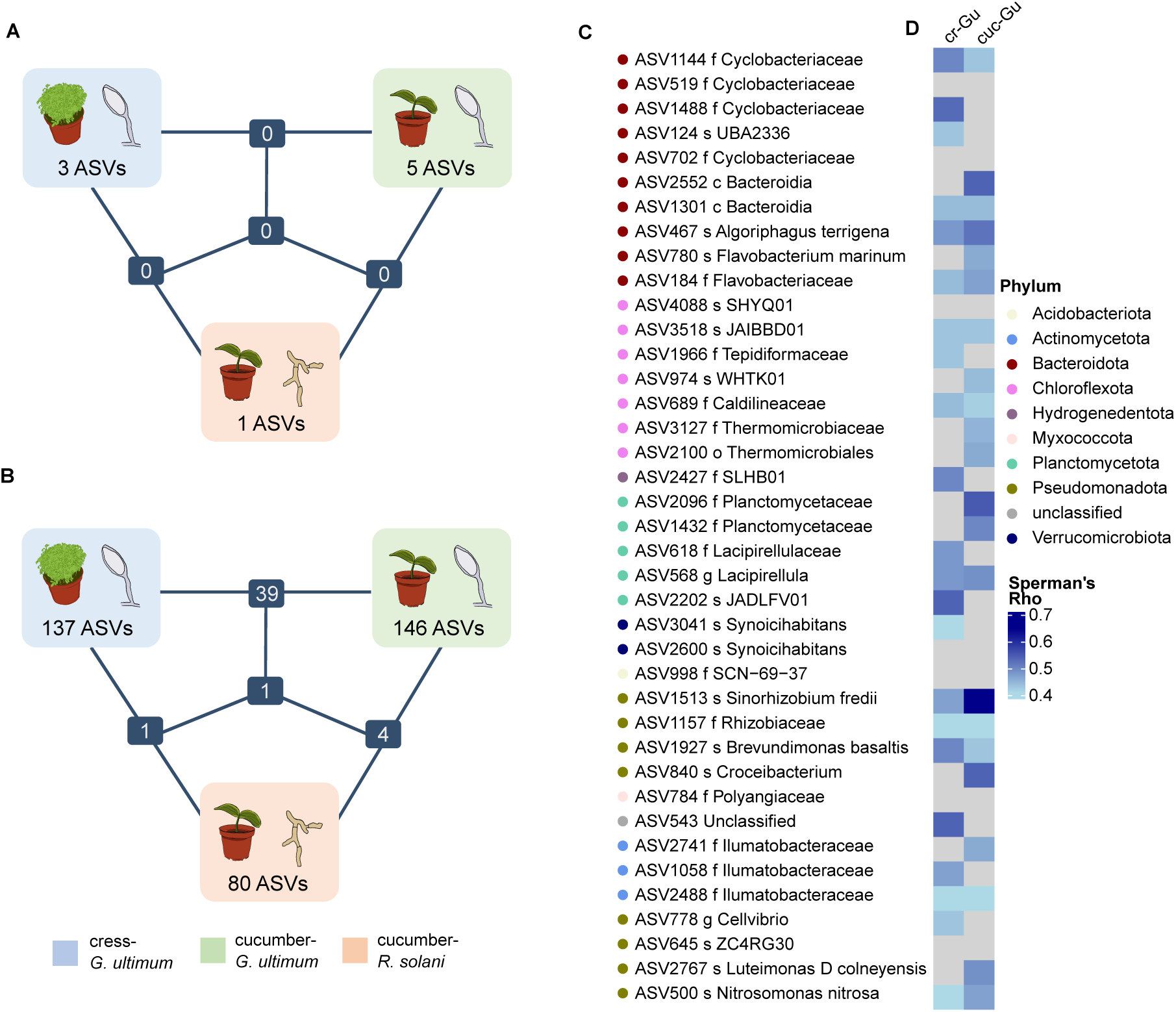
ASVs indicative of the most suppressive composts were predominantly bacteria and specific to a plant-pathogen system. Number of indicative ASVs in the three plant-pathogen systems, cress-*G. ultimum*, cucumber-*G. ultimum* and cucumber-*R. solani*, and the number shared among the systems are shown for (A) fungi and (B) bacteria. (C) Depicts the 40 indicative bacterial ASVs shared between the two *G. ultimum* systems. For all ASVs the lowest taxonomic classification based on the Genome Taxonomy Database (GTDB) is indicated (s = species, g = genus, f = family, o = order, c = class). (D) A heat map shows the correlation between ASV abundance and the diseasesuppressive activity against *G. ultimum* in cress and cucumber across all 37 composts. Spearman Rank sum correlation was used and *p*-values were adjusted for multiple testing (Benjamini & Hochberg). Nonsignificant correlations (*p*-adj *<*0.05) are indicated as gray boxes. cr-Gu: cress-*G. ultimum*, cuc-Gu: cucumber-*G. ultimum*

Most bacterial ASVs were system-specific, with more ASVs associated with *G. ultimum*-suppression in cress and cucumber than *R. solani*-suppression in cucumber (Fig. 5B, all indicative ASVs shown in S16, S17 and S18). One bacterial ASV (ASV2887, order Longimicrobiales) was indicative across all three systems. Indicative bacterial ASVs were taxonomically diverse, spanning 18 phyla. In all three systems, the most common phylum was *Pseudomonadota* (Table S10), which was also the most abundant phylum overall (Table S6). While over two thirds of the indicative ASVs were assigned to family level, genus level assignments were less frequent— 23%, 37% and 35% in the cress-*G. ultimum* system, cucumber-*G. ultimum* system and cucumber-*R. solani* system, respectively. Taxonomic overlap at the genus level was low, with few ASVs sharing the same classification (Table S10).

To identify the strongest bacterial indicators, ASVs were ranked by effect size across all statistical methods, and rank sums were used to identify the top 20 per system (Table 2). While most indicators were system-specific, 40 ASVs were shared between the two *G. ultimum* systems (Fig. 5B). These shared ASVs may serve as general indicators of *G. ultimum*-suppression and are shown in Table 5.

**Table 2:**
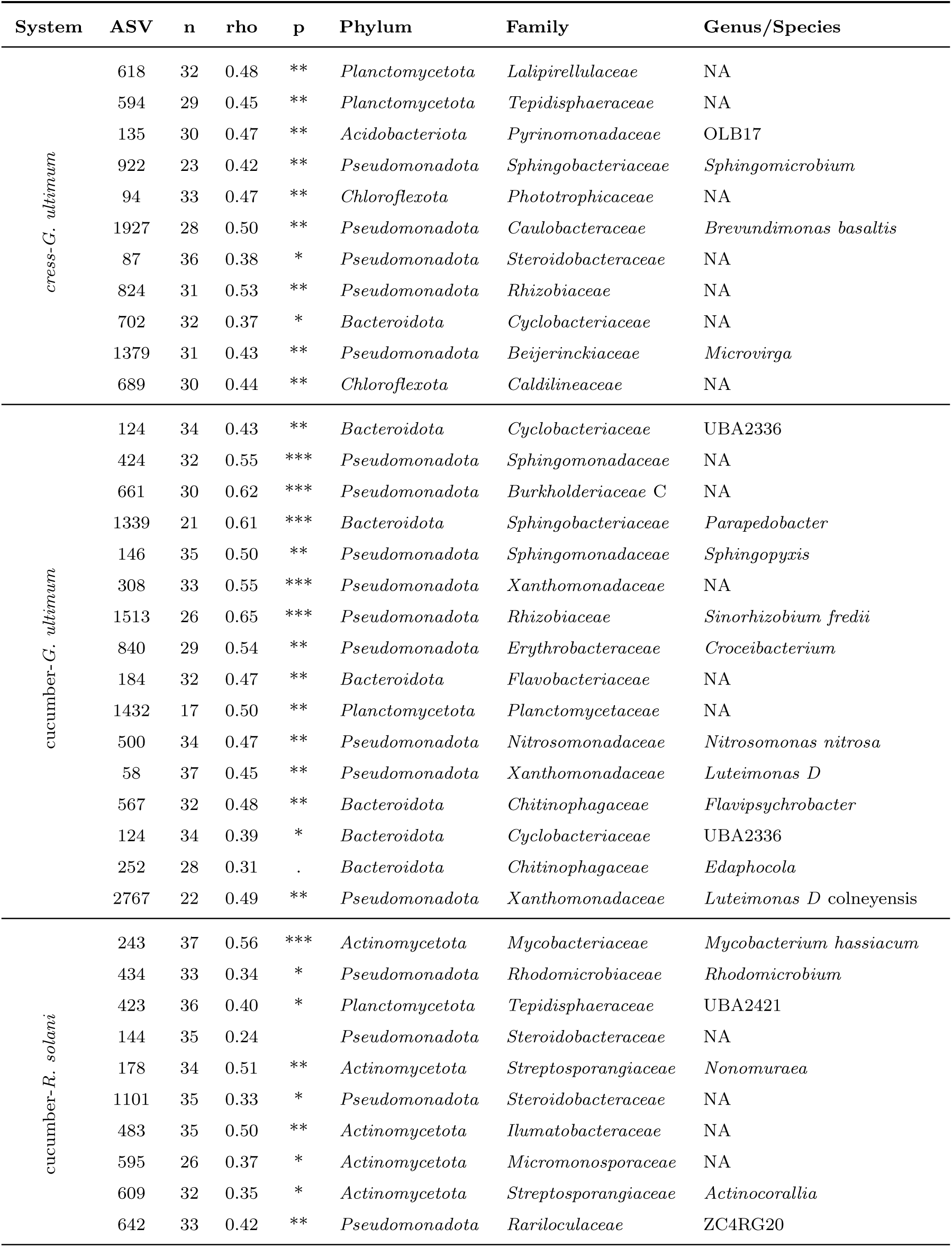
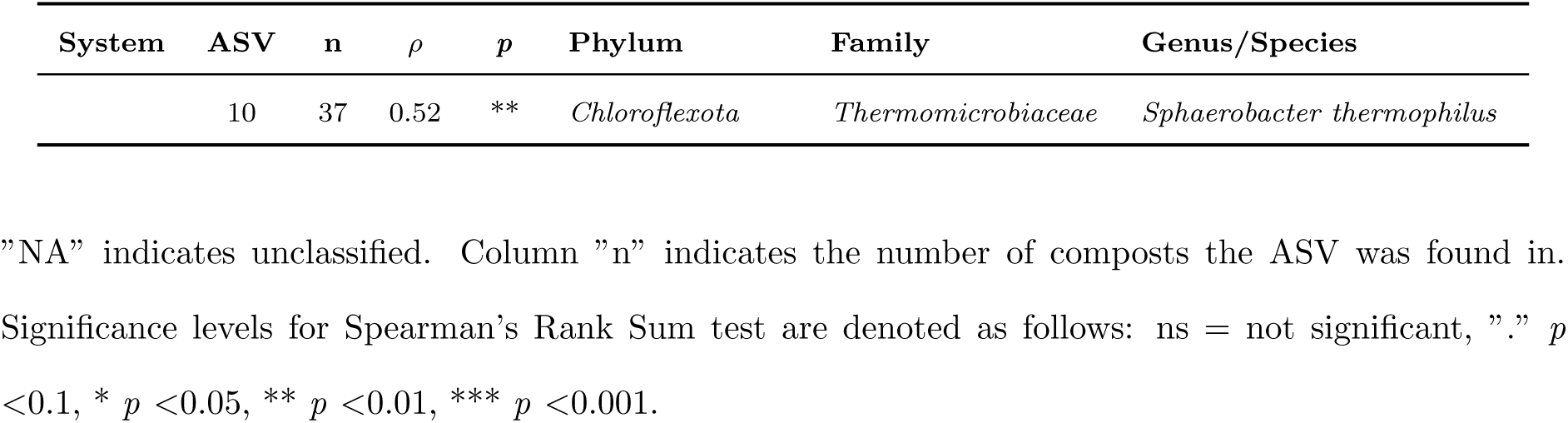
Bacterial ASVs most strongly associated with suppressive composts and their correlation with disease suppression across all 37 composts for each plant-pathogen system. Strength of association with nine most suppressive composts was determined based on the summed rank of the effect size of all four statistical methods (Point Biserial correlation, Indicator Species Analysis, M aAsLin2, ALDEx2; rank sum shown in Tables S16, S17 and S18). Only ASVs which were among the top 20 and classified at least to family level are shown.

We conducted a literature search for all 62 bacterial and 4 fungal genera assigned to indicative ASVs. Forty-four bacterial and all fungal genera have previously been reported to occur in composts (Table S13). For 28 bacterial and one fungal genus, prior evidence supported their involvement in disease suppression (Table S14).

To assess consistency with previous work, we compared the 317 ASVs from this study with 75 indicative ASVs from Mayerhofer et al. (2021a) with a BLAST search. Twelve matched at *>*99% identity to an ASV from the previous study: six were specific to cress–*G. ultimum*, four to cucumber–*G. ultimum*, one shared between *G. ultimum* systems, and one from the cucumber–*R. solani* system (Table S11). Notable matches included ASV467 (*Algoriphagus*, 99.8% identity) and ASV146 (Sphingopyxis, 100%), both among the top 20 indicators for *G. ultimum* suppression (Table 2).

### 3.5. The abundance of indicative bacterial ASVs reflects the suppression patterns across all 37 composts

The indicative bacterial ASVs were identified by comparing the nine most and nine least suppressive composts for each plant-pathogen system. To evaluate their relevance across all composts, we assessed their presence and abundance. Most of the 317 ASVs were detected in more than two-thirds of the 37 composts and had higher median relative abundance in significantly suppressive than in non-suppressive composts (0.016% vs. 0.004%) (Fig. S17). Generally, they showed intermediate abundance (0.01–0.1%) in suppressive composts and low abundance (*<*0.01%) in non-suppressive ones. A substantial proportion of the indicative ASVs correlated significantly and positively with suppression in their respective system: 80 in cress-*G. ultimum* (60.6%), 76 in cucumber-*G. ultimum* system (52.1%), and 45 in cucumber-*R. solani* system (56.3%)(*|ρ| >*0.4, p *<*0.05; Table S16, S17, S18).

To evaluate whether the indicative ASVs reflect the suppressive activity of individual composts, we summed their scaled abundances per compost for each plant-pathogen system. As expected, summed ASV abundances were lowest in the least suppressive composts and highest in the most suppressive ones (”flop v. “top” categories; Fig. 6). Notably, also composts with intermediate suppression levels, which were not included in the initial association analysis, showed increasing summed ASV abundance with increasing suppression. Across all 37 composts, summed ASV abundance was significantly correlated with disease suppression in each system (cress-*G. ultimum*: *ρ* = 0.61; cucumber-*G. ultimum*: *ρ* = 0.62; cucumber-*R. solani*: *ρ* = 0.71; all p *<*0.001).

**Figure 6:**
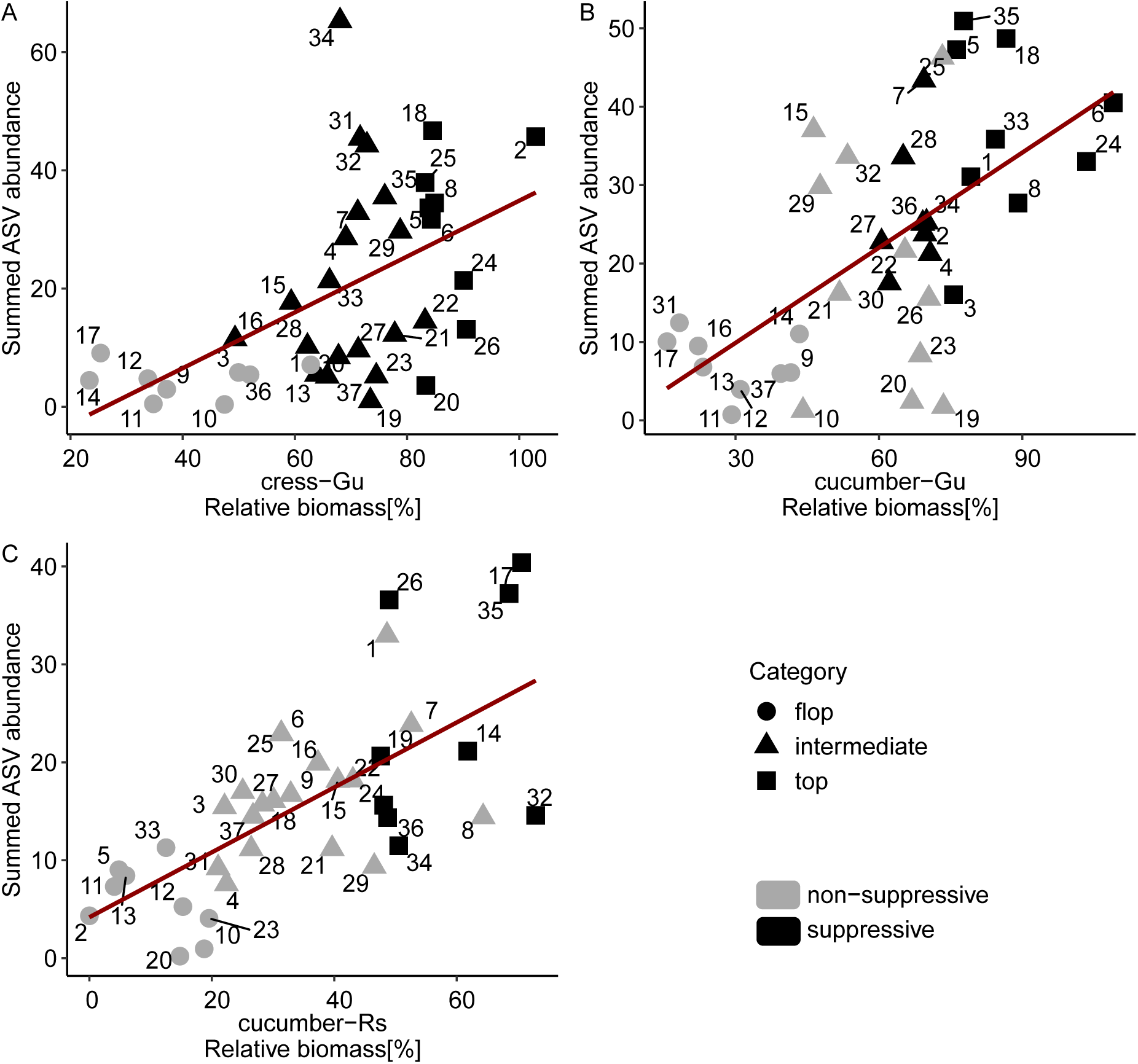
Summed abundance of indicative ASVs was correlated with disease suppression across all 37 composts for (A) the cress-*G. ultimum* system (137 ASVs), (B) the cucumber-*G. ultimum* system (147 ASVs) and (C) the cucumber-*R. solani* system (80 ASVs). To ensure equal weighting of each ASV, its abundance was scaled to a range of 0 to 1 prior to summation. The shape of the points represents the suppression category, with “top” and “flop” used to identify indicative ASVs during the association analysis. Shading indicates whether a compost was significantly suppressive compared to the no-compost control (Fig. 1). The linear trend lines are shown in dark red.

Summed ASV abundance best represented suppression in the cucumber-*R. solani* system (Fig 6C). In the cress–*G. ultimum* system, several significantly suppressive composts—most not among the top nine—showed similarly low summed ASV abundance as non-suppressive composts (Fig.6A). Conversely, some non-suppressive composts in the cucumber–*G. ultimum* system (composts 15, 25, 29, and 32) exhibited high summed abundance of indicative ASVs (Fig. 6B). These observations indicate that although the summed abundance of indicative ASV does not allow reliable prediction of the suppressive effect of all composts, the overall trends are well represented, even for composts not included in the association analysis.

## 4. Discussion

### 4.1. Suppressive activity of compost was plant-pathogen-specific and not linked to compost properties or microbial communities

In this study, most of the 37 composts suppressed disease in at least one plant–pathogen system (Fig. 1). Composts with no suppressive activity were rare, as were those with broad-spectrum efficacy. Suppression of *G. ultimum* was more frequent than that of *R. solani*, which is consistent with previous studies (Scheuerell and Mahaffee, 2005; Termorshuizen et al., 2006; Bonanomi et al., 2018; De Corato et al., 2019). However, suppression efficacy against *G. ultimum* also varied between the two host plants, cress and cucumber. These differences might be explained by plant-specific recruitment of beneficial microorganisms. Root exudates are known to shape rhizosphere microbiota (Mendes et al., 2011) and vary across plant species (Matthews et al., 2019). Plant-induced assembly of compost-associated microorganisms has been proposed as a mechanism underlying disease suppression (De Corato, 2020) and has been experimentally demonstrated in compost-mediated suppression of vascular wilt in tomato (Antoniou et al., 2017; Ding et al., 2023). Together, these findings highlight that indicators for suppressive composts need to be defined specifically for different plant-pathogen systems.

The two pathogens, *G. ultimum* and *R. solani*, prefer different environmental conditions and are primarily controlled by different mechanisms (Lamichhane et al., 2017).

*G. ultimum* is often reported to be suppressed by general microbial activity or diversity, as it competes poorly with saprophytic microorganisms in nutrient-rich environments (Scheuerell and Mahaffee, 2005). *R. solani* is suggested to be controlled by specific biocontrol organisms (Hoitink and Boehm, 1999). However, in this study, neither microbial activity nor diversity could explain suppression of *G. ultimum*. Although, disease suppression in the *G. ultimum*-cress system was positively correlated with bacterial diversity, further analysis revealed that this correlation was driven by composts from set two, which showed both low suppressive activity and low bacterial diversity. In contrast, composts from set III exhibited low bacterial diversity yet were among the most suppressive. This suggests that high bacterial diversity (relative to typical compost diversity levels) is not a prerequisite for effective *G. ultimum* suppression. This observation aligns with our previous findings in cress, as well as with studies on *G. ultimum* in cucumber and *Phy-tophthora nicotianae* in pepper, which reported similar conclusions (Mayerhofer et al., 2021a; Yu et al., 2015; Blaya et al., 2016).

The low suppressiveness of composts from set II was likely linked to the spring drought in 2022, during which they were composted (MeteoSchweiz, 2022). We suspect that this environmental stress may have interfered with the development of disease-suppressive activity against *G. ultimum*, which cannot only be explained by low microbial diversity, but is probably due to the lack of specific beneficial microorganisms. Interestingly, suppressiveness against *R. solani* appeared unaffected by the dry conditions, with composts 14 and 17 from set II being among the most effective against this pathogen.

Our analysis showed that, across all three plant–pathogen systems, compost-mediated disease suppression was not linked to the physicochemical properties of the compost, which is consistent with the conclusions of a large meta-analysis on indicators of disease-suppressive amendments (Bonanomi et al., 2010). This finding extended to fungal and bacterial community structures and alpha diversity, none of which were associated with suppression either. However, physicochemical properties and bacterial and fungal community structure were strongly associated with the composting site and were interrelated (Fig. 2 and Fig. 4). As composting sites differ in many respects, e.g. in terms of feedstock composition, composting method, local microbiota, and environmental conditions, we could not determine the main factors influencing community structures. Some studies suggest that feedstock composition strongly influences physicochemical properties (reviewed by Kong et al., 2024) and microbial community development, particularly during compost maturation (Villar et al., 2016; Estrella-González et al., 2020). In contrast, a study on 116 composts found that pH, moisture content, and total nitrogen—rather than feedstock—best explained variation in bacterial communities (Wang et al., 2020). Thus, it remains unclear whether compost properties or composting site-specific factors primarily shape the microbial communities in composts.

Bacterial and fungal diversity tended to increase as the compost matured, a trend that has already been reported for bacteria (Estrella-Gonźalez et al., 2020; HernándezLara et al., 2022). The compost maturation stage has been linked to suppressive activity against *G. ultimum* in earlier studies (Scheuerell and Mahaffee, 2005; Neher et al., 2022). Also in our previous work, we found that younger composts were more effective at suppressing *G. ultimum* than older, more mature ones (Mayerhofer et al., 2021a). However, in the larger compost set analyzed here, both young, microbial active composts (e.g., composts 5 and 25) and older, less active ones (e.g., composts 6 and 8) were able to strongly suppress *G. ultimum* in cress. Further analysis of the 17 composts from our earlier study (Mayerhofer et al., 2021a) showed that when disease suppression and compost age were tested together, only compost age had a significant association with bacterial community structure (Table S12). This suggests that the previously observed link between community structure and disease suppression is probably indirectly due to compost age.

In summary, our findings show that disease suppression by composts is not explained by physicochemical properties, overall microbial diversity, or microbial community structure, although it is likely influenced by both abiotic and biotic factors (Hadar and Papadopoulou, 2012). This emphasizes the need for more targeted analyses, particularly of specific microbial taxa, which will be discussed in the following section. Figure 7 summarizes our findings and outlines the complex relationship between disease-suppressive activity and compost characteristics.

**Figure 7:**
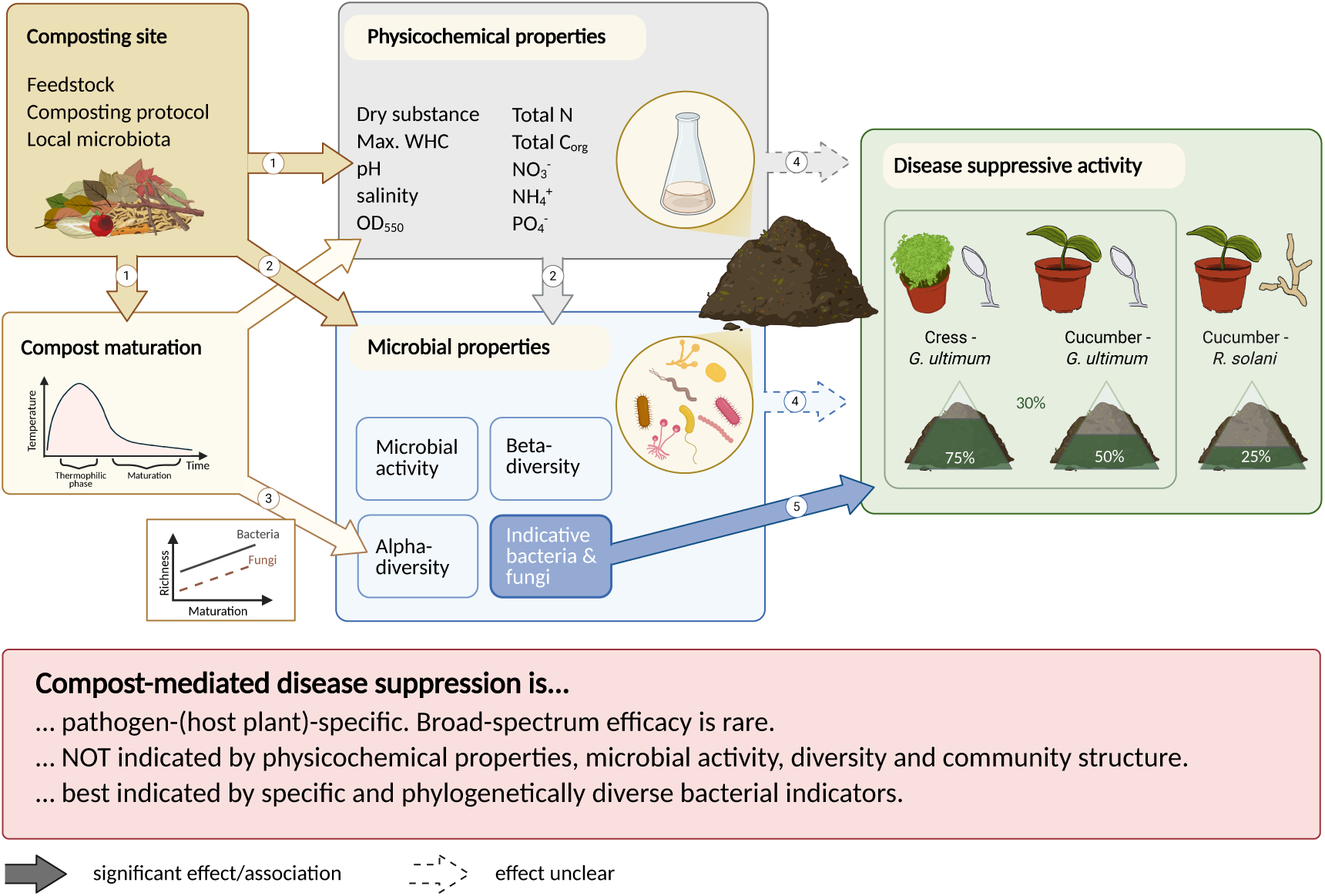
The complex relationship between disease-suppressive activity and compost characteristics. (1) Composting site-specific factors (particularly feedstock) are known to influence compost maturation and physicochemical properties (Estrella-Gonźalez et al., 2020; Bai et al., 2024) (2) Composting site, physicochemical compost properties and maturation processes influence the microbial compost properties. (3) Bacterial and fungal richness increase with compost maturation. Compost microbial communities are shaped by compost properties, composting site and maturation processes. (3) Bacterial and fungal richness increases with compost maturation. (4) The physicochemical properties, microbial activity, and alpha and beta diversity may influence the disease suppressive activity, but were not indicative for suppression in our study. (5) Association analyses between the most and least suppressive composts for each plant-pathogen system revealed indicative taxa for disease suppression. Created in https://BioRender.com

### 4.2. Bacterial ASVs comprised promising indicators for disease-suppressive composts

By comparing the nine most and least suppressive composts, we identified bacterial and a few fungal ASVs indicative of disease suppression for each plant-pathogen system. Most bacterial and all fungal ASVs were plant-pathogen specific, with the greatest overlap observed between the two *G. ultimum* systems (Figure 5). We focused on the bacterial indicators and demonstrated that the indicative bacteria were phylogenetic diverse and generally prevalent in composts. For each plant-pathogen system, their summed abundance was correlated with disease suppression across all 37 composts, making them promising candidates to serve as indicators for disease-suppressive composts.

Although fungi have been widely reported in compost-mediated disease suppression, for example *Trichoderma spp.* against *R. solani* (Kwok et al., 1987; Hoitink and Boehm, 1999), only few fungal indicators were identified in this study. Several factors may ex-plain this. First, fungal communities in composts were generally less diverse (HerńandezLara et al., 2022; De Corato et al., 2019; Galitskaya et al., 2017) and dominated by few abundant taxa, with limited overlap of rare taxa across samples. Second, greater variability among sequencing replicates suggests a more heterogeneous fungal distribution. Third, sieving composts for DNA extraction excluded larger, less-degraded wood particles. These particles may harbor fungi, such as cellulolytic species, which are often associated with less-humidified organic matter (Zhang et al., 2017) and are abundant in composts (Langarica-Fuentes et al., 2014). Thus, refining sampling and DNA extraction methods could improve fungal coverage and increase the number of indicative fungi. Nonetheless, our results suggest that bacteria are more promising indicators of disease-suppressive composts than fungi.

To enable comparison with existing literature, we focused on indicative ASVs classified to at least the genus level. However, the relevance of ASVs classified only at higher taxonomic resolution should not be overlooked. These may be identified in the future as novel taxa are described. Additionally, also these indicative ASVs hold promise for metaanalyses and may serve as genetic markers for diagnosing disease-suppressive composts (Freeland, 2017).

For the fungi, only one indicative ASV was assigned to a genus previously linked to disease suppression: an ASV associated with *G. ultimum* suppression in cucumber, classified as *Mortierella*. This genus was recently identified as a key indicator of soil suppressiveness against *G. ultimum* (Kurm et al., 2023). For the bacteria, ASVs indicative of *R. solani*-suppression in cucumber, were assigned to the genera *Streptomyces* and *Mycobacterium*, for which both suppressive activity has previously been reported (Wang et al., 2015; Postma et al., 2005). Both genera belong to the phylum Actinomycetota, to which a quarter of all ASVs indicative of *R. solani*-suppression were assigned to, and which is known for harboring many disease-suppressive strains, particularly also associated with compost and soil suppressiveness against *R. solani* (Díaz-Díaz et al., 2022; Mendes et al., 2011; Cordovez et al., 2015).

Among the bacterial ASVs indicative of *G. ultimum*-suppression, we identified 26 genera with promising links to the literature regarding disease suppression (Table S14). Many of them were reported enriched in soils suppressive to *Fusarium* or bacterial wilt with some genera specifically linked to compost-mediated effects, i.e. the genera *Agrobacterium*, *Glutamicibacter*, *Streptomyces* and *Luteimonas* (Charest et al., 2005; Borker et al., 2021; Wang et al., 2015). Particularly, the genus *Luteimonas* has been reported as an indicator of composts suppressive of *Globisporanigum irregulare* in lettuce (HernándezLara et al., 2022) and *Rhizoctonia solani* and *Sclerotinia minor* in cress (Scotti et al., 2020). Indicative ASVs assigned to this genus were not only found among the top 20 for suppression of *G. ultimum* in cucumber, but also among the ASVs indicative of *G. ultimum* independently of the host plant. Moreover, among the ASVs that were indicative for *G. ultimum*-suppression in the current and in our previous study (Mayerhofer et al., 2021a) we found ASVs classified to the genera *Algoriphagus*, *Devosia*, *Povalibacter* and *Sphingopyxis*, all of which were previously reported in the context of disease suppression (Zheng et al., 2024; Liu et al., 2024; Ou et al., 2019; Snelders et al., 2020).

Many of the genera associated with ASVs indicative of *G. ultimum* suppression were also previously found to be enriched in the rhizosphere of tomato plants grown in *Fusarium*-suppressive soils (Jaiswal et al., 2017). These include *Brevundimonas*, *Devosia*, *Cellvibrio*, *Chthoniobacter*, *Flavobacterium*, *Microvirga*, *Novosphingobium*, *Shinella*, *Sphingobium*, and *Sphingopyxis*. This observation is particularly noteworthy given the growing body of evidence suggesting that disease suppression is conferred by complex microbial networks, rather than by individual strains (Liu et al., 2023). Thus, it is plausible that the identified bacteria contribute to *G. ultimum* suppression collectively, functioning as part of a microbial consortium.

### 4.3. Towards general indicators for disease-suppressive composts and the use of compost microorganisms for crop protection

The identified bacteria (and few fungi) indicative for suppressive composts represent a promising starting point for predicting disease suppression by composts. However, their presence or relative abundance in composts is not sufficient to robustly model disease suppression. Composts provide a complex growing environment for microorganisms and it is well described that the activity particularly of microorganisms with biocontrol activity depends on the properties of the substrate they are applied with (and applied to) (Bonanomi et al., 2018). Thus, we should aim for predictive models that integrate microbiome data with compost properties. We attempted such modeling, but it was hindered by two main limitations: strong composting site effects and too few observations compared to the number of potential predictors. Although, to our knowledge, this is the largest dataset of composts tested for disease suppression and profiled for bacterial and fungal communities to date.

To overcome these limitations, we propose narrowing down the scope, for instance, by focusing on compost at similar maturation stages, produced from similar feedstock, or tracking composts over time. Moreover, meta-studies, as successfully applied to suppressive soils (Zhang et al., 2022; Sagova-Mareckova et al., 2023; Wang and Zou, 2024), could help identify general predictors. Additionally, the observation that composts with large differences in abitoic and biotic proprieties can show suppressive activity, suggests that suppression results from several mechanisms (Hadar and Papadopoulou, 2012). Disentangling these mechanisms *in situ* and directly comparing composts with similar functional profiles would be crucial but remains experimentally challenging.

However, even with extensive datasets, the high taxonomic variability among composts likely limits the discovery of universal microbial indicators. Similar challenges have been reported in identifying taxonomic indicators of compost maturation (Estrella-González et al., 2020). We suggest that despite the high taxonomic diversity, functional redundancy exists among compost microorganisms, as has been reported for soil microbiota (Chen et al., 2022). Metagenomic approaches, such as shotgun sequencing, will help identify functional genes associated with disease suppression, as shown in studies on suppressive soils (Carríon et al., 2019; Tracanna et al., 2021). However, these sequencing technologies are still costly and the analysis and interpretation of complex communities—such as those found in compost— challenging (Anthony et al., 2024). In parallel, we need studies that expand our knowledge of compost microbiota potentially involved in disease suppression, for which the current study sets an important foundation.

Our study identified promising bacterial and few fungal indicators that will support the development of diagnostic tools and guide the targeted isolation of key microorgan-isms. Testing these microorganisms for disease-suppressive activity, individually and in consortia, combined with genomic and metabolic profiling, could reveal genes and functions involved in suppression (Lutz et al., 2020). This would deepen our understanding of the mechanisms driving disease suppression and support the development of reliable predictors and potentially microbial inoculants to enhance compost suppressiveness. Together, these strategies could advance the use of compost as a sustainable solution for promoting plant health.

## CRediT authorship contribution statement

**Anja Logo:** Writing — original draft, Writing — review & editing, Methodology, Visualization, Formal analysis, Conceptualization, Investigation. **Benedikt Boppŕe**: Investigation, Methodology, Writing — review & editing. **Jacques Fuchs**: Conceptualization, Resources, Funding acquisition. **Monika Maurhofer**: Writing — review & editing, Supervision. **Thomas Oberhänsli**: Project administration, Methodology, Conceptualization, Funding acquisition, Writing — review & editing. **Barbara Thürig**: Writing — review & editing, Methodology, Conceptualization, Funding acquisition. **Franco Widmer**: Project administration, Funding acquisition, Conceptualization. **Pascale Flury**: Writing — original draft, Writing — review & editing, Methodology, Conceptualization, Supervision, Funding acquisition. **Johanna Mayerhofer**: Writing — original draft, Writing — review & editing, Formal analysis, Conceptualization, Supervision, Methodology, Funding acquisition.

## Declaration of generative AI and AI-assisted technologies in the writing process

During the preparation of this work the authors used *Writing Assistant* by Mohammed Ziara (ChatGPT 4o, OpenAI) and DeepL in order to fix spelling, grammar and enhancing clarity of the text. After using this tool, the authors reviewed and edited the content as needed and take full responsibility for the content of the published article.

## Data Availability

All raw sequences have been deposited at the NCBI Sequence Read Archive under the BioProject accession number PRJNA1200637 (bacteria) and PRJNA1201072 (fungi). R code and session info are available on GitHub (https://github.com/logoa/Bioassay_ Bacteria_paper_1.git, public once accepted for publication).

## Acronyms

C_org_/N_tot_: carbon-nitrogen ratio.
G. ultimum: Globisporangium ultimum.
R. solani: Rhizoctonia solani.
ASV: amplicon sequence variant.
C_org_: total organic carbon.
CaCl_2_: calcium chloride.
DS: dry substance.
FDA: fluorescein diacetate.
N_min_: total mineral nitrogen.
N_tot_: total nitrogen.
NH_4_^+^: ammonium.
NO_2_^-^: nitrite.
NO_3_^-^: nitrate.
NO_3_^-^-N/N_min_: nitrification index.
OD_550_: soluble humic substances.
PCA: Principle Component Analysis.
PO_4_^3-^: phosphate.
WHC: water holding capacity.

## Acknowledgments

We thank all the people that supported us in the lab, in particular Malgorzata Glowala (former intern at FiBL), Tabea Koch (Agroscope), Nadine Peter (FiBL) and Sonja Rein-hard (FiBL), as well as Eva Burgunder and Mónica Camareno Rodriquez (both University of Basel), Miro Zehnder, Martina Fischbach, Alicia Nussbaumer, Verena Huber, Dominik Moser, Nicolas Enggist (former interns or civil servants at FiBL). We also thank Jan Wächli (University of Basel) for the support with the statistics and the Analytic team at FiBL, in particular Anton Kuhn and Adolphe Munyangabe, for their support for physicochemical and microbial compost characterization. We are grateful to the composting facilities in North-western Switzerland that provided us the tested composts. We gratefully acknowledge the Functional Genomics Center Zurich (FGCZ) of University of Zurich and ETH Zurich, and in particular Maria Domenica Moccia, for the support in Amplicon sequencing.

