## Supplementary Materials for "Analyses of 37 composts revealed microbial taxa associated with disease suppressiveness"

Additional Material and Methods, Figures and Tables for the paper "*Analyses of 37 composts revealed microbial taxa associated with disease suppressiveness*".

*Authors*: Anja Logo, Benedikt Boppré, Jacques Fuchs, Monika Maurhofer, Thomas Oberhänsli, Barbara Thürig, Franco Widmer, Johanna Mayerhofer & Pascale Flury

*Date*: May 28, 2025

**Supplementary Figures** **7**

**Supplementary Tables** **24**

#### Supplementary Material and Methods

*A) Physicochemical properties of composts*

Water and $CaCl_2$-extracts were prepared by adding 50 g of fresh compost to 500 mL of deionized water or to 500 mL 0.01 M $CaCl_2$, and shaking horizontally at 75 rpm for 60 min. The pH of the compost was measured in the unfiltered $CaCl_2$-extract directly after shaking using a pH-meter (SevenEasy, Mettler Toledo, Columbus, OH, USA). Both extracts were filtered through cellulose filter paper (MN 619, Machery-Nagel, Düren, Germany). The salinity was assessed by measuring the electrical conductivity in the water-extract (FiveGo Conductivity Meter, Mettler-Toledo, Columbus OH, USA) and converting to compost salinity using the following formula: salinity [$KCl_{equivalent}$ (kg $DS)^{-1}$] = electrical conductivity [mS $cm^{-1}$] x 583.4 / DS [%]. The content of soluble humic substances ($OD_{550}$), which can be used as an indicator for compost maturation (Oshins et al., 2022), was measured in the water-extract using a spectrophotometer at a wavelength of 550 nm (Genesys 150, ThermoFisher Scientific, Waltham, MA, USA). Mineral nutrients of the $CaCl_2$-extracts were determined with the Berthelot's reagent for ammonium (Krom, 1980), the cadmium-reduction method (Gal et al., 2004) for nitrate and nitrite and Molybdenum blue method (Drummond and Maher, 1995) for $PO_4^{3-}$ and measured using Smartchem 450 Discrete Analyser (AMS Alliance, Guidonia, Italy).

*B) Microbial activity of composts*

To assess the microbial activity of the composts, the water content of compost samples was adjusted to 60% of the maximum WHC and the samples were incubated for equilibration at 25°C for seven days. FDA hydrolysis [$\mu$g FDA (min g $DS)^{-1}$] of composts was assessed by shaking 1 g of compost in 20 mL phosphate buffer (60 mM, pH 7.6) containing 400 mg FDA (98%, Aldrich, USA) at 90 rpm for 20 min. The reaction was stopped by adding 20 mL Acetone (99.5%, Roth, Germany). Fluorescence was measured with a fluorescence spectrometer (excitation at 490 nm, emission at 525 nm, TECAN infinite 200 PRO, Switzerland). A negative control sample without the addition of FDA was included in the analysis.

To assess basal respiration of the composts, 10 g dry substance equivalent of fresh compost was weighed into perforated 25 mL centrifuge tubes and placed in 250 mL Schott bottles in the presence of 0.1 M NaOH as a $CO_2$-trap. After a 24 h incubation period at 25°C the NaOH in the bottle was replaced with 40 mL of fresh NaOH and further incubated for 120 h with once exchanging the NaOH after 48 h. The remaining NaOH was determined by automated titration (808 Titrando, Methorm, Zofingen, Switzerland) with 0.1M HCl after precipitation of absorbed $C0_2$ by excess $BaCl_2$. Based on the amount of acid used for titration, the respiration [mg $CO^2$-C $(h)^{-1}$] was calculated according to Alef and Nannipieri (1995) and averaged over the two incubation periods.

*C) Statistical methods for association analysis*

Point Biserial corrleation (setting "r.g") and Indicator Species Analysis ("indVal.g") were applied using the "*multipatt*" function (*Indicspecies*, Cáceres and Legendre (2009)). Differential abundance was assessed using MaAsLin2 and ALDEx2, both of which were recommended in the method comparison study by Nearing et al. (2022). MaAsLin2 was run with default settings except for arcsine square-root transformation and no data scaling (Mallick et al., 2021). For ALDEx2 we employed the "*aldex.clr*" function with log-ratio transformation, followed by "*aldex.test*" for group differences and "*aldex.effect*" for effect sizes (Fernandes et al., 2014). Rarefied read counts were used for PBC, ISA, and MaAsLin2 and unrarefied read counts for ALDEx2.

### Supplementary Figures

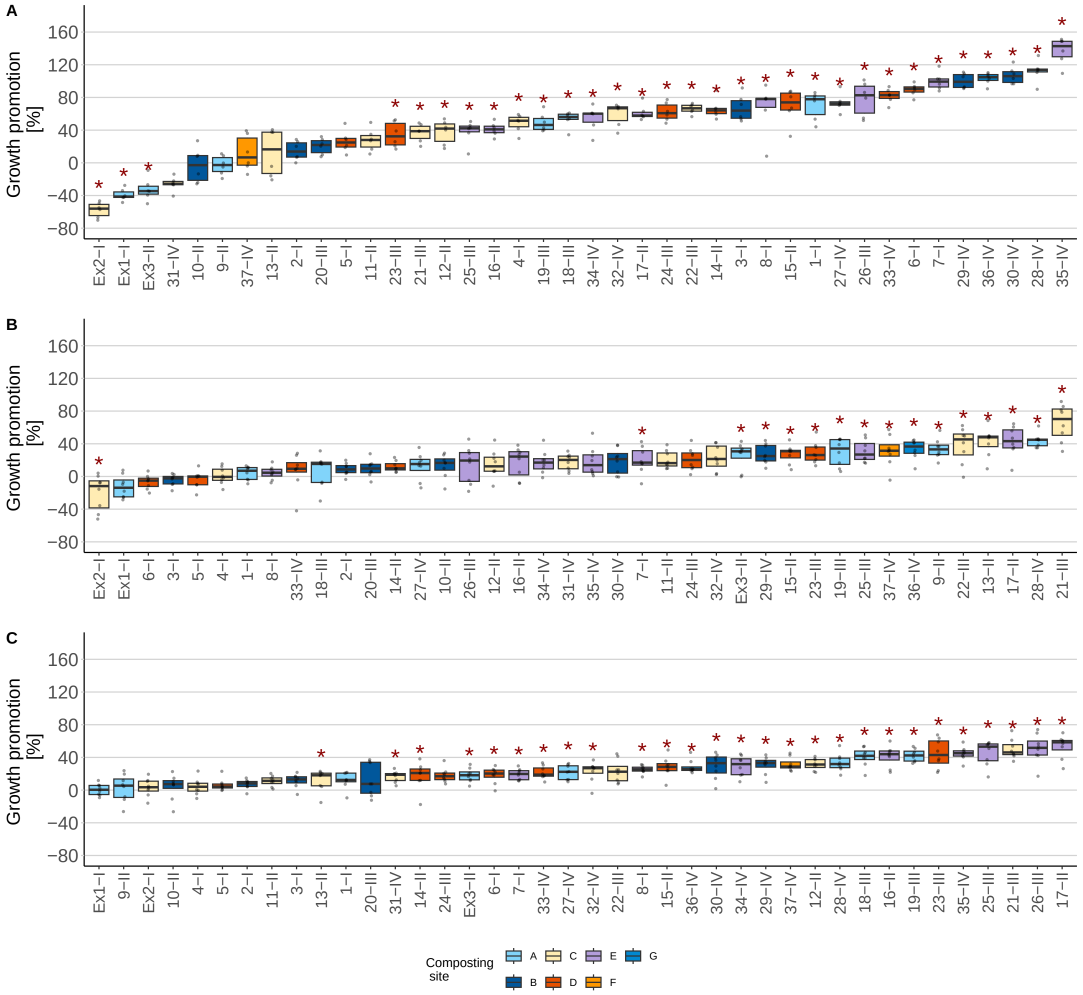

Figure S1: **Growth promotion of composts 1-37 and excluded composts Ex1-3.**: (A) Cress-*G. ultimum*, (B) cucumber-*G. ultimum*, and (C) cucumber-*R. solani*. Growth promotion was calculated by dividing the shoot biomass without pathogen by the average shoot biomass without pathogen of the no-compost control (NC) of the respective compost set. The roman numbers (I-IV) after the compost number indicate the compost set (I = May 2022, II = July 2022, III = September 2022, IV = May 2023). Grey points represent growth promotion for individual pots, boxes show the interquartile range, and dashes denote the median of six replicates for cress or eight replicates for the cucumber systems. Treatments are ordered by increasing mean. Red asterisks indicate significant differences between compost treatments and the no-compost control of the respective set (Tukey HSD test, $p$ <0.05, or pairwise Wilcoxon test, $p$ <0.05). Box colors highlight the composting site.

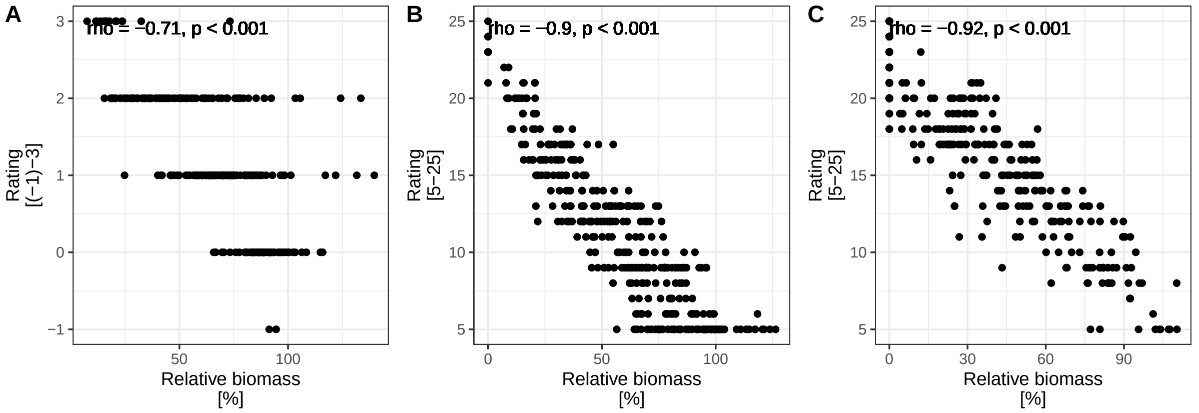

Figure S2: **Spearman Rank Sum correlation between relative biomass and disease symptoms rating for the three plant-pathogen-systems** (A) cress-*G. ultimum*-system (B) cucumber-*G. ultimum*-system and (C) cucumber-*R. solani*-system For pathogen concentrations see caption Figure S3. For the cucumber assays, each seedling was rated separately and was summed for all five seedlings. The following disease ratings were used: cress-*G. ultimum*: -1 better growth than peat without pathogen, 0 healthy, 1 some yellow leaves, reduced growth, 2 half of leaves yellow, growth substantially reduced, 3 majority of leaves yellow, growth extremely reduced; cucumber-*G. ultimum*: 1 healthy, 2 reduced growth, 3 reduced growth, yellow, deformed, 4 death after emergence, 5 not emerged; cucumber-*R. solani*: 1 healthy, 2 brown lesions at the stem but still standing, 3 brown lesions and bend over, 4 dead after emergence, 5 not emerged.

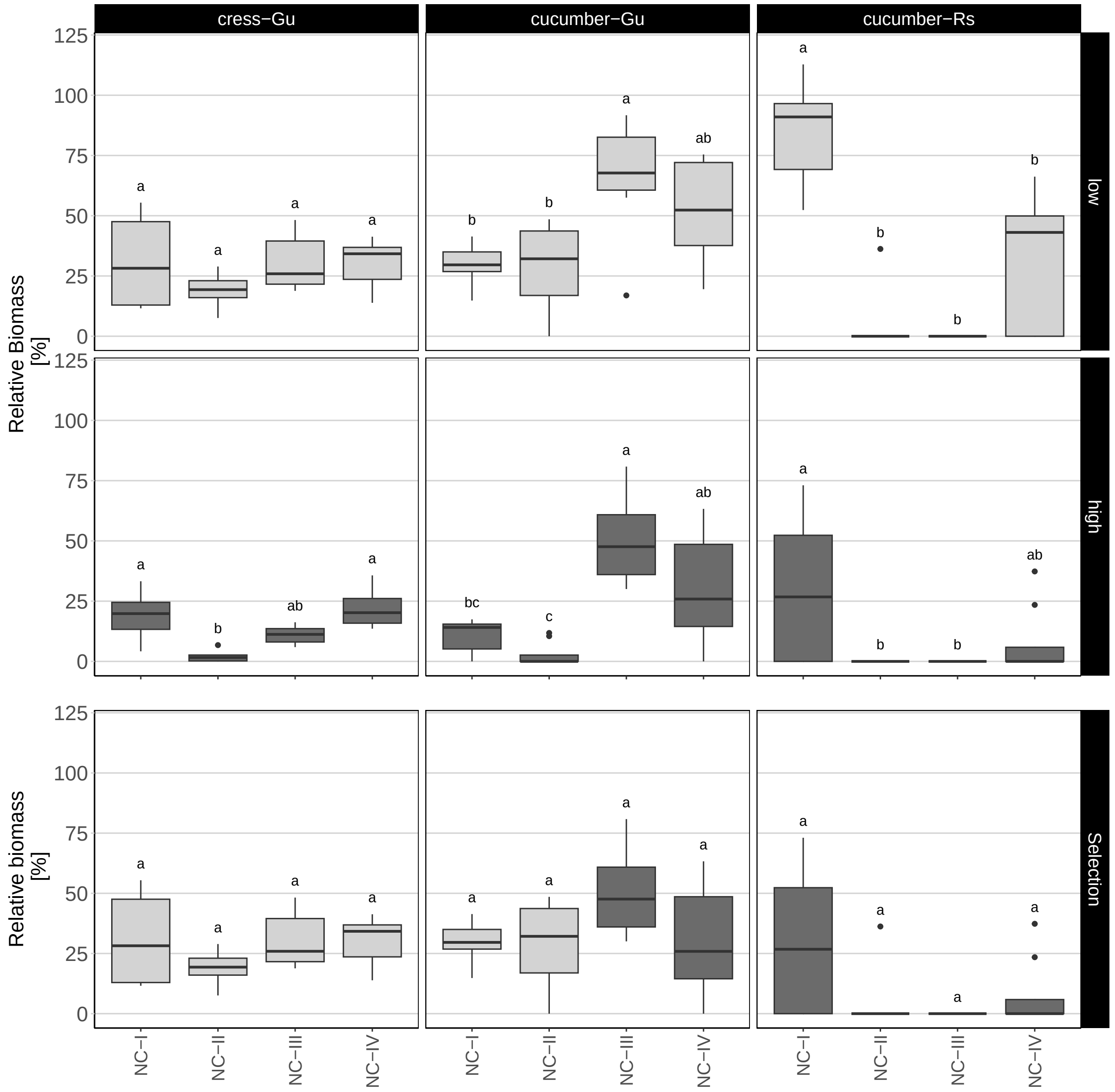

Figure S3: **Comparison of disease-suppressive activity of the no-compost control across experiments**. Pathogen pressure can vary between experiments. Therefore, two pathogen concentrations were tested for all experiments. For *G. ultimum* "low" corresponds to 0.45 g pathogen per liter of substrate and "high" for 1.35 g per liter. For *R. solani* "low" corresponds to three agar plugs with a diameter of 0.8 cm covered with *R. solani* and "high" for 1.4 cm plugs. In batch IV for *R. solani*, "low" corresponds to 0.6 cm plugs, and "high to 0.8 cm plugs. For each compost set, the pathogen concentration was selected to ensure that the relative biomass in the no-compost control was not significantly different among compost sets ( ANOVA followed by Tukey HSD test, $p$ <0.05, or Kruskal-Wallis followed by pairwise Wilcoxon test, $p$ <0.05). This resulted in the following concentrations: Cress-*G. ultimum*: 0.45 g *G. ultimum* per liter of substrate in sets I-IV, Cucumber-*G. ultimum*: 0.45 g *G. ultimum* per liter of substrate in sets I and II, and 1.35 g *G. ultimum* per liter of substrate in sets III and IV, cucumber-*R. solani*: three plugs with a diameter of 0.8 cm in sets II–IV and 1.4 cm in set I.

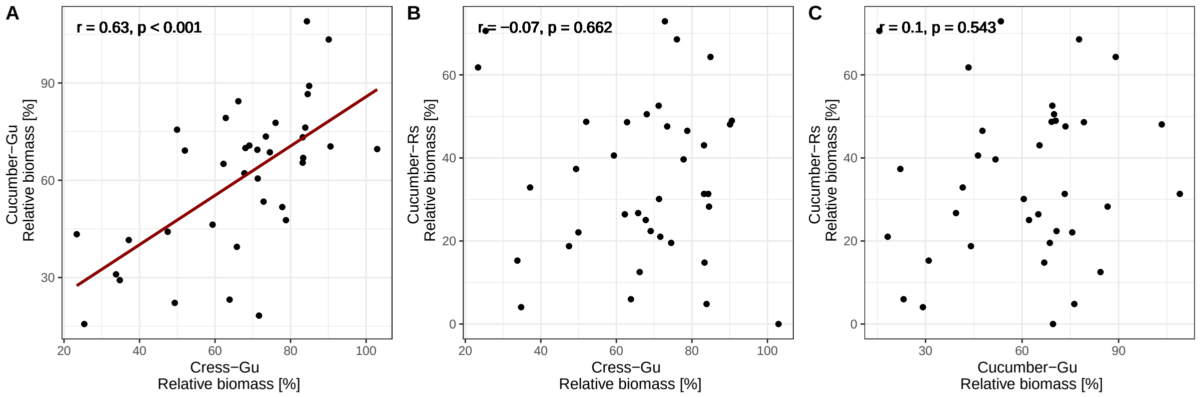

Figure S4: **Correlation of disease-supressive activity of composts between the plant-pathogens systems:** (A) cress-*G. ultimum* and cucumber-*G. ultimum*, (B) cress-*G. ultimum* and cucumber-*R. solani*, (C) cucumber-*G. ultimum* and cucumber-*R. solani*. Disease suppression is shown as relative biomass calculated by dividing the shoot biomass per pot with pathogen by the average shoot biomass of all pots of the same treatment without pathogen.

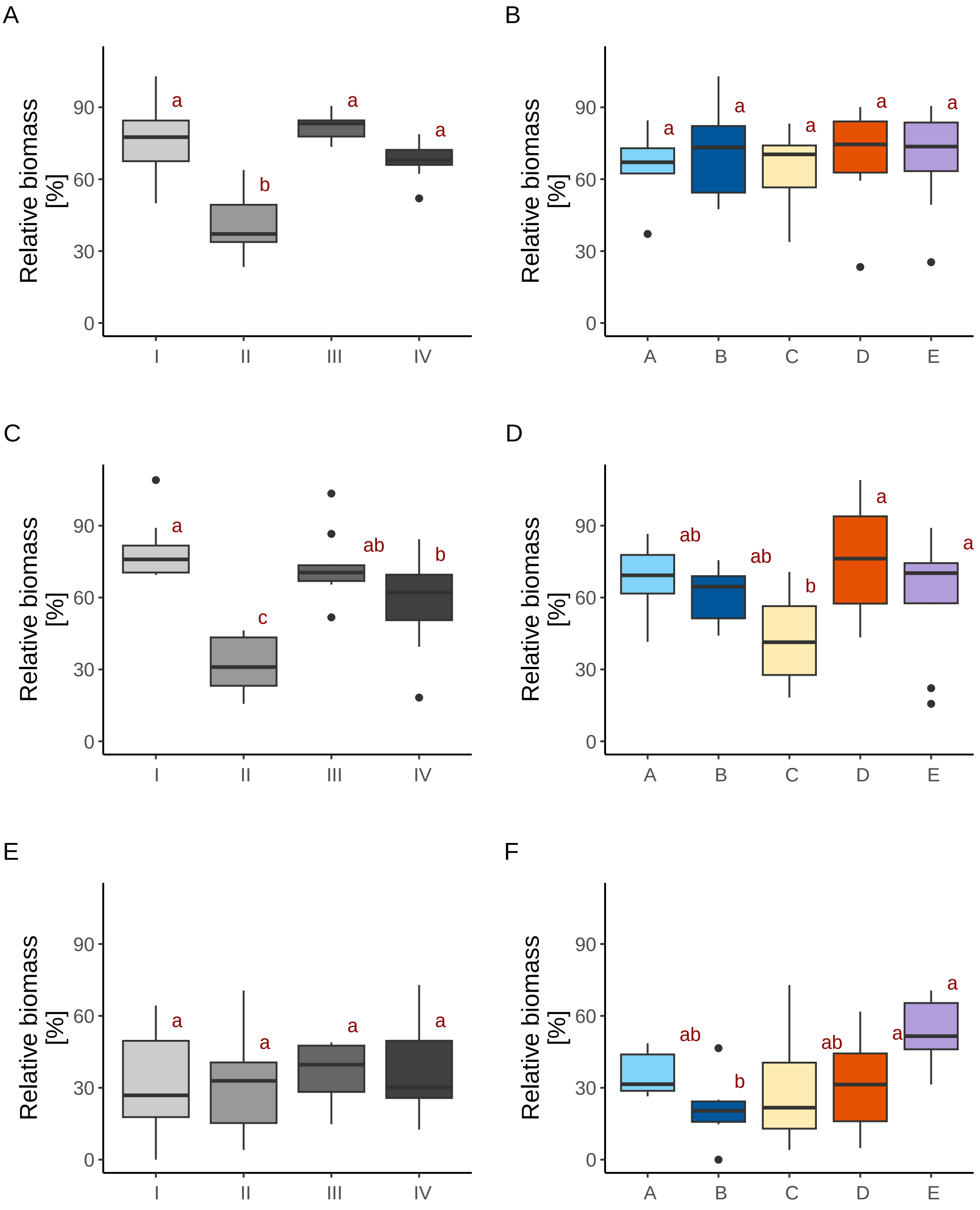

Figure S5: **Disease-suppressive activity of composts grouped by (A, C, E) compost set and (B, D, F) composting site** for (A, B) cress-*G. ultimum*, (C, D) cucumber-*G. ultimum* and (E, F) cucumber-*R. solani*. Composts originating from sites F and G had only one sample each. Therefore, these two composts (36, 37) were excluded from the analysis. Boxes show the interquartile range and dashes denote the median. Groups sharing the same letter within a plot are not statistically different (ANOVA followed by pairwise Tukey HSD test ($p$ <0.05)). Statistics: compost set from top to bottom: F = 21.97, $p$ <0.001, F = 16.79, $p$ <0.001, F = 0.34, $p$ = 0.799 and composting site: F = 0.18, $p$ = 0.948, F = 2.43, $p$ = 0.07, F = 3.43, $p$ = 0.02

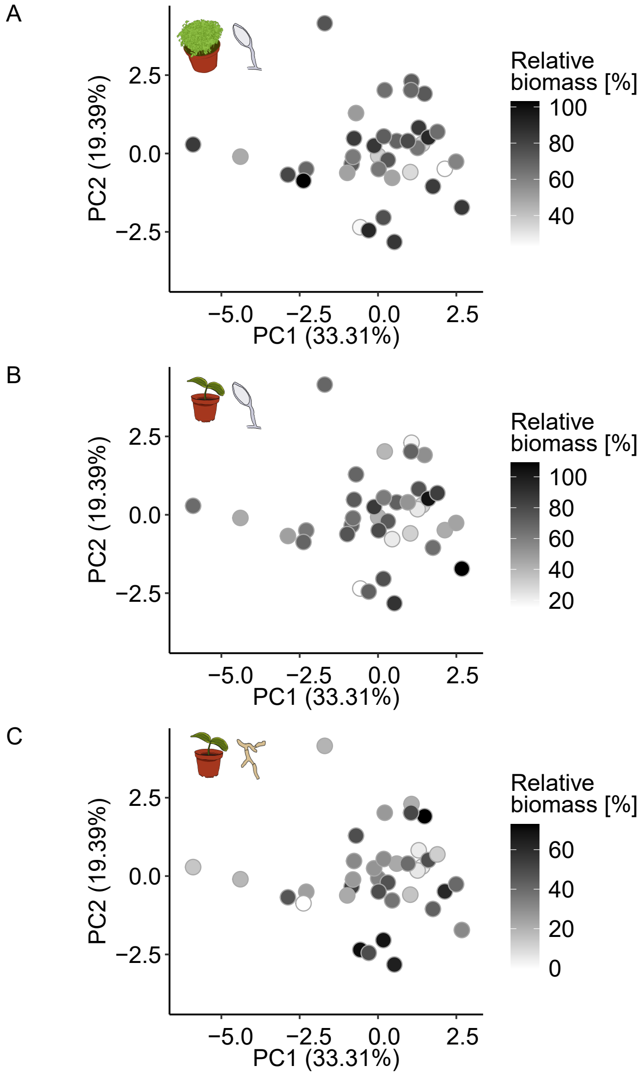

Figure S6: **Principal Component Analysis based on compost properties.** The Euclidean distance between composts was calculated based on nine selected compost properties (indicated with an asterisk in Table 1). Strongly co-correlated compost properties ( $|\rho|$ >0.8, Supplementary Figure S7) and properties with missing data (basal respiration, compost age) or which were calculated from other properties (nitrification index, $N_{min}$, $C_{org}/N_{tot}$) were removed. The plots display the first and second principal component. Dots represent composts colored according to their disease suppression in (A) the cress-*G. ultimum*-system, (B) the cucumber-*G. ultimum*-system and (C) the cucumber-*R. solani*-system.

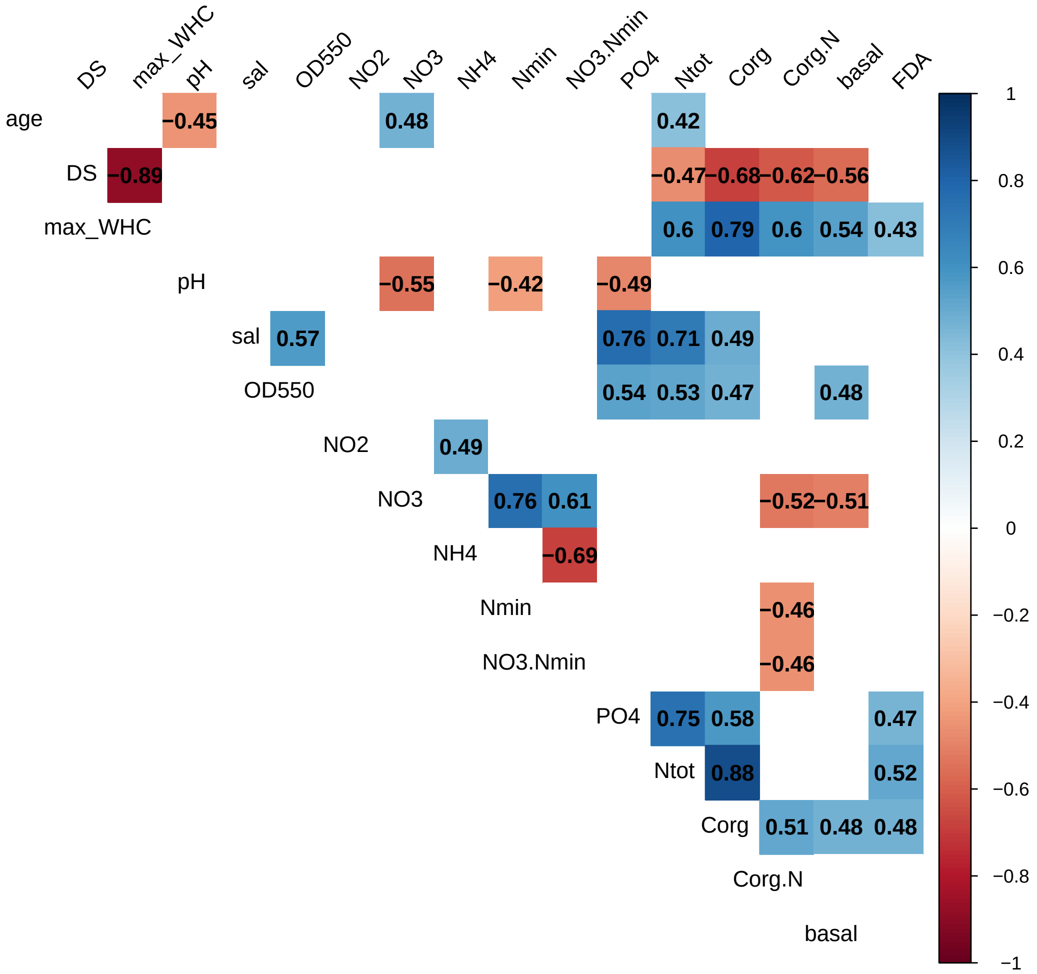

Figure S7: **Correlation among abiotic and biotic compost properties**. *P*-values were adjusted for multiple testing using Benjamini & Hochberg correction. Pearson's correlation was used when both properties were normally distributed, otherwise Spearman's Rank sum test.

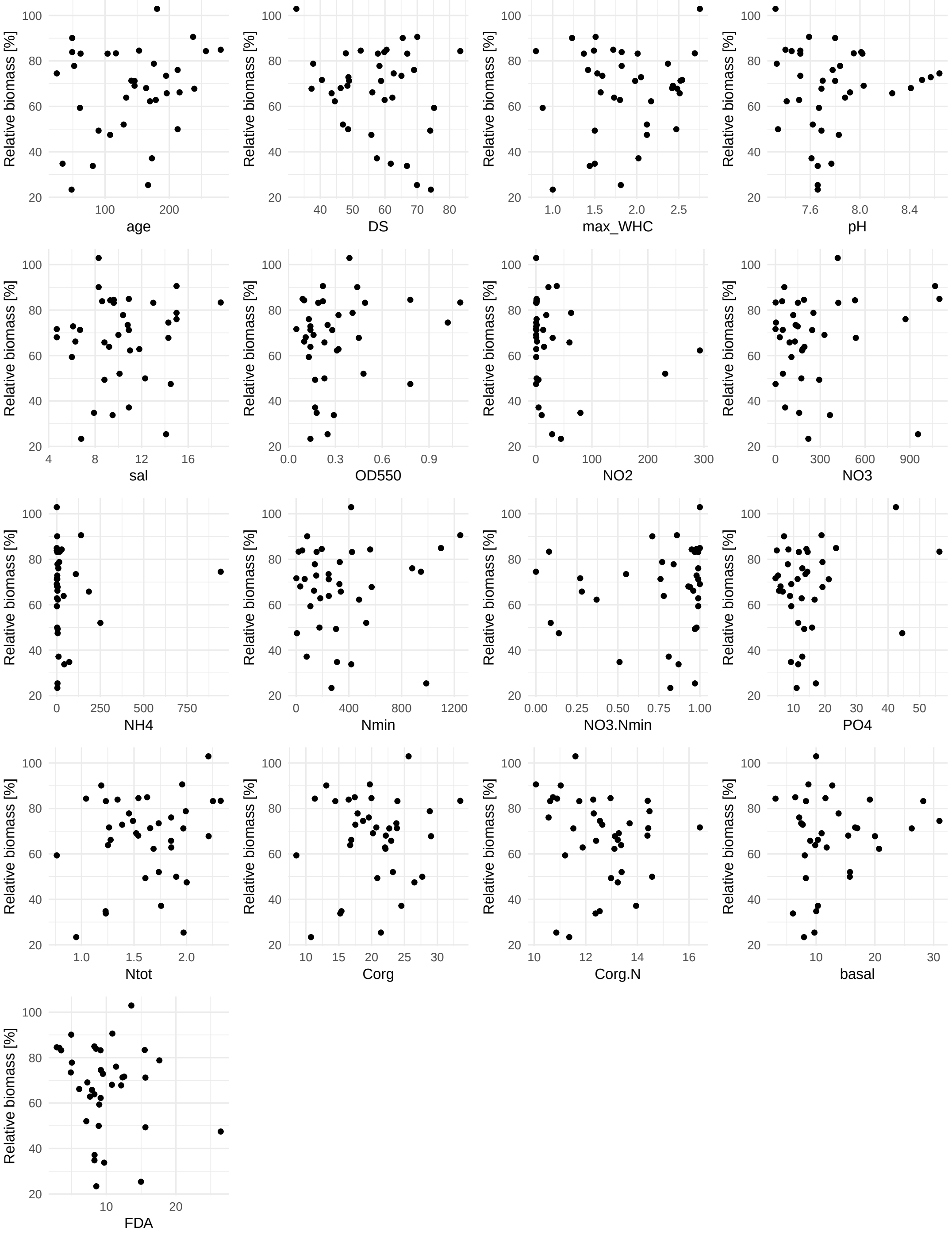

Figure S8: **Correlation plots of mean disease suppression in the cress-*G. ultimum*-system and compost properties** such as compost age [d], dry substance (DS) [%], water holding capacity [g $H_20$ (g DS)$^{-1}$], pH, salinity (sal) [g $KCl_{eq}$ (kg DS)$^{-1}$], Soluble humic substances ($OD_{550}$), $NO_2^-$-N [mg (kg DS)$^{-1}$] ,$NO_3^-$-N [mg (kg DS)$^{-1}$] , $NH_4^+$-N [mg (kg DS)$^{-1}$] , total mineral nitrogen ($N_{min}$) [mg (kg DS)$^{-1}$] , $NO_3^-$-N/$N_{min}$-ratio, $PO_4^{3-}$ [mg (kg DS)$^{-1}$] , total nitrogen ($N_{tot}$) [%], total organic carbon ($C_{org}$) [%],$C_{org}/N_{tot}$, basal respiration (basal) [mg $CO^2$-C (h)$^{-1}$] , FDA hydrolysis [$\mu$g FDA (min g DS)$^{-1}$] . Correlations were based on 37 composts, except for compost age 35 and basal respiration only 34 were available. Statistics are shown in Table 1.

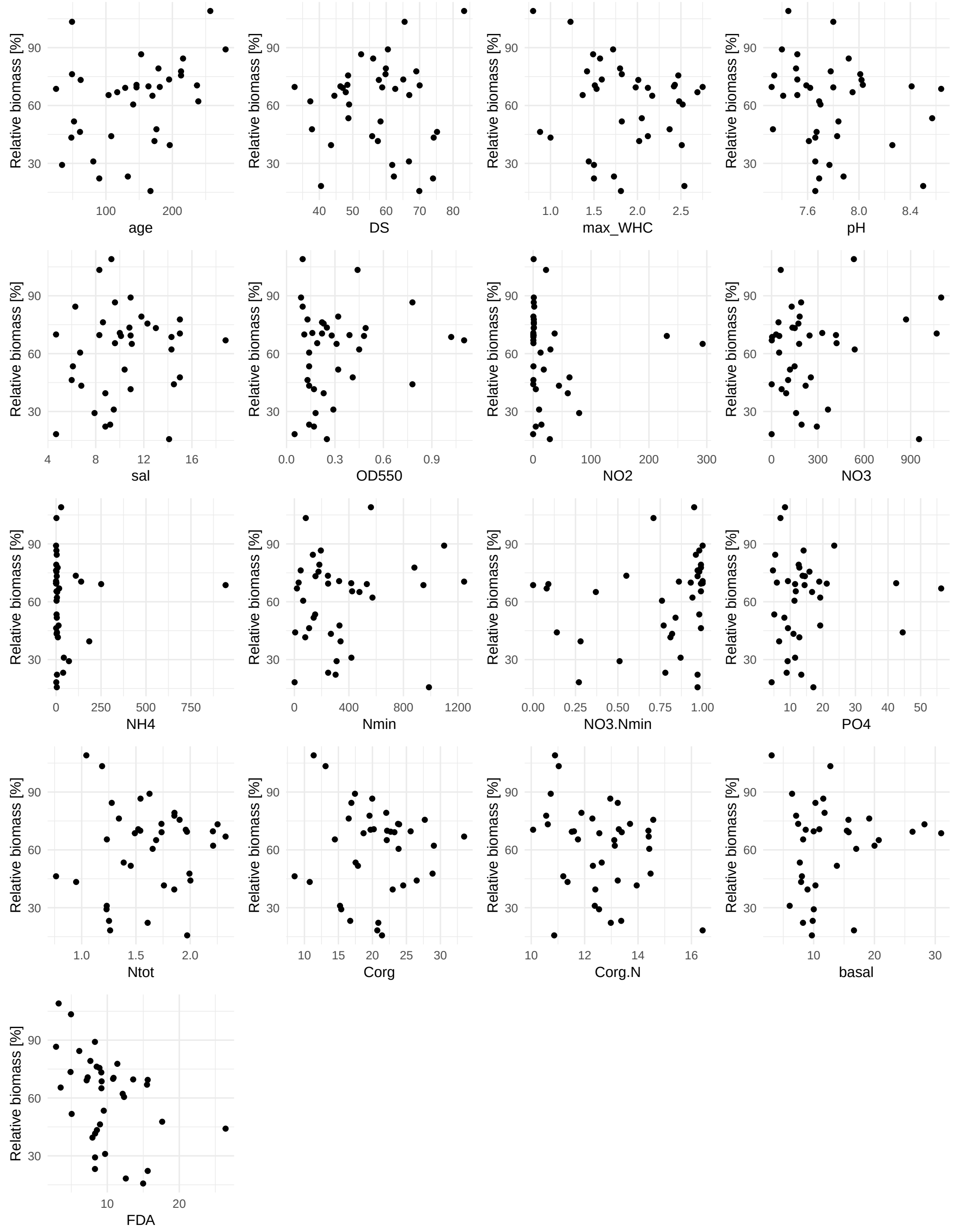

Figure S9: **Correlation plots of mean disease suppression in the cucumber-*G. ultimum*-system and compost properties** such as compost age [d], dry substance (DS) [%], water holding capacity [g $H_20$ (g DS)$^{-1}$], pH, salinity (sal) [g $KCl_{eq}$ (kg DS)$^{-1}$], Soluble humic substances ($OD_{550}$), $NO_2^-$-N [mg (kg DS)$^{-1}$] ,$NO_3^-$-N [mg (kg DS)$^{-1}$] , $NH_4^+$-N [mg (kg DS)$^{-1}$] , total mineral nitrogen ($N_{min}$) [mg (kg DS)$^{-1}$] , $NO_3^-$-N/$N_{min}$-ratio, $PO_4^{3-}$ [mg (kg DS)$^{-1}$] , total nitrogen ($N_{tot}$) [%], total organic carbon ($C_{org}$) [%],$C_{org}/N_{tot}$, basal respiration (basal) [mg $CO^2$-C (h)$^{-1}$] , FDA hydrolysis [$\mu$g FDA (min g DS)$^{-1}$] . Correlations were based on 37 composts, except for compost age 35 and basal respiration only 34 were available. Statistics are shown in Table 1.

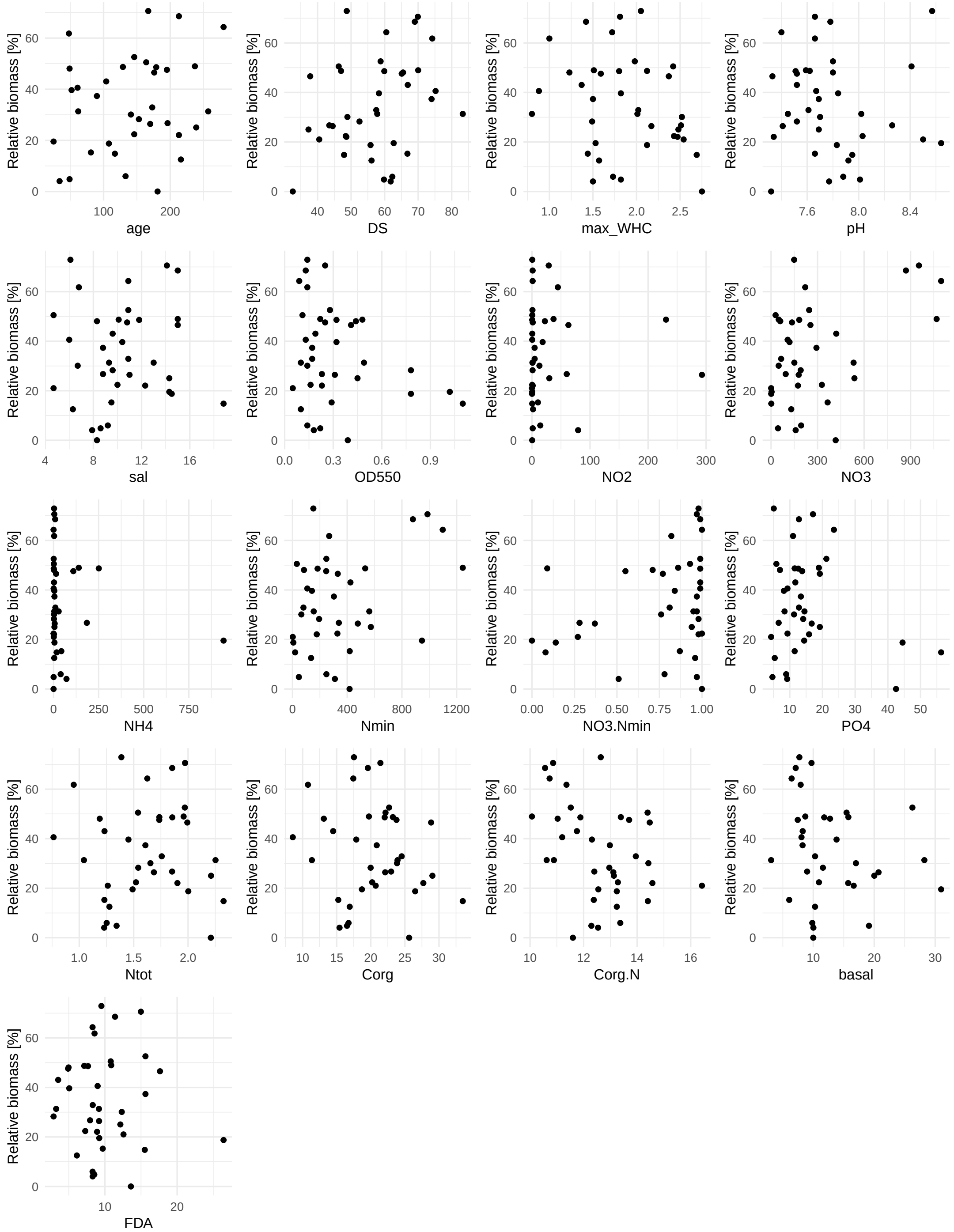

Figure S10: **Correlation plots of mean disease suppression in the cucumber-*R. solani*-system and compost properties** such as compost age [d], dry substance (DS) [%], water holding capacity [g $H_20$ (g DS)$^{-1}$], pH, salinity (sal) [g $KCl_{eq}$ (kg DS)$^{-1}$], Soluble humic substances ($OD_{550}$), $NO_2^-$-N [mg (kg DS)$^{-1}$] ,$NO_3^-$-N [mg (kg DS)$^{-1}$] , $NH_4^+$-N [mg (kg DS)$^{-1}$] , total mineral nitrogen ($N_{min}$) [mg (kg DS)$^{-1}$] , $NO_3^-$-N/$N_{min}$-ratio, $PO_4^{3-}$ [mg (kg DS)$^{-1}$] , total nitrogen ($N_{tot}$) [%], total organic carbon ($C_{org}$) [%],$C_{org}/N_{tot}$, basal respiration (basal) [mg $CO^2$-C (h)$^{-1}$] , FDA hydrolysis [$\mu$g FDA (min g DS)$^{-1}$] . Correlations were based on 37 composts, except for compost age 35 and basal respiration only 34 were available. Statistics are shown in Table 1.

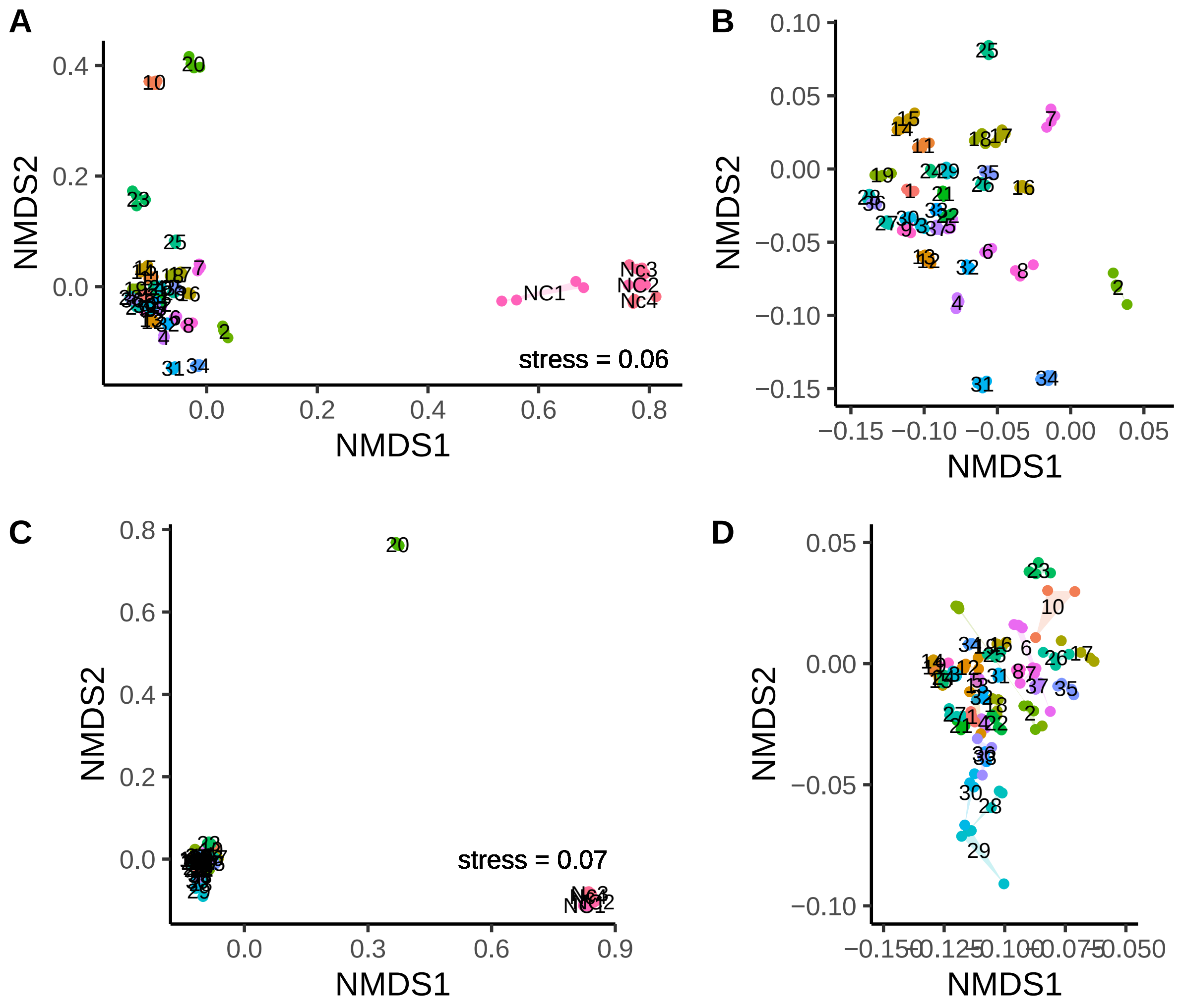

Figure S11: **Beta diversity based on Bray-Curtis dissimilarity was assessed among all (A, C) 164 bacterial samples and (C, D) 162 fungal samples**. The samples are colored by substrate ID (compost 1-17, non-compost control NC 1-4). The four replicates of each substrate are connected to form a polygon. The left plot displays all samples, while the right plot provides a zoomed-in view of the point cloud containing the majority of the samples.

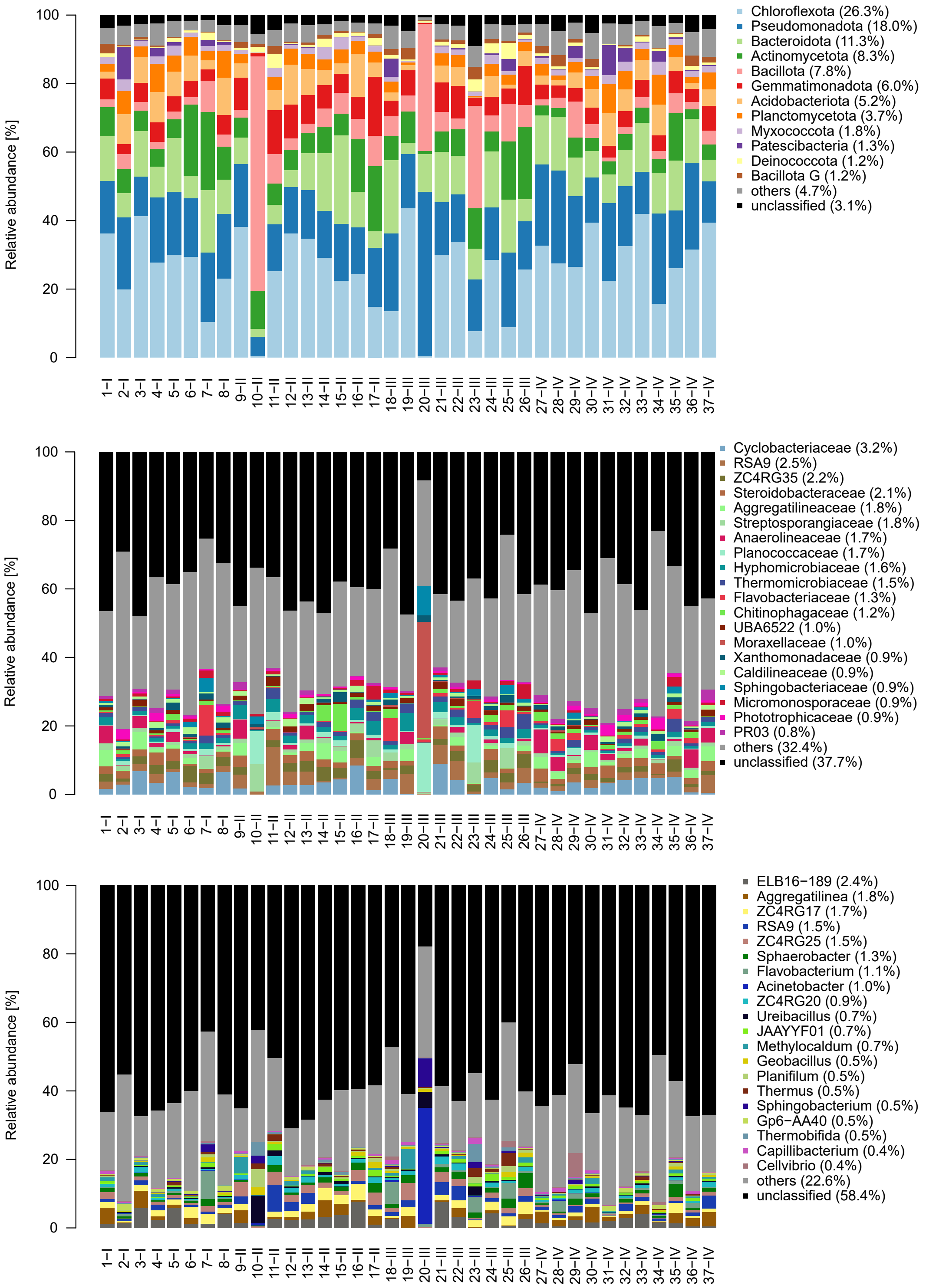

Figure S12: **Taxonomic composition of the bacterial communities of the 37 composts** at (A) phylum, (B) family and (C) genus level. Taxonomic composition was calculated based on rarefied read numbers and averaged for the sequencing replicates (n=4).

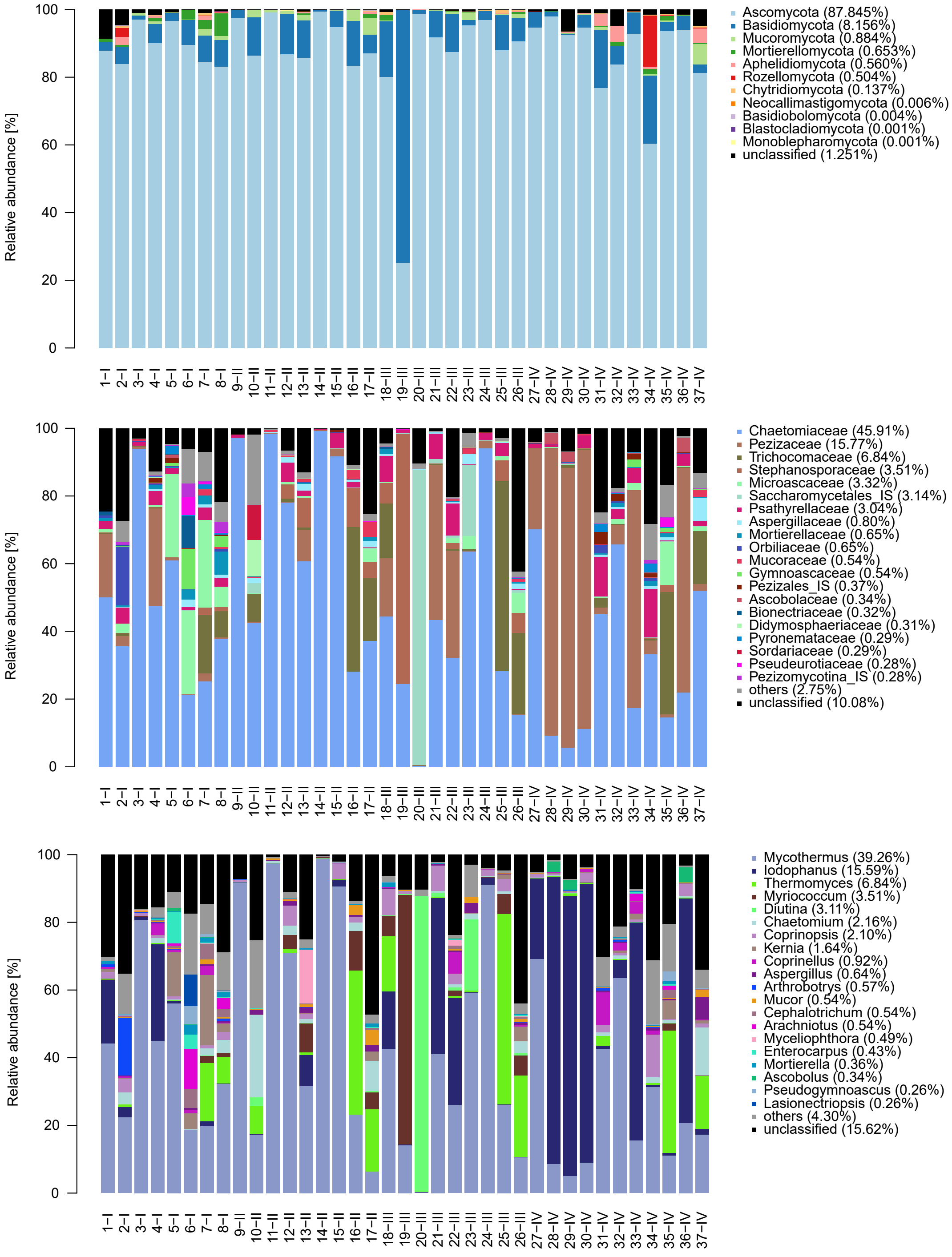

Figure S13: **Taxonomic composition of the fungal communities of the 37 composts** at (A) phylum, (B) family and (C) genus level. Taxonomic composition was calculated based on rarefied read numbers and averaged for the sequencing replicates (n=4).

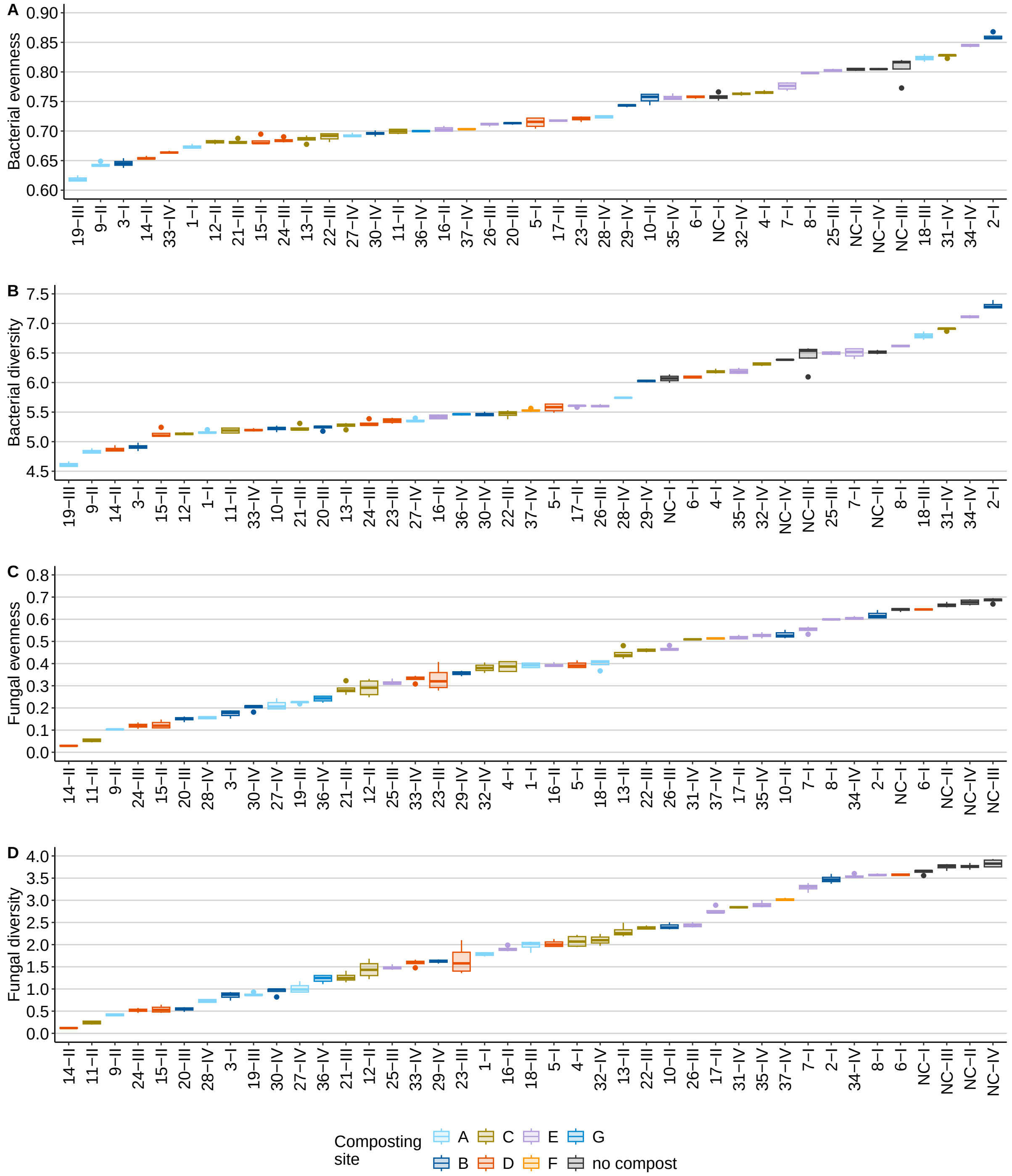

Figure S14: **Shannon eveness and Shannon diversity index of the bacterial communities and fungal communities**. Boxplots display (A) bacterial eveness, (B) bacterial diversity, (C) fungal evenness and (D) fungal diversity of the 37 composts (1-37) and the no-compost control (NC) of the four compost sets (I-IV) ordered by their means (n =4). The boxes represent the interquartile range (IQR, $25^{th}$ to $75^{th}$ percentile), the horizontal line within the box the median and the whiskers shows the minima and maxima of the data up to 1.5 x IQR. Outliers are shown as individual data points. Boxplots are ordered by mean.

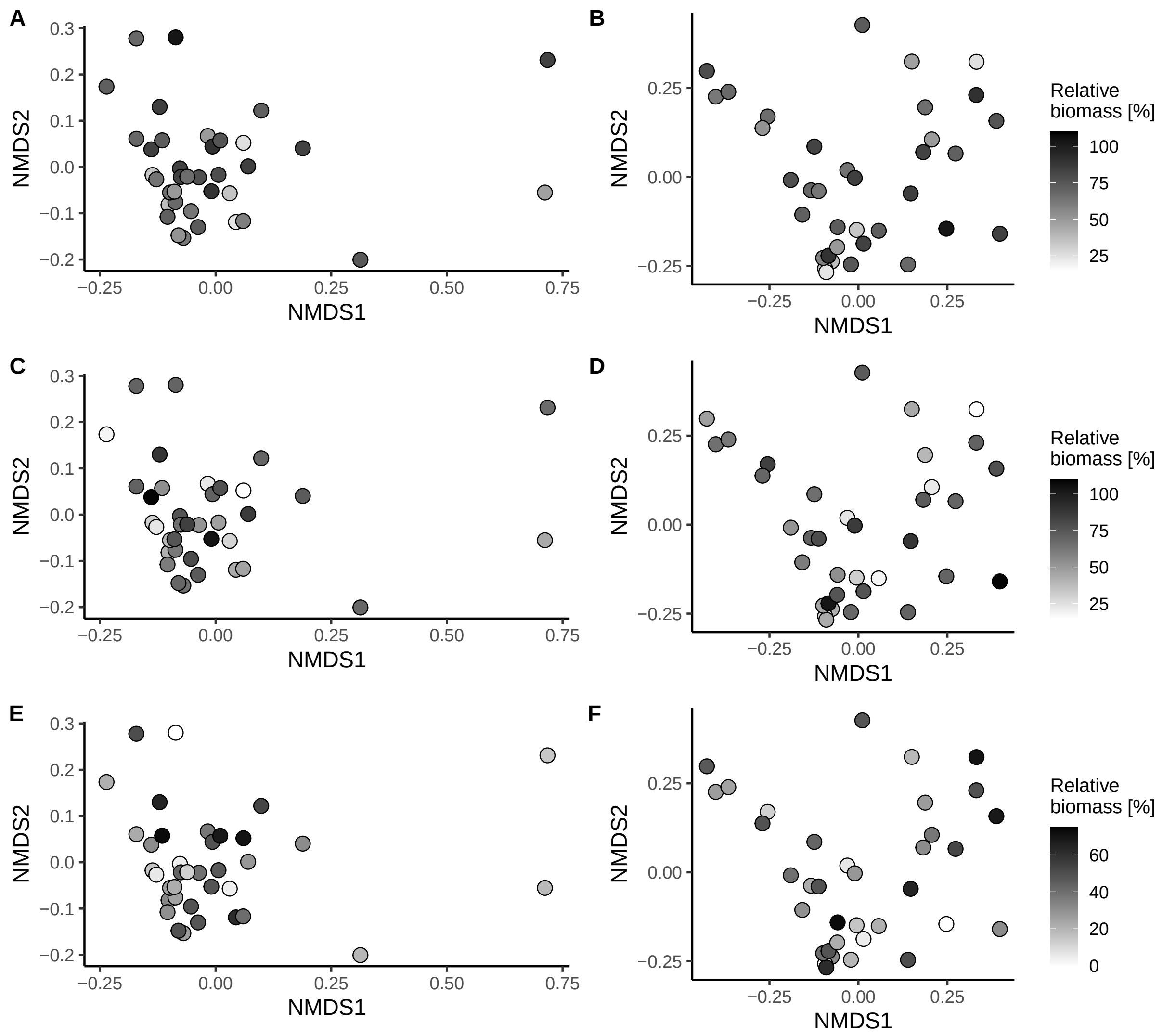

Figure S15: **(A, C, E) bacterial and (B, D, F) fungal community structure colored by disease-suppressive activity.** Community structure is based on the mean Bray-Curtis dissimilarity of four replicates for 37 composts for bacterial communities and 36 composts for fungal communities (a plot including the here excluded compost 20 is shown in Supplementary Figure S11). The ordination plot is colored by the disease-suppressive activity of the composts in (A, B) the cress-*G. ultimum* system, (C, D) the cucumber-*G. ultimum* system and (E, F) the cucumber-*R. solani* system.

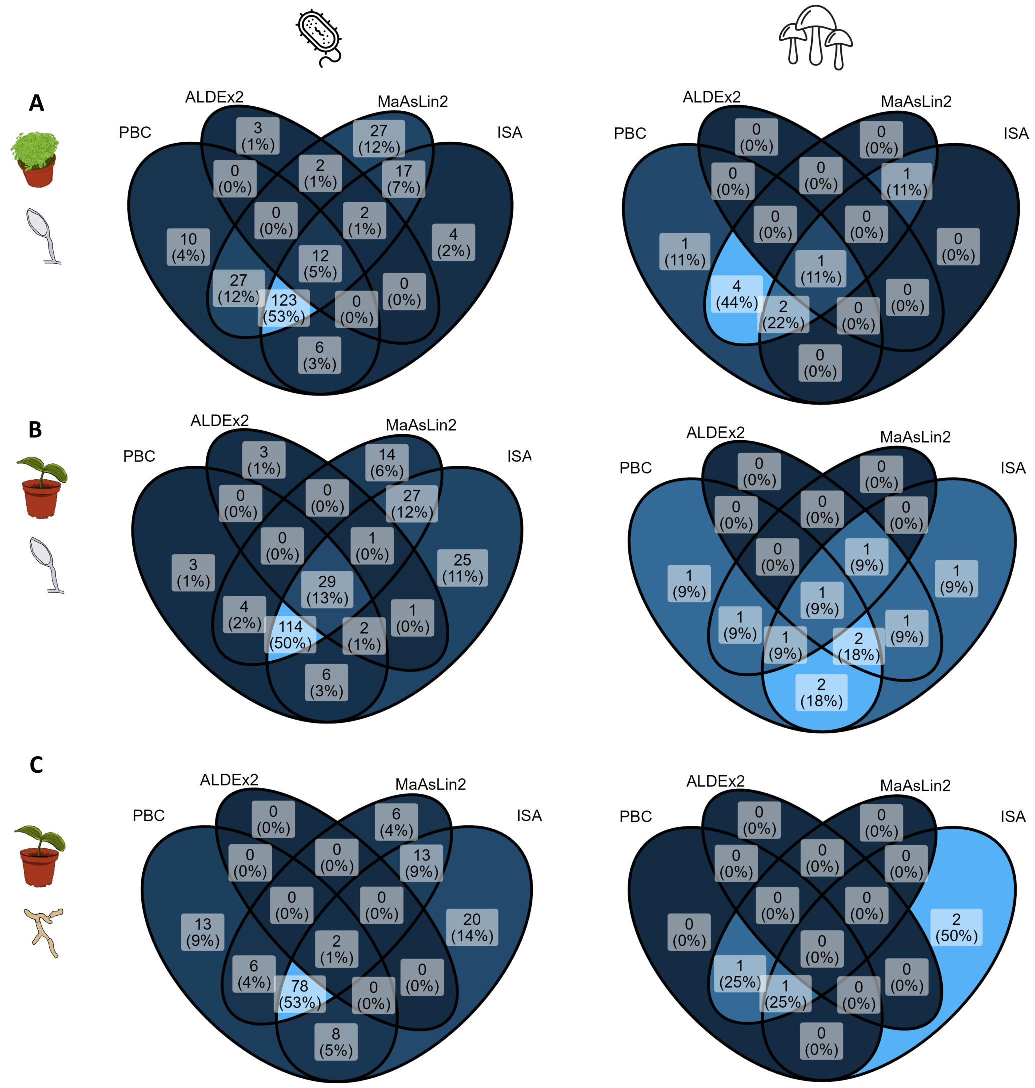

Figure S16: **Number of ASVs indicative of disease-suppressive composts identified by different statistical methods in the three plant-pathogen systems (A) cress-*G. ultimum*, (B) cucumber-*G. ultimum*, (C) and cucumber-*R. solani*.** The left column shows results for bacteria, while the right column shows results for fungi. Association tests were performed by comparing ASVs between the nine most suppressive and nine least suppressive composts for each plant-pathogen system. Only ASVs present in at least six out of the 18 selected composts were considered. ASVs were considered indicative if they were statistically significantly associated with the most suppressive composts ($p < 0.05$) in at least three out of four statistical methods (Point-biserial correlation (PBC), Indicator species analysis (ISA), MaAsLin2, ALDEx2). Venn diagrams were constructed using the `ggVennDiagram` function from the same-named R package (Gao and Dusa, 2024).

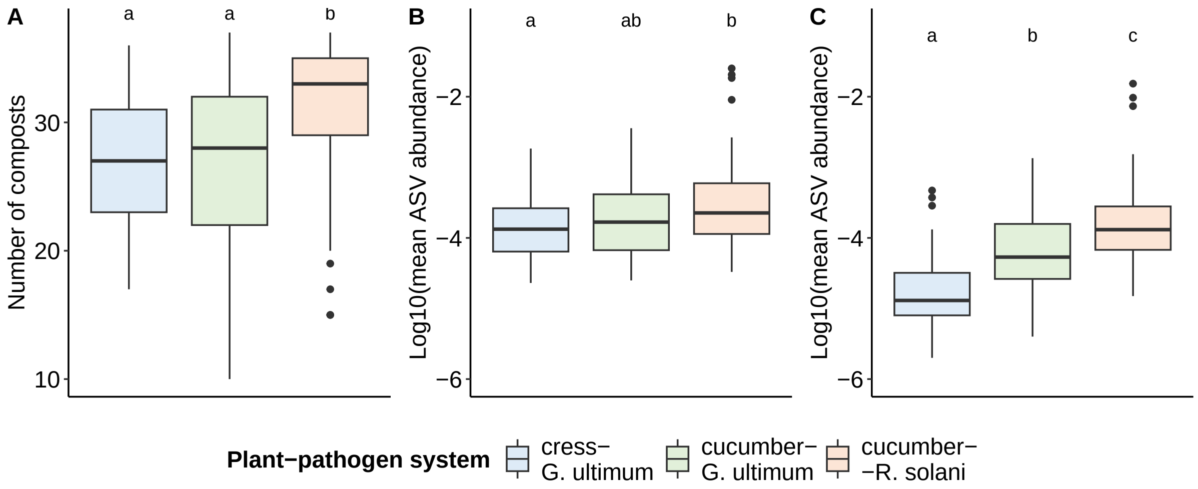

Figure S17: **Prevalence of indicative ASVs**. Boxplots displays: (A) the number of composts the indicative ASVs were detected, (B) their mean relative abundance across the disease-suppressive composts, (C) their mean relative abundance across non-suppressive composts for each plant-pathogen system (137, 146 and 80 ASVs, respectively). For plotting the relative abundances were log10 transformed. Suppressive and non-suppressive composts for each plant-pathogen system are identified in Figure 1. Statistical significance was assessed using Kruskal-Wallis tests followed by pairwise Wilcox tests: (A) Chi-squared = 47.4, $p < 0.001$; (B) Chi-squared = 12.9, $p = 0.002$; and (C) Chi-squared = 136.4, $p < 0.001$ (C).

#### Supplementary Tables

Table S1: **Overview composting process for the different composting sites.** During the thermophilic phase of the composting process, all composting companies turned the composting piles regularly, monitored the temperature development, and adjusted the humidity by watering the compost pile when necessary. Triangular compost piles were additionally covered with a compost fleece to control humidity in the compost pile.

| Site | Composting process | Feedstock composition |
|---|---|---|
| A | Tabular piles with turning every 1-2 weeks<br>Compost is sieved and stored covered<br>Storage is turned manually once a week to keep aerated | 10-45% green waste from the gardener<br>10-40% leaves<br>25-40% hay/grass<br>5-15% variable inputs: soil, mature compost, digestate (sieve residue) |
| B | Tabular piles with turning every 1-2 weeks<br>Compost is sieved and stored covered (for gardening) or uncovered (agricultural composts: 2, 10, 29)<br>Covered storage is turned manually once a week to keep aerated | 0-20% green waste<br>0-10% leaves<br>60-80% digestate sieve residue<br>5-20% variable inputs: leaves, manure, digestate, cellulose, paper mill waste |
| C | Small triangle piles with 2-3 turnings per week<br>Maturation in large piles, covered with a fleece but not aerated<br>Sieving and covered storage | 75% green waste from the gardener<br>20% mature compost<br>5% soil |
| D | Small triangle piles with 2-3 turnings per week<br>Compag system for two weeks for sanitation and volume reduction<br>Aeration during covered storage | 55-65% municipal green waste<br>20% horse manure<br>10% soil<br>5% vegetable waste<br>0-10% mature compost |
| E | Small triangle piles on field edge<br>Turning only a few times<br>Sieving and storage in small covered piles | 100% green waste from farm area |
| F | Small triangle piles with 1-2 turnings per week<br>Sieving and storage in small covered piles | 100% municipal green waste |
| G | Tabular piles with turning every 1-2 weeks<br>Sieving and storage in small covered piles<br>Storage is turned manually once a week to keep aerated | 25-30% woody green waste<br>40% leaves<br>20-25% green waste |

Table S2: **Composition, composting site, compost age of all 40 composts.** Pick up set I-IV ($16^{th}$ of May, $18^{th}$ of July, $26^{th}$ of September 2022, $28^{th}$ of April 2023). All information were obtained from the composting protocol provided by the compost producers. For compost 31 and 32 the protocol was missing.

| **ID** | **Set** | **Site** | **Start date** | **Compost age** | **Feedstock composition** |
|---|---|---|---|---|---|
| Ex1 | I | A | *26.01.2022* | 110 | green waste (30%), leaves (30%), grass (30%), mature compost (5%), digestate (5%) |
| 1 | I | A | *18.11.2021* | 179 | green waste (10%), leaves (40%), grass (25%), mature compost (10%) |
| 2 | I | B | *16.11.2021* | 181 | green waste (10%), leaves (10%), digestate sieve residue (70%) |
| 3 | I | B | *15.10.2021* | 213 | green waste (15%), leaves (5%),digestate sieve residue (60%), digestate (20%) |
| Ex2 | I | C | *07.03.2022* | 70 | green waste (75%), soil (5%), mature compost (20%) |
| 4 | I | C | *21.12.2021* | 146 | green waste (75%), soil (5%), mature compost (20%) |
| 5 | I | D | *28.03.2022* | 49 | green waste (65%), soil (10%), manure (20%), vegetable waste (5%) |
| 6 | I | D | *01.09.2021* | 257 | green waste (55%), soil (10%), mature compost (10%), manure (20%), vegetable waste (5%) |
| 7 | I | E | *21.12.2021* | 146 | green waste (100%) |
| 8 | I | E | *09.08.2021* | 280 | green waste (100%) |
| Ex-3 | II | A | *15.03.2022* | 125 | green waste (35%), leaves (30%), grass (25%), soil (10%), digestate sieve residue (5%) |
| 9 | II | A | *26.01.2022* | 173 | green waste (30%), leaves (30%), grass (30%), mature compost (5%), digestate (5%) |
| 10 | II | B | *01.04.2022* | 108 | green waste (20%), digestate sieve residue (70%), other (10%) |
| 11 | II | C | *14.06.2022* | 34 | green waste (75%), mature compost (20%), soil (5%) |
| 12 | II | C | *28.04.2022* | 81 | green waste (75%), mature compost (20%), soil (5%) |
| 13 | II | C | *07.03.2022* | 133 | green waste (75%), mature compost (20%), soil (5%) |
| 14 | II | D | *31.05.2022* | 48 | green waste (55%), soil (10%), mature compost (10%), manure (20%), vegetable waste (5%) |
| 15 | II | D | *18.05.2022* | 61 | green waste (55%), soil (10%), mature compost (10%), manure (20%), vegetable waste (5%) |
| 16 | II | E | *19.04.2022* | 90 | green waste (100%) |
| 17 | II | E | *01.02.2022* | 167 | green waste (100%) |
| 18 | III | A | *26.04.2022* | 153 | green waste (25%), leaves (30%), grass (40%), digestate sieve residue (5%) |
| 19 | III | A | *15.03.2022* | 195 | green waste (35%), leaves (30%), grass (25%), soil (5%), mature compost (5%) |
| 20 | III | B | *01.06.2022* | 117 | green waste (15%), digestate sieve residue (80%), other (5%) |
| 21 | III | C | *05.08.2022* | 52 | green waste (75%), soil (5%), mature compost (20%) |
| 22 | III | C | *14.06.2022* | 104 | green waste (75%), soil (5%), mature compost (20%) |

*Continued on next page*

| ID | Set | Site | Start date | Compost age | Feedstock composition |
|---|---|---|---|---|---|
| 23 | III | D | *01.09.2022* | 25 | green waste (55%), soil (10%), mature compost (10%), manure (20%), vegetable waste (5%) |
| 24 | III | D | *08.08.2022* | 49 | green waste (55%), soil (10%), mature compost (10%), manure (20%), vegetable waste (5%) |
| 25 | III | E | *26.07.2022* | 62 | green waste (100%) |
| 26 | III | E | 01.02.2022 | 237 | green waste (100%) |
| 27 | IV | A | 08.12.2022 | 141 | green waste (35%), leaves (40%), grass (25%) |
| 28 | IV | A | 09.11.2022 | 170 | green waste (20%), leaves (35%), grass (40%), digestate sieve residue (5%) |
| 29 | IV | B | 03.11.2022 | 176 | digestate sieve residue (70%), manure (15%), other (15%) |
| 30 | IV | B | 01.09.2022 | 239 | green waste (20%), digestate sieve residue (70%), other (10%) |
| 31 | IV | C | NA | NA | NA |
| 32 | IV | C | NA | NA | NA |
| 33 | IV | D | 24.09.2022 | 216 | green waste (55%), soil (10%), mature compost (10%), manure (20%), vegetable waste (5%) |
| 34 | IV | E | 15.11.2022 | 164 | green waste (100%) |
| 35 | IV | E | 27.09.2022 | 213 | green waste (100%) |
| 36 | IV | G | 20.12.2022 | 129 | green waste (50%), grass (40%), cellulose (10%) |
| 37 | IV | F | 18.10.2022 | 196 | green waste (100%) |

Table S3: **Compost characteristics of the 40 composts**.

| **ID** | **age** | **DS** | **WHC** | **pH** | **sal** | **$OD_{550}$** | **$NO_2^-$-N** | **$NO_3^-$-N** | **$NH_4^+$-N** | **$N_{min}$** | **$NO_3^-$-N/ $N_{min}$** | **$PO_4^{3-}$-P** | **$N_{tot}$** | **$C_{org}$** | **$C_{org}$/ $N_{tot}$** | **basal** | **FDA** |
|---|---|---|---|---|---|---|---|---|---|---|---|---|---|---|---|---|---|
| Ex-1 | 110 | 53.5 | 2.3 | 7.7 | 9.3 | 0.15 | 0.6 | 3.7 | 3.5 | 7.8 | 0.48 | 9.8 | 1.6 | 24.7 | 15.3 | 14.8 | 9.2 |
| 1 | 179 | 59.9 | 1.8 | 7.5 | 11.8 | 0.32 | 0.5 | 181.8 | 1.0 | 183.3 | 0.99 | 12.6 | 1.9 | 22.0 | 11.9 | 11.8 | 7.7 |
| 2 | 181 | 32.6 | 2.8 | 7.3 | 8.3 | 0.39 | 0.4 | 416.8 | 0.0 | 417.2 | 1.00 | 42.5 | 2.2 | 25.6 | 11.6 | 10.0 | 13.6 |
| 3 | 213 | 48.6 | 2.5 | 7.3 | 12.3 | 0.23 | 1.3 | 173.6 | 2.2 | 177.1 | 0.98 | 15.9 | 1.9 | 27.7 | 14.6 | 15.7 | 8.9 |
| Ex-1 | 70 | 56.0 | 2.1 | 8.1 | 7.3 | 0.14 | 0.2 | 6.7 | 0.7 | 7.6 | 0.89 | 3.7 | 1.2 | 17.4 | 14.0 | 8.9 | 6.7 |
| 4 | 146 | 48.4 | 2.4 | 8.0 | 10.0 | 0.16 | 0.4 | 327.9 | 0.0 | 328.3 | 1.00 | 9.3 | 1.5 | 20.2 | 13.3 | 10.9 | 7.3 |
| 5 | 49 | 59.8 | 1.8 | 8.0 | 8.6 | 0.22 | 1.3 | 44.8 | 0.3 | 46.4 | 0.97 | 4.7 | 1.3 | 16.5 | 12.3 | 19.1 | 8.5 |
| 6 | 257 | 83.3 | 0.8 | 7.5 | 9.3 | 0.10 | 0.9 | 532.5 | 28.2 | 561.6 | 0.95 | 8.4 | 1.0 | 11.4 | 10.9 | 3.1 | 3.2 |
| 7 | 146 | 58.8 | 2.0 | 7.8 | 10.9 | 0.28 | 0.9 | 245.8 | 0.7 | 247.4 | 0.99 | 21.2 | 2.0 | 22.7 | 11.5 | 26.3 | 15.6 |
| 8 | 280 | 60.5 | 1.7 | 7.4 | 10.9 | 0.09 | 1.2 | 1097.4 | 0.0 | 1098.6 | 1.00 | 23.5 | 1.6 | 17.4 | 10.7 | 6.4 | 8.3 |
| Ex-2 | 125 | 56.4 | 2.1 | 7.7 | 10.2 | 0.16 | 4.4 | 71.1 | 11.4 | 86.9 | 0.82 | 11.5 | 1.8 | 25.6 | 14.6 | 11.3 | 8.8 |
| 9 | 173 | 57.5 | 2.0 | 7.6 | 10.9 | 0.17 | 4.7 | 65.0 | 10.1 | 79.8 | 0.81 | 12.8 | 1.8 | 24.5 | 14.0 | 10.3 | 8.3 |
| 10 | 108 | 55.8 | 2.1 | 7.8 | 14.5 | 0.78 | 0.2 | 0.9 | 5.1 | 6.2 | 0.14 | 44.5 | 2.0 | 26.5 | 13.2 | NA | 26.4 |
| 11 | 34 | 61.8 | 1.5 | 7.8 | 7.9 | 0.18 | 79.5 | 158.8 | 71.6 | 309.9 | 0.51 | 9.2 | 1.2 | 15.4 | 12.5 | 10.0 | 8.3 |
| 12 | 81 | 66.8 | 1.4 | 7.7 | 9.5 | 0.29 | 10.3 | 364.9 | 43.1 | 418.3 | 0.87 | 11.5 | 1.2 | 15.2 | 12.4 | 6.1 | 9.7 |
| 13 | 133 | 62.3 | 1.7 | 7.9 | 9.2 | 0.14 | 14.4 | 194.1 | 39.4 | 247.9 | 0.78 | 8.9 | 1.3 | 16.7 | 13.4 | 9.8 | 8.3 |
| 14 | 48 | 74.2 | 1.0 | 7.7 | 6.8 | 0.14 | 44.6 | 220.4 | 3.1 | 268.1 | 0.82 | 11.0 | 0.9 | 10.8 | 11.4 | 7.9 | 8.6 |
| 15 | 61 | 75.2 | 0.9 | 7.7 | 6.0 | 0.13 | 0.5 | 107.1 | 0.5 | 108.1 | 0.99 | 9.3 | 0.8 | 8.6 | 11.2 | 8.1 | 9.0 |
| 16 | 90 | 74.0 | 1.5 | 7.7 | 8.8 | 0.17 | 4.6 | 292.9 | 5.0 | 302.5 | 0.97 | 13.4 | 1.6 | 20.9 | 13.0 | 8.2 | 15.6 |
| 17 | 167 | 69.9 | 1.8 | 7.7 | 14.1 | 0.25 | 28.9 | 954.1 | 4.1 | 987.1 | 0.97 | 17.1 | 2.0 | 21.4 | 10.9 | 9.7 | 15.0 |
| 18 | 153 | 52.5 | 1.5 | 7.5 | 9.6 | 0.78 | 1.1 | 191.2 | 2.0 | 194.3 | 0.98 | 14.1 | 1.5 | 20.0 | 13.0 | 11.6 | 2.9 |
| 19 | 195 | 65.1 | 1.6 | 7.5 | 10.8 | 0.25 | 1.4 | 135.1 | 109.6 | 246.1 | 0.55 | 13.8 | 1.7 | 23.8 | 13.7 | 7.5 | 4.9 |
| 20 | 117 | 47.9 | 2.7 | 8.0 | 18.8 | 1.10 | 0.4 | 1.5 | 17.2 | 19.1 | 0.08 | 56.3 | 2.3 | 33.5 | 14.4 | NA | 15.5 |
| 21 | 52 | 58.3 | 1.8 | 7.8 | 10.4 | 0.32 | 18.4 | 119.2 | 4.1 | 141.7 | 0.84 | 8.2 | 1.5 | 17.9 | 12.3 | 13.8 | 5.1 |
| 22 | 104 | 66.9 | 1.4 | 7.5 | 9.6 | 0.19 | 0.6 | 420.6 | 2.5 | 423.7 | 0.99 | 11.7 | 1.2 | 14.5 | 11.8 | 8.3 | 3.5 |
| 23 | 25 | 62.7 | 1.5 | 8.6 | 14.3 | 1.02 | 0.5 | 3.5 | 942.7 | 946.7 | 0.00 | 14.4 | 1.5 | 18.7 | 12.6 | 31.0 | 9.2 |
| 24 | 49 | 65.5 | 1.2 | 7.8 | 8.3 | 0.44 | 22.3 | 59.4 | 2.1 | 83.8 | 0.71 | 7.0 | 1.2 | 13.1 | 11.0 | 12.7 | 5.0 |
| 25 | 62 | 57.8 | 2.0 | 8.0 | 13.0 | 0.49 | 1.0 | 150.2 | 3.8 | 155.0 | 0.97 | 14.5 | 2.3 | 23.9 | 10.6 | 28.2 | 9.2 |
| 26 | 237 | 70.0 | 1.5 | 7.6 | 15.0 | 0.22 | 37.2 | 1068.3 | 139.6 | 1245.1 | 0.86 | 18.9 | 2.0 | 19.7 | 10.1 | 8.7 | 10.9 |
| 27 | 141 | 48.9 | 2.5 | 7.7 | 6.7 | 0.14 | 12.9 | 49.0 | 2.8 | 64.7 | 0.76 | 11.3 | 1.7 | 23.8 | 14.4 | 17.0 | 12.3 |
| 28 | 170 | 44.5 | 2.2 | 7.4 | 11.0 | 0.31 | 292.8 | 178.5 | 6.2 | 477.5 | 0.37 | 16.7 | 1.7 | 22.1 | 13.1 | 20.7 | 9.2 |
| 29 | 176 | 37.8 | 2.4 | 7.3 | 15.0 | 0.41 | 62.8 | 254.1 | 13.9 | 330.8 | 0.77 | 19.2 | 2.0 | 28.8 | 14.5 | NA | 17.6 |
| 30 | 239 | 37.3 | 2.5 | 7.7 | 14.3 | 0.45 | 29.9 | 537.9 | 5.0 | 572.8 | 0.94 | 19.2 | 2.2 | 29.0 | 13.1 | 20.0 | 12.1 |
| 31 | NA | 40.5 | 2.5 | 8.5 | 4.7 | 0.05 | 0.0 | 0.5 | 1.2 | 1.7 | 0.27 | 4.3 | 1.3 | 20.7 | 16.4 | 16.6 | 12.6 |
| 32 | NA | 48.7 | 2.1 | 8.6 | 6.1 | 0.14 | 0.6 | 149.0 | 2.8 | 152.4 | 0.98 | 5.1 | 1.4 | 17.5 | 12.6 | 7.7 | 9.5 |
| 33 | 216 | 56.1 | 1.6 | 7.9 | 6.3 | 0.10 | 1.9 | 130.2 | 3.6 | 135.7 | 0.96 | 5.4 | 1.3 | 16.9 | 13.2 | 10.3 | 6.1 |
| 34 | 164 | 46.3 | 2.4 | 8.4 | 4.7 | 0.11 | 0.6 | 29.2 | 1.5 | 31.3 | 0.93 | 5.9 | 1.5 | 22.2 | 14.4 | 15.5 | 10.8 |
| 35 | 213 | 69.0 | 1.4 | 7.8 | 15.0 | 0.13 | 1.2 | 870.3 | 8.9 | 880.4 | 0.99 | 12.8 | 1.9 | 19.6 | 10.6 | 7.1 | 11.4 |
| 36 | 129 | 47.0 | 2.1 | 7.6 | 10.1 | 0.48 | 230.9 | 49.8 | 250.8 | 531.5 | 0.09 | 11.5 | 1.7 | 23.2 | 13.4 | 15.8 | 7.1 |
| 37 | 196 | 43.5 | 2.5 | 8.3 | 8.8 | 0.23 | 59.9 | 94.5 | 184.5 | 338.9 | 0.28 | 6.6 | 1.9 | 23.0 | 12.4 | 9.0 | 7.9 |

DS = dry substance, WHC = max. waterholding capacity, sal = salinity, $OD_{550}$ = humic soluble substances, basal = basal respiration, FDA = FDA hydrolysis. For units see Table S9

Table S4: **Number of sequences and ASVs for the different filtering steps of sequencing data.** High-quality sequences represent those retained after quality control using the bioinformatics pipeline. For the rarefied dataset, values are provided both with and without the non-compost control (NC). The final column indicates the percentage of sequences or ASVs lost relative to the preceding step. For fungi, three samples were excluded due to having fewer than 10,000 reads.

| | **Step** | **NC included** | **Type** | **Total** | **Min** | **Max** | **Mean** | **Lost (%)** |
|---|---|---|---|---|---|---|---|---|
| **Bacteria** | High-quality sequences | yes | Sequences | 19,933,068 | 36,402 | 330,947 | 121,543 | |
| | | | ASVs | 22,761 | 1,381 | 7,575 | 4,037 | |
| | Bacterial sequences | yes | Sequences | 19,706,220 | 35,918 | 328,121 | 120,160 | 1.1% |
| | | | ASVs | 22,608 | 1,375 | 7,530 | 4,006 | 0.7% |
| | After normalization (35,928 reads) | yes | Sequences | 5,823,664 | 34,741 | 36,758 | 35,510 | 70.4% |
| | | | ASVs | 22,269 | 881 | 5,823 | 2,730 | 1.5% |
| | | no | Sequences | 5,255,759 | 35,741 | 36,758 | 35,512 | |
| | | | ASVs | 19,129 | 881 | 5,823 | 2,696 | |
| **Fungi** | High-quality sequences | yes | Sequences | 23,267,141 | 1,583 | 206,235 | 132,200 | |
| | | | ASVs | 5,078 | 134 | 1,302 | 675 | |
| | Fungal sequences | yes | Sequences | 18,886,962 | 1,313 | 186,565 | 115,164 | 18.8% |
| | | | ASVs | 2,913 | 93 | 740 | 371 | 42.6% |
| | After normalization (10,211 reads) | yes | Sequences | 1,634,025 | 10,084 | 10,234 | 10,149 | 91.3% |
| | | | ASVs | 2,767 | 28 | 379 | 157 | 5.0% |
| | | no | Sequences | 1,472,158 | 10,084 | 10,234 | 10,153 | |
| | | | ASVs | 2,309 | 28 | 379 | 147 | |

Table S5: **Differences in disease-suppressive activity among composts and no-compost controls**. An ANOVA was conducted to compare substrates within each set. For disease suppression in set I within the cucumber-*G. ultimum* system, normality assumptions were violated; therefore, a Kruskal-Wallis test was applied instead.

| **Plant-pathogen system** | **Compost set** | **Pathogen concentration** | **F / Chi-sq** | ***p*** |
|---|---|---|---|---|
| cress-*G. ultimum* | I | 0.45 | 8.38 | 0.001 |
| | II | 0.45 | 6.26 | 0.001 |
| | III | 0.45 | 8.18 | 0.001 |
| | IV | 0.45 | 5.95 | 0.001 |
| cucumber-*G. ultimum* | I | 0.45 | 39.28 | 0.001 |
| | II | 0.45 | 2.42 | 0.019 |
| | III | 1.35 | 5.83 | 0.001 |
| | IV | 1.35 | 9.48 | 0.001 |
| cucumber-*R. solani* | I | 1.4 | 5.59 | 0.001 |
| | II | 0.8 | 7.44 | 0.001 |
| | III | 0.8 | 2.8 | 0.001 |
| | IV | 0.8 | 5.96 | 0.001 |

Table S6: **The five most common ASV classifications at the phylum, family, and genus levels for the 19,129 bacterial and 2,309 fungal ASVs identified across the 37 composts**. Among 64 bacterial phyla, 18 were present in all composts. At the family level, 75.3% of bacterial ASVs were assigned to 759 families, with 90 found in all composts, while 48.2% of bacterial ASVs were assigned to 1,582 genera, 78 of which were shared across all samples. For fungi, 3 of 11 phyla were found in all composts. At the family level, 67.7% of fungal ASVs were assigned to 197 families, with six found in all composts, and at the genus level, 57.8% of fungal ASVs were assigned to 375 genera, seven of which were present in all composts.

| | N | **Phylum** | N | **Family** | N | **Genus** |
|---|---|---|---|---|---|---|
| **Bacteria** | 5481 | *Pseudomonadota* | 510 | *Sphingobacteriaceae* | 324 | *Flavobacterium* |
| | 2309 | *Bacteroidota* | 453 | *Sphingomonadaceae* | 186 | *Sphingobacterium* |
| | 1516 | *Actinomycetota* | 389 | *Chitinophagaceae* | 122 | *Acinetobacter* |
| | 1227 | *Patescibacteria* | 384 | *Flavobacteriaceae* | 92 | *Cellvibrio* |
| | 1161 | *Bacillota* | 328 | *Burkholderiaceae B* | 84 | *Gp6-AA40* |
| | 956 | Unclassified | 4731 | Unclassified | 9902 | Unclassified |
| **Fungi** | 1231 | *Ascomycota* | 114 | *Aspergillaceae* | 54 | *Mucor* |
| | 427 | *Basidiomycota* | 83 | *Microascaceae* | 41 | *Penicillium* |
| | 106 | *Chytridiomycota* | 55 | *Mortierellaceae* | 37 | *Aspergillus* |
| | 87 | *Mucoromycota* | 50 | *Mucoraceae* | 26 | *Coprinopsis* |
| | 68 | *Mortierellomycota* | 67 | *Chaetomiaceae* | 25 | *Talaromyces* |
| | 289 | Unclassified | 750 | Unclassified | 975 | Unclassified |

N = Number of ASVS

Table S7: **ANOVA tables for differences in alpha diversity metrics among composts.**

| **Group** | **Metric** | **F-value** | ***p*** |
|---|---|---|---|
| | Richness | 778.7 | <0.001 |
| Bacteria | Shannon evenness | 450.2 | <0.001 |
| | Shannon diversity | 587.8 | <0.001 |
| | Richness | 347 | <0.001 |
| Fungi | Shannon evenness | 448 | <0.001 |
| | Shannon diversity | 543.9 | <0.001 |

Table S8: **Relationships between compost properties and microbial activity with the bacterial and fungal communities.** The analysis was based on all 37 composts, except for basal respiration (n = 34) and compost age (n = 35). Statistics: Spearman's rank-sum correlation was used to assess the relationship between alpha diversity metrics and compost properties, while PERMANOVA was applied to evaluate beta diversity and compost properties. $P$-values were adjusted ($p$-adj) for multiple testing for each community metric separately using the Benjamini & Hochberg correction.

| | | **Richness** | | | **Shannon Evenness** | | | **Shannon Diversity** | | | **Community Structure** | | | |
|---|---|---|---|---|---|---|---|---|---|---|---|---|---|---|
| **Bacteria** | | $\rho$ | $p$ | $p$-adj | $\rho$ | $p$ | $p$-adj | $\rho$ | $p$ | $p$-adj | $R^2$ [%] | F | $p$ | $p$-adj |
| **Compost properties** | Compost age | 0.48 | ** | * | 0.13 | ns | ns | 0.30 | . | ns | 6.2 | 2.18 | * | * |
| | Dry substance | -0.35 | * | ns | -0.34 | * | ns | -0.34 | * | ns | 6.8 | 2.54 | ** | * |
| | WHC | 0.23 | ns | ns | 0.30 | . | ns | 0.27 | ns | ns | 6.4 | 2.41 | ** | * |
| | pH | -0.03 | ns | ns | 0.20 | ns | ns | 0.11 | ns | ns | 4.6 | 1.71 | . | . |
| | Salinity | -0.14 | ns | ns | 0.03 | ns | ns | -0.04 | ns | ns | 7.5 | 2.84 | ** | * |
| | $OD_{550}$ | -0.25 | ns | ns | 0.01 | ns | ns | -0.10 | ns | ns | 12.2 | 4.87 | ** | ** |
| | $NO_2^-$-N | -0.15 | ns | ns | -0.41 | * | ns | -0.25 | ns | ns | 5.2 | 1.93 | . | . |
| | $NO_3^-$-N | 0.35 | * | ns | 0.09 | ns | ns | 0.25 | ns | ns | 6.1 | 2.28 | * | * |
| | $NH_4^+$-N | -0.43 | ** | . | -0.29 | . | ns | -0.35 | * | ns | 5.2 | 1.91 | . | . |
| | $N_{min}$ | 0.21 | ns | ns | 0.11 | ns | ns | 0.21 | ns | ns | 4.3 | 1.57 | . | . |
| | $NO_3^-$-N/$N_{min}$ | 0.47 | ** | * | 0.20 | ns | ns | 0.33 | * | ns | 8.3 | 3.16 | ** | ** |
| | $PO_4^{3-}$-P | -0.03 | ns | ns | 0.17 | ns | ns | 0.07 | ns | ns | 11.3 | 4.46 | ** | ** |
| | $N_{tot}$ | 0.15 | ns | ns | 0.25 | ns | ns | 0.21 | ns | ns | 6.1 | 2.27 | ** | * |
| | $C_{org}$ | 0.07 | ns | ns | 0.16 | ns | ns | 0.10 | ns | ns | 5.9 | 2.18 | * | * |
| | $C_{org}/N_{tot}$ | -0.17 | ns | ns | -0.14 | ns | ns | -0.19 | ns | ns | 5.9 | 2.19 | ** | * |
| **Microbial activity** | Basal respiration | 0.07 | ns | ns | 0.18 | ns | ns | 0.15 | ns | ns | 7.4 | 2.56 | ** | * |
| | FDA hydrolysis | 0.07 | ns | ns | 0.34 | * | ns | 0.22 | ns | ns | 8.6 | 3.28 | ** | ** |
| **Fungi** | | | | | | | | | | | | | | |
| **Compost properties** | Compost age | 0.38 | * | ns | 0.41 | * | ns | 0.42 | * | ns | 8.8 | 3.19 | ** | * |
| | Dry substance | -0.11 | ns | ns | -0.01 | ns | ns | -0.03 | ns | ns | 7.4 | 2.79 | ** | * |
| | WHC | 0.14 | ns | ns | 0.06 | ns | ns | 0.09 | ns | ns | 4.6 | 1.69 | ns | ns |
| | pH | 0.13 | ns | ns | 0.03 | ns | ns | 0.09 | ns | ns | 4.3 | 1.56 | ns | ns |
| | Salinity | -0.17 | ns | ns | 0.03 | ns | ns | 0.00 | ns | ns | 10.2 | 3.97 | ** | * |
| | $OD_{550}$ | -0.38 | * | ns | -0.21 | ns | ns | -0.28 | . | ns | 5.2 | 1.94 | . | . |
| | $NO_2^-$-N | -0.23 | ns | ns | -0.40 | * | ns | -0.39 | * | ns | 7.5 | 2.84 | ** | * |
| | $NO_3^-$-N | 0.33 | * | ns | 0.31 | . | ns | 0.32 | . | ns | 9.4 | 3.63 | ** | * |
| | $NH_4^+$-N | -0.24 | ns | ns | -0.21 | ns | ns | -0.21 | ns | ns | 1.3 | 0.47 | ns | ns |
| | $N_{min}$ | 0.37 | * | ns | 0.24 | ns | ns | 0.27 | ns | ns | 7.0 | 2.62 | * | * |
| | $NO_3^-/N_{min}$ | 0.40 | * | ns | 0.37 | * | ns | 0.37 | * | ns | 4.4 | 1.62 | ns | ns |
| | $PO_4^{3-}$-P | -0.12 | ns | ns | 0.08 | ns | ns | 0.02 | ns | ns | 8.1 | 3.07 | ** | * |
| | $N_{tot}$ | 0.01 | ns | ns | 0.18 | ns | ns | 0.14 | ns | ns | 10.5 | 4.12 | ** | * |
| | $C_{org}$ | -0.12 | ns | ns | 0.00 | ns | ns | -0.04 | ns | ns | 7.2 | 2.70 | * | * |
| | $C_{org}/N_{tot}$ | -0.23 | ns | ns | -0.27 | ns | ns | -0.27 | ns | ns | 6.4 | 2.41 | * | * |
| **Microbial activity** | Basal respiration | -0.12 | ns | ns | -0.21 | ns | ns | -0.21 | ns | ns | 2.5 | 0.81 | ns | ns |
| | FDA hydrolysis | 0.09 | ns | ns | 0.17 | ns | ns | 0.15 | ns | ns | 5.8 | 2.15 | * | * |

Units are provided in Table S9. Significance levels are denoted as follows: ns = not significant, "." $p$ <0.1, * $p$ <0.05, ** $p$ <0.01, *** $p$ <0.001.

Table S9: **Relationship between compost sets and composting sites with physicochemical properties, microbial activities, and alpha- and beta-diversity of bacterial and fungal communities.** The analysis included all 37 composts for compost set comparisons and 35 composts collected from composting sites A–F for site-specific comparisons. Three data points were missing for basal respiration (10, 20, 29), and two were missing for compost age (31, 32). Statistics: A Kruskal-Wallis test was used for univariate analysis, while PERMANOVA was applied for multivariate analysis. $P$-values were adjusted ($p$-adj) for multiple testing separately for physicochemical properties/microbial activities and alpha diversity metrics.

| | | **Compost set** | | | **Composting site** | | |
|---|---|---|---|---|---|---|---|
| | | **Chi-sq** | **p** | **p-adj** | **Chi-sq** | **p** | **p-adj** |
| **Compost properties** | Compost age [days] | 11.22 | * | . | 9.21 | . | . |
| | DS [%] | 14.63 | ** | * | 15.67 | ** | ** |
| | Max. WHC [a] | 9.68 | * | . | 16.1 | ** | ** |
| | pH | 3.51 | ns | ns | 8.77 | . | . |
| | Salinity [b] | 3.58 | ns | ns | 12.33 | * | * |
| | $OD_{550}$ | 8.28 | * | ns | 9.66 | * | . |
| | $NO_2^-$-N [c] | 6.21 | ns | ns | 1.04 | ns | ns |
| | $NO_3^-$-N [c] | 3.65 | ns | ns | 6.36 | ns | ns |
| | $NH_4^+$-N [c] | 10.7 | * | . | 0.36 | ns | ns |
| | $N_{min}$ [c] | 0.37 | ns | ns | 3.38 | ns | ns |
| | $NO_3^-/N_{min}$ | 12.13 | ** | . | 4.09 | ns | ns |
| | $PO_4^{3-}$-P [c] | 2.81 | ns | ns | 22.63 | *** | *** |
| | $N_{tot}$ [%] | 2.36 | ns | ns | 28.23 | *** | *** |
| | $C_{org}$ [%] | 3.55 | ns | ns | 26.98 | *** | *** |
| | $C_{org}/N_{tot}$ | 5.7 | ns | ns | 14.05 | ** | * |
| **Microbial activity** | Basal respiration [d] | 4.89 | ns | ns | 2.34 | ns | ns |
| | FDA hydrolysis [e] | 4.95 | ns | ns | 15.52 | ** | * |
| **Alpha diversity** | Bacterial richness | 15.20 | ** | ** | 6.68 | ns | ns |
| | Bacterial Shannon evenness | 5.53 | ns | ns | 7.16 | ns | ns |
| | Bacterial Shannon diversity | 10.66 | * | * | 8.21 | ns | ns |
| | Fungal richness | 10.82 | * | * | 11.20 | * | . |
| | Fungal Shannon evenness | 5.90 | ns | ns | 8.77 | . | . |
| | Fungal Shannon diversity | 6.87 | . | ns | 9.98 | * | . |
| | | **F ($R^2$)** | ***p*** | ***p*-adj** | **F ($R^2$)** | ***p*** | ***p*-adj** |
| **Multivariate analysis** | PCA physicochemical properties | 2.01 (15.4%) | ** | NA | 3.68 (42.4%) | *** | NA |
| | Bacteria community structure | 1.55 (12.3%) | * | NA | 2.72 (26.6%) | *** | NA |
| | Fungi community structure | 1.80 (14.0%) | * | NA | 2.71 (26.5%) | *** | NA |

ns = not significant, "." $p$ <0.1, "*" $p$ <0.05, "**" $p$ <0.01, "***" $p$ <0.001,

[a] [g $H_2$0 (g DS)$^{-1}$] , [b] [g $KCl_{eq}$ (kg DS)$^{-1}$], $KCl_{eq}$ = Potassium chloride-equivalent, [c] [mg (kg DS)$^{-1}$] , [d] [mg $CO_2$-C h$^{-1}$], [e] [$\mu$g FDA (min g DS)$^{-1}$]

Table S10: **Taxonomic classification of indicative ASVs.** The table provides the number of distinct classifications (N) and the percentage of ASVs classified (%) at the phylum, family, and genus levels. Additionally, it highlights the most prevalent ASV classifications at each level, specifying the number of indicative ASVs assigned to these classifications.

| **plant-path system** | **Phylum** | | | **Family** | | | **Genus** | | |
|---|---|---|---|---|---|---|---|---|---|
| | **N (%)** | | **Classification** | **N (%)** | | **Classification** | **N (%)** | | **Classification** |
| *G. ultimum*-cress | 12 (96%) | 49 | *Pseudomonadota* | 53 (71%) | 9 | *Rhizobiaceae* | 48 (23%) | 3 | *Synoicihabitans* |
| | | 18 | *Chloroflexota* | | 8 | *Cyclobacteriaceae* | | 3 | *Devosia* |
| | | 18 | *Bacteroidota* | | 6 | *Opitutaceae* | | 2 | *Microvirga, Cellvirbio, Flavobacterium* |
| *G. ultimum*-cucumber | 14 (96%) | 54 | *Pseudomonadota* | 53 (91%) | 10 | *Sphingomonadaceae* | 50 (37%) | 4 | *Flavobacterium* |
| | | 26 | *Bacteroidota* | | 8 | *Cyclobacteriaceae* | | 3 | *W-Chloroflexi-9, VFJN01, Streptomyces, Luteimonas_D* |
| | | 17 | *Actinomycetota* | | 6 | *Xanthomonadaceae* | | | |
| *R. solani*-cucumber | 12 (95%) | 21 | *Pseudomonadota* | 35 (70%) | 6 | *Streptosproangiacaeae* | 27 (35%) | 4 | *Aggregatillinea* |
| | | 20 | *Actinomycetota* | | 4 | *Steroidobacteraceae Thermomicrobiaceae, Aggregatilineaceae* | | 3 | *Sphearobacter, Nonomuraea* |
| | | 13 | *Chloroflexota* | | | | | | |

Table S11: **Indicative ASVs which had a sequence match of at least 99% with one of the 75 indicative ASVs in Mayerhofer et al. (2021)**. In this study, ASVs were classified using the GTDB in this study. In our previous project, sequences were also clustered into ASVS but referred to as ESV (exact sequence variant) and classified with the SILVA database.

| **ASV** | **ESV** | **plant-path** | **match [%]** | **GTDB phylum** | **GTDB lowest** | **SILVA lowest** |
|---|---|---|---|---|---|---|
| 87 | 1266 | cr-Gu | 99.3 | *Pseudomonadota* | f *Steroidobacteraceae* | g *Steroidobacter* |
| 1113 | 1266 | cr-Gu | 99.1 | *Pseudomonadota* | s *Povalibacter uvarum* | g *Steroidobacter* |
| 652 | 250 | cr-Gu | 99.3 | *Pseudomonadota* | g *Devosia* | g *Devosia* |
| 1431 | 3619 | cr-Gu | 99.6 | *Chloroflexota* | f UBA6265 | f JG30-KF-CM45 |
| 1710 | 3867 | cr-Gu | 99.6 | *Pseudomonadota* | f *Rhizobiaceae* | o *Rhizobiales* |
| 146 | 162 | cuc-Gu | 100.0 | *Pseudomonadota* | g *Sphingopyxis* | g *Sphingopyxis* |
| 1823 | 11259 | cuc-Gu | 99.5 | *Pseudomonadota* | o *Pseudomonadales* | c *Gammaproteobacteria* |
| 3139 | 11259 | cuc-Gu | 99.8 | *Pseudomonadota* | o *Pseudomonadales* | c *Gammaproteobacteria* |
| 2470 | 7931 | cuc-Gu | 100.0 | *Pseudomonadota* | f SG8-39 | o *Betaproteobacteriales* |
| 467 | 64 | Gu | 99.8 | *Bacteroidota* | s *Algoriphagus terrigena* | g *Algoriphagus* |
| 2708 | 2082 | Gu | 99.3 | *Pseudomonadota* | o *Rhizobiales* | g Pseudorhodoplanes |
| 269 | 48 | cuc-Rs | 99.0 | *Pseudomonadota* | o *Rhizobiales* | o *Rhizobiales* |

cr-Gu = cress-*G. ultimum*, cuc-Gu = cucumber-*G. ultimum*, cuc-Rs = cucumber-*R. solani*,
c = class, o = order, f = family, g = genus, s = species

Table S12: **Effect of compost age and disease suppression in cress-*G. ultimum* combined on bacterial community structure of the 17 composts published by Mayerhofer et al. (2021)**. Compost age was only available for 12 of the 17 composts. Bacterial community structure was based on Bray-Curtis dissimilarities using mean relative abundance of ASVs. Dissimilarity based linear models were performed with PERMANOVA.

| **Factor** | **Df** | **explained variability** | **Pseudo-F** | ***p*-value** |
|---|---|---|---|---|
| age | 1 | 17.6 | 2.1 | 0.038 |
| Disease suppression | 1 | 7.5 | 0.9 | 0.524 |
| Residuals | 9 | 75 | | |
| Total | 11 | 100 | | |

Table S13: **Genera assigned to indicative ASVs for which their presence was previously reported in composts**

| cr-Gu | cuc-Gu | cuc-Rs | Genus | Literature on presence in compost |
|---|---|---|---|---|
| | ASV864 | | *Advenella* | Isolated from multiple waste compost (Tondello et al., 2022) |
| | | ASV4<br>ASV3067<br>ASV23055<br>ASV23950 | *Aggregatilinea* | (Fan et al., 2024) |
| ASV3229 | | | *Agrobacterium* | Isolate from paper sludge compost (Charest et al., 2005) |
| ASV467 | ASV467 | | *Algoriphagus* | Rice straw compost (Gavande et al., 2021), indicator maturation phase (Tortosa et al., 2021) |
| | | ASV298 | *Anseongella* | (Ji et al., 2023) |
| | ASV90 | | *Arenimonas* | Isolated from compost (Jin et al., 2012) |
| ASV1927 | ASV1927 | | *Brevundimonas* | Detected in composts (Antoniou et al., 2017), denitrifying bacteria (Sun et al., 2019) |
| | | ASV2570 | *Caldicoprobacter* | (Che et al., 2021) |
| | | ASV2729 | *Calditerricola* | Isolated from compost (Moriya et al., 2011) |
| ASV2571 | | | *Cellulosimicrobium* | (Hu et al., 2021a) |
| ASV231<br>ASV778 | ASV778 | | *Cellvibrio* | Indicator cooling phase (van Dijk et al., 2023) maturation phase (Li et al., 2020) |
| | | ASV298 | *Coprinellus* (fungi) | (Kurakov and Bilanenko, 2023) |
| ASV840 | | | *Croceibacterium* | (Gao et al., 2024) |
| ASV325<br>ASV399<br>ASV652 | ASV2820 | | *Devosia* | Indicator for thermophilic stage (Li et al., 2020), maturation phase (Blaya et al., 2016) |
| | ASV632<br>ASV4877 | | *Enhygromyxa* | Sheep manure composting (Cai et al., 2023) |
| | ASV587 | | *Ferrovibrio* | Isolated from compost (Wang et al., 2022b) |
| | | ASV710 | *Filomicrobium* | Isolated from compost (Li et al., 2024) |
| ASV780<br>ASV975 | ASV65<br>ASV561<br>ASV780 | | *Flavobacterium* | Several strains isolated from composts (Kim et al., 2012) |
| ASV739 | | | *Glutamicibacter* | Isolated from compost (Borker et al., 2021) |
| | ASV1137 | | *Glycomyces* | Indicator for cooling phase (Zhang et al., 2021) |
| | | ASV1249 | *Hyphomicrobium* | Detected in composts (Yang et al., 2023) |
| | ASV9 | | *Kernia (fungi)* | (Zhao et al., 2023) |
| ASV2767<br>ASV903 | ASV58<br>ASV849<br>ASV2767 | | *Luteimonas* | Isolated from compost (Young et al., 2007), humification processes (Tortosa et al., 2021) |
| | ASV752 | | *Membranicola* | (Lu et al., 2022) |
| | ASV1598 | | *Methylosinus* | (Wei et al., 2016) |
| ASV1379<br>ASV1408 | | | *Microvirga* | Abundant in thermophilic stage, nitrogen fixer (Parab et al., 2023) |
| | ASV122 | | *Mortierella (fungi)* | Indicator for compost maturation (Kurakov and Bilanenko, 2023) |

*Continued on next page*

| cr-Gu | cuc-Gu | cuc-Rs | Genus | Literature on presence in compost |
|---|---|---|---|---|
| | | ASV243 | *Mycobacterium* | Isolated from compost (Lima-Junior et al., 2016) |
| ASV500 | ASV500 | | *Nitrosomonas* | Indicator for maturation phase, nitrifier (Wang et al., 2022a) |
| | | ASV178<br>ASV658<br>ASV1067 | *Nonomuraea* | (Wu and Liu, 2016) |
| | ASV438 | | *Paracoccus* | (Tortosa et al., 2021) |
| | ASV1339<br>ASV1583 | | *Parapedobacter* | Isolated from compost (Kim et al., 2010) |
| ASV1781 | | | *Phenylobacterium* | Isolated from compost (Weon et al., 2008) |
| ASV1113 | | ASV794 | *Povalibacter* | Indicator for cooling phase (Zhong et al., 2020) |
| | ASV406 | | *Pseudorhizobium* | (Cai et al., 2023) |
| | ASV327 | | *Pseudoxanthomonas* | Isolated from compost (Lin et al., 2019) |
| ASV2260 | | | *Pusillimonas* | (Zainudin et al., 2020) |
| | | ASV434 | *Rhodomicrobium* | (Hu et al., 2021b) |
| ASV1700 | | | *Rhizomicrobium* | Dominant in maturation phase (Bohrer et al., 2023) |
| | | ASV76 | *Scedosporium* (fungi) | Indicator for maturation phase (Gu et al., 2017) |
| ASV1678 | | | *Shinella* | Isolated from compost (Vaz-Moreira et al., 2010) |
| | | ASV10<br>ASV3223<br>ASV8464 | *Sphaerobacter* | (Storey et al., 2015; Ren et al., 2023) |
| | ASV280 | | *Sphingobacterium* | Isolated from compost (Kim et al., 2006) |
| | ASV1563 | | *Sphingobium* | Isolate from compost (Vaz-Moreira et al., 2009) |
| ASV146 | | | *Sphingopyxis* | Indicator for cooling phase (Tian et al., 2013) |
| | ASV452<br>ASV737<br>ASV1749 | ASV791 | *Streptomyces* | Isolated from compost (Shen et al., 2021; Salamoni et al., 2010) |
| | | ASV102<br>ASV105 | *Thermomonospora* | Isolated from compost (Wu et al., 2019) |

cr-Gu: cress-*G. ultimum*, cuc-Gu: cucumber-*G. ultimum*, cuc-RS: cucumber-*R. solani*

Table S14: **Genera assigned to indicative ASVs for potential involvement in disease suppression was previously reported.**

| cr-Gu | cuc-Gu | cuc-Rs | Genus | Literature search results |
|---|---|---|---|---|
| | ASV864 | | *Advenella* | Low abundant member of suppressive syncom against root rot (Li et al., 2021) |
| ASV3229 | | | *Agrobacterium* | Control of crown call (Ryder et al., 1991), disease-suppressive compost isolate (Charest et al., 2005) |
| ASV467 | ASV467 | | *Algoriphagus* | Enriched in rhizosphere suppressive of bacterial wilt (Zheng et al., 2024) |
| | ASV90 | | *Arenimonas* | Indicator for suppressive soils (Ou et al., 2019) |
| ASV1927 | ASV1927 | | *Brevundimonas* | Control of *Fusarium redolens* in chickpea (Bekkar and Zaim, 2024) |
| ASV231<br>ASV778 | ASV778 | | *Cellvibrio* | Associated with bacterial wilt decrease (Messiha et al., 2023) |
| ASV427 | | | *Chthoniobacter* | Consortia suppressive of *Fusarium oxysporum* (Kudjordjie et al., 2023) |
| ASV325<br>ASV399<br>ASV652 | ASV2820 | | *Devosia* | Responsible for suppressive conditions in rhizosphere (Liu et al., 2024) |
| ASV780<br>ASV975 | ASV65<br>ASV561<br>ASV780 | | *Flavobacterium* | Part of consortium suppression of *R. solani* (Carrión et al., 2019) |
| ASV739 | | | *Glutamicibacter* | Isolate from compost with plant beneficial properties (Borker et al., 2021) |
| | ASV1137 | | *Glycomyces* | Antifungal properties (Ting et al., 2010) |
| | | ASV1249 | *Hyphomicrobium* | Enriched under suppressive conditions (Cao et al., 2024) |
| ASV2767<br>ASV903 | ASV58<br>ASV849<br>ASV2767 | | *Luteimonas* | Enriched in suppressive composts (Scotti et al., 2020; Hernández-Lara et al., 2022) |
| ASV1379<br>ASV1408 | | | *Microvirga* | Enriched in *Fusarium* wilt suppressive soils (Siegel-Hertz et al., 2018) |
| | ASV122 | | *Mortierella* (Fungi) | Associated with soils suppressive to *Fusarium* wilt (Khatri et al., 2023; De Corato et al., 2019; Xiong et al., 2017), enriched in soil suppressive of *G. ultimum* (Kurm et al., 2023) |
| | | ASV243 | *Mycobacterium* | Enriched in community suppressive of *Pythium aphanidermatum* in cucumber (Postma et al., 2005), production of siderophores (Meena et al., 2017) |
| ASV500 | ASV500 | | *Nitrosomonas* | Enriched under suppressive conditions (Shen et al., 2019) |
| | ASV3060 | | *Novosphingobium* | Enriched under suppressive conditions (Deng et al., 2022) |
| ASV1781 | | | *Phenylobacterium* | Associated with bacterial wilt control (Ding et al., 2023) |
| ASV1113 | | ASV794 | *Povalibacter* | Enriched in *Fusarium oxysporum* suppressive soils (Ou et al., 2019) |
| | ASV327 | | *Pseudoxanthomonas* | Significantly associated with *Fusarium* wilt incidence (Wen et al., 2023) |

*Continued on next page*

Table 3 **Bacterial genera and a fungal genus (continued)**

| cressGu | cucGu | cucRs | Genus | Disease Suppression |
|---|---|---|---|---|
| ASV1700 | | | *Rhizomicrobium* | Enriched in soil suppressive of *Fusarium* wilt (Lv et al., 2023) |
| ASV1678 | | | *Shinella* | Enriched by biochar application leading to *Fusarium* suppression (Jaiswal et al., 2017) |
| ASV1513 | ASV1513 | ASV3191 | *Sinorhizobium* | Enriched under suppressive conditions (Xue et al., 2015) |
| | ASV280 | | *Sphingobacterium* | Suppression of fungal pathogens (Xu et al., 2020), suppression of *Fusarium* (Matsuda et al., 2001) |
| | ASV1563 | | *Sphingobium* | Associated with bacterial wilt control (Ding et al., 2023) |
| | ASV146 | | *Sphingopyxis* | Antagonist of *Verticillium* pathogen (Snelders et al., 2020), part of a suppressive consortium (Fujiwara et al., 2016), enriched on healthy plants (Gu et al., 2022) |
| | ASV452 ASV737 ASV1749 | ASV791 | *Streptomyces* | Suppresses tomato wilt (Shen et al., 2021; Salamoni et al., 2010), antimicrobial properties (Kinkel et al., 2012), suppression of *R. solani* in pepper plants (Wang et al., 2015) |
| ASV2839 | | | *Telluria* | Potential root-knot nematode control (Oka, 2010) |

cr-Gu = cress-*G. ultimum*, cuc-Gu = *G. ultimum*-cucumber, cuc-Rs = cucumber-*R. solani*

Table S15: **Fungal ASVs indicative of the nine most suppressive composts** compared to the nine least suppressive. ASVs were ordered by increasing ranksum for each plant-pathogen system.

| **System** | **ASV** | **PBC** | **ISA** | **MAS** | **ALX** | **Rank** [a] | **n**[b] | $\rho$ | $\boldsymbol{p}$ | **Phylum** | **Lowest taxonomy** |
|---|---|---|---|---|---|---|---|---|---|---|---|
| cr-Gu | 116 | 0.4 | 0.87 | 0.022 | NA | 5 | 22 | 0.34 | 0.037 | *Ascomycota* | *f Pezizales IS* |
| cr-Gu | 127 | 0.44 | 0.88 | 0.022 | NA | 3 | 20 | 0.42 | 0.009 | *Ascomycota* | *f Ascodesmidaceae* |
| cr-Gu | 185 | 0.31 | 0.81 | 0.014 | NA | 9 | 17 | 0.32 | 0.055 | *Ascomycota* | *f Pezizales IS* |
| cuc-Gu | 122 | 0.41 | 0.88 | 0.053 | 0.85 | 7 | 16 | 0.50 | 0.002 | *Mortierellomycota* | *s Mortierella yunnanensis* |
| cuc-Gu | 9 | NA | 0.94 | 0.115 | 0.64 | 11 | 34 | 0.37 | 0.025 | *Ascomycota* | *g Kernia* |
| cuc-Gu | 29 | 0.37 | 0.95 | 0.048 | NA | 11 | 29 | 0.39 | 0.016 | *Basidiomycota* | *g Coprinellus* |
| cuc-Gu | 45 | 0.37 | 0.74 | NA | 0.68 | 13 | 17 | 0.31 | 0.060 | *Ascomycota* | *p Ascomycota* |
| cuc-Gu | 76 | 0.36 | 0.81 | NA | 0.61 | 16 | 14 | 0.41 | 0.013 | *Ascomycota* | *s Scedosporium prolificans* |
| cuc-Rs | 64 | 0.48 | 0.88 | 0.06 | NA | 1 | 24 | 0.43 | 0.007 | *Ascomycota* | *o Microascales* |

[a] Rank sum: indicative ASVs were ranked based on the effect size for all four statistical methods (PBC, ISA, MAS, ALX) and then the ranks were summed up. [b] Number of composts the ASVs was detected. PBC = Point Biserial Correlation, ISA = Indicator Species Analysis, MAS = MaAsLins, ALX = ALDEx2. cu-Gu = cress-*G. ultimum*, cuc-Gu = cucumber-*G. ultimum*, cuc-Rs = cucumber-*R. solani*.

Table S16: **Bacterial ASVs indicative of the nine most suppressive composts in the cress-*G. ultimum* system.** For abbreviations see Table S15

| ASV | PBC | ISA | MAS | ALX | Rank | n | rho | p | Phylum | Lowest taxonomy |
|---|---|---|---|---|---|---|---|---|---|---|
| 618 | 0.6 | 0.92 | 0.017 | NA | 62 | 32 | 0.48 | 0.003 | *Planctomycetota* | *f Lacipirellulaceae* |
| 594 | 0.69 | 0.93 | 0.012 | NA | 71 | 29 | 0.45 | 0.006 | *Planctomycetota* | *f Tepidisphaeraceae* |
| 242 | 0.56 | 0.91 | 0.022 | NA | 84 | 30 | 0.50 | 0.002 | *Acidobacteriota* | *o Vicinamibacterales* |
| 622 | 0.57 | 0.92 | 0.013 | NA | 87 | 28 | 0.55 | 0.000 | *Gemmatimonadota* | *g Palsa-1233* |
| 198 | 0.56 | 0.91 | 0.019 | NA | 88 | 29 | 0.45 | 0.005 | unclassified | *unclassified* |
| 543 | 0.53 | 0.92 | 0.018 | 0.59 | 89 | 22 | 0.54 | 0.001 | unclassified | *unclassified* |
| 135 | 0.56 | 0.9 | 0.024 | NA | 92 | 30 | 0.47 | 0.003 | *Acidobacteriota* | *g OLB17* |
| 922 | 0.64 | 0.92 | 0.01 | NA | 93 | 23 | 0.42 | 0.009 | *Pseudomonadota* | *g Sphingomicrobium* |
| 1257 | 0.56 | 0.94 | 0.01 | NA | 101 | 20 | 0.58 | 0.000 | *Planctomycetota* | *o Phycisphaerales* |
| 94 | 0.53 | 0.91 | 0.03 | NA | 102 | 33 | 0.47 | 0.004 | *Chloroflexota* | *f Phototrophicaceae* |
| 1927 | 0.53 | 0.96 | 0.01 | 0.69 | 104 | 28 | 0.50 | 0.002 | *Pseudomonadota* | *s Brevundimonas basaltis* |
| 87 | 0.5 | 0.92 | 0.028 | NA | 105 | 36 | 0.38 | 0.022 | *Pseudomonadota* | *f Steroidobacteraceae* |
| 2750 | 0.6 | 0.92 | 0.009 | NA | 113 | 25 | 0.50 | 0.002 | unclassified | unclassified |
| 174 | 0.49 | 0.92 | 0.023 | NA | 114 | 29 | 0.41 | 0.012 | unclassified | unclassified |
| 824 | 0.49 | 0.92 | 0.015 | 0.55 | 120 | 31 | 0.53 | 0.001 | *Pseudomonadota* | *f Rhizobiaceae* |
| 702 | 0.6 | 0.88 | 0.013 | NA | 122 | 32 | 0.37 | 0.024 | *Bacteroidota* | *f Cyclobacteriaceae* |
| 698 | 0.65 | 0.87 | 0.013 | NA | 124 | 32 | 0.47 | 0.003 | *Planctomycetota* | *o Pirellulales* |
| 1379 | 0.52 | 0.95 | 0.01 | NA | 126 | 31 | 0.43 | 0.008 | *Pseudomonadota* | *g Microvirga* |
| 214 | 0.48 | 0.93 | 0.015 | NA | 128 | 29 | 0.54 | 0.001 | *Patescibacteria* | s JACDBH01 sp013694995 |
| 689 | 0.52 | 0.91 | 0.014 | 0.5 | 129 | 30 | 0.44 | 0.007 | *Chloroflexota* | *f Caldilineaceae* |
| 124 | 0.55 | 0.87 | 0.026 | NA | 131 | 34 | 0.43 | 0.009 | *Bacteroidota* | s UBA2336 sp002425185 |
| 295 | 0.53 | 0.89 | 0.017 | NA | 133 | 32 | 0.49 | 0.002 | *Pseudomonadota* | *c Alphaproteobacteria* |
| 107 | 0.56 | 0.86 | 0.02 | NA | 134 | 35 | 0.30 | 0.071 | *Pseudomonadota* | s SYSU-D60015 sp003576705 |
| 500 | 0.5 | 0.91 | 0.016 | NA | 135 | 34 | 0.41 | 0.011 | *Pseudomonadota* | *s Nitrosomonas nitrosa* |
| 519 | 0.54 | 0.9 | 0.013 | NA | 138 | 29 | 0.32 | 0.052 | *Bacteroidota* | *f Cyclobacteriaceae* |
| 851 | 0.54 | 0.9 | 0.013 | NA | 138 | 30 | 0.41 | 0.012 | *Chloroflexota* | *f UBA2979* |
| 184 | 0.49 | 0.9 | 0.024 | NA | 138 | 32 | 0.44 | 0.006 | *Bacteroidota* | *f Flavobacteriaceae* |
| 234 | 0.55 | 0.87 | 0.019 | NA | 139 | 31 | 0.37 | 0.026 | *Pseudomonadota* | *o Burkholderiales* |
| 316 | 0.46 | 0.93 | 0.015 | NA | 139 | 23 | 0.45 | 0.006 | *Verrucomicrobiota* | *s Nibricoccus aquaticus* |
| 548 | 0.57 | 0.87 | 0.013 | NA | 141 | 32 | 0.39 | 0.016 | *Acidobacteriota* | f SCN-69-37 |
| 568 | 0.61 | 0.85 | 0.014 | 0.49 | 144 | 33 | 0.48 | 0.003 | *Planctomycetota* | *g Lacipirellula* |
| 998 | 0.56 | 0.89 | 0.01 | NA | 147 | 33 | 0.32 | 0.053 | *Acidobacteriota* | f SCN-69-37 |
| 382 | 0.44 | 0.95 | 0.014 | NA | 148 | 35 | 0.33 | 0.046 | *Pseudomonadota* | *f Devosiaceae* |
| 427 | 0.46 | 0.95 | 0.012 | NA | 149 | 30 | 0.40 | 0.015 | *Verrucomicrobiota* | *g Chthoniobacter* |
| 789 | 0.53 | 0.92 | 0.009 | NA | 151 | 23 | 0.45 | 0.006 | *Chloroflexota* | *p Chloroflexota* |
| 2571 | 0.6 | 0.91 | 0.007 | NA | 156 | 24 | 0.36 | 0.028 | *Actinomycetota* | *g Cellulosimicrobium* |
| 294 | 0.46 | 0.91 | 0.016 | NA | 158 | 31 | 0.38 | 0.021 | *Chloroflexota* | *f Caldilineaceae* |
| 396 | 0.55 | 0.86 | 0.015 | NA | 159 | 33 | 0.36 | 0.029 | *Pseudomonadota* | *o Burkholderiales* |
| 2552 | 0.55 | 0.92 | 0.007 | 0.54 | 159 | 19 | 0.40 | 0.015 | *Bacteroidota* | *c Bacteroidia* |
| 571 | 0.4 | 0.98 | 0.014 | NA | 161 | 26 | 0.37 | 0.024 | *Chloroflexota* | *f Roseiflexaceae* |
| 840 | 0.51 | 0.89 | 0.013 | NA | 163 | 29 | 0.38 | 0.019 | *Pseudomonadota* | *s Croceibacterium sp001897135* |
| 325 | 0.52 | 0.86 | 0.02 | NA | 164 | 25 | 0.41 | 0.012 | *Pseudomonadota* | *g Devosia* |

*Continued on next page*

| ASV | PBC | ISA | MAS | ALX | Rank | n | rho | p | Phylum | Lowest taxonomy |
|---|---|---|---|---|---|---|---|---|---|---|
| 1301 | 0.44 | 0.95 | 0.011 | NA | 166 | 30 | 0.44 | 0.007 | *Bacteroidota* | *c Bacteroidia* |
| 1678 | 0.46 | 0.95 | 0.009 | NA | 173 | 35 | 0.45 | 0.005 | *Pseudomonadota* | *s Shinella granuli* |
| 517 | 0.35 | 0.93 | 0.017 | NA | 173 | 31 | 0.43 | 0.007 | *Patescibacteria* | s J137 sp003694175 |
| 1566 | 0.43 | 0.94 | 0.011 | NA | 175 | 24 | 0.56 | 0.000 | *Pseudomonadota* | s SG8-41 sp023266245 |
| 739 | 0.45 | 0.94 | 0.01 | NA | 176 | 34 | 0.38 | 0.020 | *Actinomycetota* | *s Glutamicibacter nicotianae* |
| 1080 | 0.59 | 0.89 | 0.007 | NA | 178 | 20 | 0.47 | 0.004 | *Pseudomonadota* | *f Sphingomonadaceae* |
| 1431 | 0.54 | 0.89 | 0.009 | NA | 179 | 26 | 0.41 | 0.012 | *Chloroflexota* | f UBA6265 |
| 1113 | 0.35 | 0.93 | 0.015 | NA | 180 | 24 | 0.52 | 0.001 | *Pseudomonadota* | *s Povalibacter uvarum* |
| 268 | 0.47 | 0.88 | 0.016 | NA | 181 | 32 | 0.41 | 0.012 | *Bacteroidota* | *f Cyclobacteriaceae* |
| 467 | 0.36 | 0.92 | 0.016 | NA | 181 | 28 | 0.48 | 0.002 | *Bacteroidota* | *s Algoriphagus terrigena* |
| 735 | 0.38 | 0.93 | 0.013 | NA | 184 | 25 | 0.49 | 0.002 | *unclassified* | unclassified |
| 399 | 0.42 | 0.94 | 0.01 | NA | 185 | 32 | 0.43 | 0.009 | *Pseudomonadota* | *s Devosia sp024707125* |
| 832 | 0.51 | 0.88 | 0.011 | NA | 186 | 28 | 0.36 | 0.029 | *Pseudomonadota* | *c Gammaproteobacteria* |
| 1253 | 0.59 | 0.87 | 0.008 | NA | 186 | 19 | 0.44 | 0.006 | *Fibrobacterota* | *s Chersky-265 sp019509785* |
| 717 | 0.52 | 0.87 | 0.012 | NA | 187 | 32 | 0.40 | 0.014 | *Bacteroidota* | *f Cyclobacteriaceae* |
| 2247 | 0.61 | 0.89 | 0.006 | NA | 188 | 25 | 0.49 | 0.002 | *Pseudomonadota* | *o Rhizobiales* |
| 780 | 0.45 | 0.92 | 0.01 | NA | 189 | 20 | 0.38 | 0.020 | *Bacteroidota* | *s Flavobacterium marinum* |
| 891 | 0.5 | 0.88 | 0.011 | NA | 190 | 31 | 0.46 | 0.004 | *Planctomycetota* | *f Lacipirellulaceae* |
| 1144 | 0.32 | 0.92 | 0.014 | 0.54 | 190 | 20 | 0.50 | 0.002 | *Bacteroidota* | *f Cyclobacteriaceae* |
| 974 | 0.62 | 0.84 | 0.009 | 0.43 | 191 | 29 | 0.36 | 0.030 | *Chloroflexota* | s WHTK01 sp009377745 |
| 2202 | 0.65 | 0.88 | 0.006 | NA | 191 | 22 | 0.54 | 0.001 | *Planctomycetota* | s JADLFV01 sp020849625 |
| 498 | 0.46 | 0.91 | 0.01 | NA | 193 | 30 | 0.44 | 0.006 | *Patescibacteria* | *o Saccharimonadales* |
| 1058 | 0.62 | 0.85 | 0.008 | NA | 198 | 34 | 0.47 | 0.003 | *Actinomycetota* | *f Ilumatobacteraceae* |
| 1513 | 0.56 | 0.84 | 0.009 | 0.62 | 200 | 26 | 0.47 | 0.004 | *Pseudomonadota* | *s Sinorhizobium fredii* |
| 1777 | 0.52 | 0.89 | 0.008 | NA | 205 | 29 | 0.40 | 0.014 | *Bacteroidota* | *o Cytophagales* |
| 1729 | 0.56 | 0.86 | 0.008 | NA | 206 | 22 | 0.48 | 0.003 | *Planctomycetota* | *f Phycisphaeraceae* |
| 1341 | 0.43 | 0.92 | 0.009 | NA | 209 | 27 | 0.28 | 0.099 | *Pseudomonadota* | *f Rhizobiaceae* |
| 231 | 0.36 | 0.88 | 0.02 | NA | 215 | 25 | 0.49 | 0.002 | *Pseudomonadota* | *g Cellvibrio* |
| 955 | 0.58 | 0.82 | 0.009 | NA | 215 | 28 | 0.37 | 0.026 | *Pseudomonadota* | *f Burkholderiaceae B* |
| 2418 | 0.47 | 0.94 | 0.006 | NA | 217 | 28 | 0.32 | 0.055 | *Pseudomonadota* | *f Rhizobiaceae* |
| 778 | 0.34 | 0.91 | 0.013 | NA | 217 | 26 | 0.43 | 0.008 | *Pseudomonadota* | *g Cellvibrio* |
| 1488 | 0.42 | 0.93 | 0.008 | NA | 218 | 21 | 0.53 | 0.001 | *Bacteroidota* | *f Cyclobacteriaceae* |
| 1696 | 0.46 | 0.92 | 0.007 | NA | 219 | 20 | 0.46 | 0.004 | *Pseudomonadota* | *s Vitreimonas sp900696695* |
| 1432 | 0.49 | 0.83 | 0.013 | 0.67 | 220 | 17 | 0.24 | 0.161 | *Planctomycetota* | *f Planctomycetaceae* |
| 744 | 0.56 | 0.83 | 0.009 | NA | 221 | 32 | 0.35 | 0.033 | *Pseudomonadota* | f CACIAM-22H2 |
| 3229 | 0.55 | 0.9 | 0.005 | NA | 222 | 29 | 0.52 | 0.001 | *Pseudomonadota* | *s Agrobacterium salinitolerans* |
| 645 | 0.46 | 0.87 | 0.011 | NA | 224 | 30 | 0.30 | 0.070 | *Pseudomonadota* | s ZC4RG30 sp003242275 |
| 1966 | 0.56 | 0.87 | 0.006 | NA | 225 | 29 | 0.43 | 0.007 | *Chloroflexota* | *f Tepidiformaceae* |
| 1824 | 0.41 | 0.96 | 0.007 | NA | 226 | 24 | 0.47 | 0.003 | *Verrucomicrobiota* | *s Synoicihabitans sp023386835* |
| 2427 | 0.52 | 0.9 | 0.006 | NA | 227 | 27 | 0.50 | 0.002 | *Hydrogenedentota* | f SLHB01 |
| 2811 | 0.62 | 0.84 | 0.006 | 0.52 | 231 | 23 | 0.49 | 0.002 | *Pseudomonadota* | *f Sphingomonadaceae* |
| 975 | 0.48 | 0.86 | 0.01 | NA | 231 | 24 | 0.38 | 0.021 | *Bacteroidota* | *s Flavobacterium marinum* |
| 1145 | 0.57 | 0.84 | 0.007 | NA | 233 | 24 | 0.43 | 0.007 | *Pseudomonadota* | *o Burkholderiales* |
| 1534 | 0.46 | 0.89 | 0.008 | NA | 239 | 30 | 0.54 | 0.001 | *Chloroflexota* | s W-Chloroflexi-9 sp023228825 |
| 1392 | 0.5 | 0.85 | 0.009 | NA | 243 | 25 | 0.36 | 0.028 | *Acidobacteriota* | *g Gp6-AA40* |

*Continued on next page*

| ASV | PBC | ISA | MAS | ALX | Rank | n | rho | p | Phylum | Lowest taxonomy |
|---|---|---|---|---|---|---|---|---|---|---|
| 914 | 0.4 | 0.88 | 0.011 | NA | 243 | 31 | 0.36 | 0.029 | *Chloroflexota* | s QEUY01 sp003577365 |
| 1157 | 0.36 | 0.91 | 0.009 | NA | 244 | 27 | 0.41 | 0.011 | *Pseudomonadota* | *f Rhizobiaceae* |
| 1261 | 0.54 | 0.84 | 0.008 | NA | 245 | 31 | 0.43 | 0.009 | *Pseudomonadota* | *c Alphaproteobacteria* |
| 937 | NA | 0.89 | 0.01 | 0.64 | 246 | 24 | 0.31 | 0.063 | *Chloroflexota* | s UBA6265 sp023957015 |
| 1781 | 0.56 | 0.82 | 0.007 | NA | 251 | 27 | 0.44 | 0.007 | *Pseudomonadota* | *s Phenylobacterium sp024699985* |
| 260 | 0.35 | 0.93 | 0.007 | NA | 252 | 29 | 0.29 | 0.079 | *Actinomycetota* | *f Micromonosporaceae* |
| 3097 | 0.57 | 0.85 | 0.005 | NA | 253 | 30 | 0.45 | 0.005 | *Planctomycetota* | *o Pirellulales* |
| 1075 | 0.39 | 0.91 | 0.008 | NA | 255 | 27 | 0.27 | 0.103 | *Chloroflexota* | *f Chloroflexaceae* |
| 4088 | 0.64 | 0.84 | 0.004 | NA | 257 | 31 | 0.35 | 0.032 | *Chloroflexota* | s SHYQ01 sp903917615 |
| 2096 | NA | 0.9 | 0.009 | 0.53 | 258 | 18 | 0.29 | 0.078 | *Planctomycetota* | *f Planctomycetaceae* |
| 1277 | 0.55 | 0.85 | 0.006 | NA | 258 | 30 | 0.33 | 0.049 | *Verrucomicrobiota* | s Opi-474 sp903830425 |
| 2602 | 0.46 | 0.88 | 0.007 | 0.49 | 259 | 23 | 0.41 | 0.012 | *Bacteroidota* | *o Rhodothermales* |
| 1264 | 0.5 | 0.81 | 0.01 | NA | 262 | 22 | 0.41 | 0.011 | *Verrucomicrobiota* | *f Opitutaceae* |
| 1134 | 0.42 | 0.86 | 0.01 | NA | 262 | 25 | 0.45 | 0.005 | *Pseudomonadota* | *f Devosiaceae* |
| 403 | 0.4 | 0.86 | 0.011 | NA | 264 | 23 | 0.49 | 0.002 | *Bacteroidota* | *f Chitinophagaceae* |
| 592 | 0.42 | 0.87 | 0.009 | NA | 265 | 23 | 0.45 | 0.005 | *Pseudomonadota* | g CADEED01 |
| 4401 | 0.54 | 0.85 | 0.006 | NA | 265 | 17 | 0.50 | 0.002 | *Pseudomonadota* | *o Rhizobiales* |
| 3041 | 0.49 | 0.88 | 0.005 | NA | 273 | 24 | 0.41 | 0.012 | *Verrucomicrobiota* | *s Synoicihabitans sp023386835* |
| 826 | 0.38 | 0.89 | 0.008 | NA | 274 | 27 | 0.27 | 0.108 | *Bacteroidota* | *s Kaistella haifensis* |
| 2741 | 0.57 | 0.82 | 0.005 | NA | 274 | 31 | 0.39 | 0.018 | *Actinomycetota* | *f Ilumatobacteraceae* |
| 2488 | 0.57 | 0.82 | 0.005 | NA | 274 | 21 | 0.41 | 0.011 | *Actinomycetota* | *f Ilumatobacteraceae* |
| 2210 | 0.46 | 0.88 | 0.006 | NA | 278 | 27 | 0.32 | 0.051 | *Chloroflexota* | *p Chloroflexota* |
| 1408 | 0.4 | 0.88 | 0.008 | NA | 279 | 32 | 0.32 | 0.055 | *Pseudomonadota* | *g Microvirga* |
| 1198 | 0.55 | 0.85 | 0.004 | NA | 281 | 27 | 0.37 | 0.026 | *Gemmatimonadota* | *c Gemmatimonadetes* |
| 1046 | 0.48 | 0.85 | 0.007 | NA | 283 | 31 | 0.31 | 0.061 | unclassified | unclassified |
| 784 | 0.52 | 0.79 | 0.008 | NA | 284 | 22 | 0.33 | 0.047 | *Myxococcota* | *f Polyangiaceae* |
| 2193 | 0.47 | 0.84 | 0.008 | NA | 285 | 18 | 0.48 | 0.003 | *Pseudomonadota* | *f Rhizobiaceae* |
| 1291 | 0.41 | 0.87 | 0.008 | NA | 287 | 22 | 0.49 | 0.002 | *Bacteroidota* | *f Chitinophagaceae* |
| 1700 | 0.56 | 0.81 | 0.005 | NA | 291 | 20 | 0.37 | 0.024 | *Pseudomonadota* | *g Rhizomicrobium* |
| 652 | 0.37 | 0.86 | 0.009 | NA | 292 | 23 | 0.29 | 0.079 | *Pseudomonadota* | *g Devosia* |
| 2887 | 0.48 | 0.84 | 0.007 | NA | 292 | 20 | 0.38 | 0.020 | *Gemmatimonadota* | *o Longimicrobiales* |
| 2767 | 0.49 | 0.82 | 0.007 | NA | 297 | 22 | 0.37 | 0.025 | *Pseudomonadota* | *s Luteimonas D colneyensis* |
| 2134 | 0.46 | 0.86 | 0.006 | NA | 299 | 31 | 0.38 | 0.021 | *Actinomycetota* | s SHVJ01 sp016870355 |
| 2309 | 0.58 | 0.78 | 0.004 | NA | 300 | 25 | 0.29 | 0.086 | *Actinomycetota* | *o Acidimicrobiales* |
| 2238 | 0.42 | 0.86 | 0.007 | NA | 303 | 21 | 0.51 | 0.001 | *Pseudomonadota* | *c Gammaproteobacteria* |
| 1532 | 0.45 | 0.81 | 0.009 | NA | 309 | 18 | 0.48 | 0.003 | *Patescibacteria* | o UBA9983 A |
| 1710 | 0.47 | 0.82 | 0.007 | NA | 310 | 19 | 0.44 | 0.007 | *Pseudomonadota* | *f Rhizobiaceae* |
| 3127 | 0.53 | 0.82 | 0.004 | NA | 314 | 30 | 0.30 | 0.070 | *Chloroflexota* | *f Thermomicrobiaceae* |
| 4699 | 0.51 | 0.82 | 0.005 | NA | 317 | 24 | 0.39 | 0.016 | *Pseudomonadota* | s SCGC-AG-212-J23 sp005881595 |
| 903 | 0.35 | 0.81 | 0.011 | NA | 318 | 18 | 0.49 | 0.002 | *Pseudomonadota* | *s Luteimonas B weifangensis* |
| 2708 | 0.48 | 0.82 | 0.006 | NA | 320 | 27 | 0.41 | 0.011 | *Pseudomonadota* | *o Rhizobiales* |
| 1815 | 0.44 | 0.85 | 0.006 | NA | 324 | 25 | 0.33 | 0.046 | *Myxococcota* | s JABFXX01 sp005887925 |
| 3518 | 0.49 | 0.84 | 0.004 | NA | 324 | 28 | 0.43 | 0.008 | *Chloroflexota* | s JAIBBD01 sp019694855 |
| 4340 | 0.44 | 0.86 | 0.005 | NA | 326 | 25 | 0.40 | 0.014 | *Actinomycetota* | *f Solirubrobacteraceae* |
| 2839 | 0.51 | 0.82 | 0.004 | NA | 327 | 32 | 0.25 | 0.138 | *Pseudomonadota* | *g Telluria* |

*Continued on next page*

| **ASV** | **PBC** | **ISA** | **MAS** | **ALX** | **Rank** | **n** | **rho** | **p** | **Phylum** | **Lowest taxonomy** |
|---|---|---|---|---|---|---|---|---|---|---|
| 2366 | 0.48 | 0.82 | 0.005 | NA | 333 | 25 | 0.32 | 0.051 | *Patescibacteria* | *o Saccharimonadales* |
| 2329 | 0.47 | 0.77 | 0.006 | NA | 345 | 20 | 0.36 | 0.029 | *Acidobacteriota* | f SCN-69-37 |
| 2600 | 0.42 | 0.8 | 0.007 | NA | 350 | 22 | 0.28 | 0.098 | *Verrucomicrobiota* | *s Synoicihabitans sp023386835* |
| 1463 | 0.42 | 0.79 | 0.007 | NA | 351 | 24 | 0.37 | 0.025 | *Verrucomicrobiota* | f UBA11358 |
| 2100 | 0.41 | 0.84 | 0.005 | NA | 359 | 24 | 0.32 | 0.055 | *Chloroflexota* | *o Thermomicrobiales* |

Table S17: **Bacterial ASVs indicative of the nine most suppressive composts in the cucumber-*G. ultimum* system.** For abbreviations see Table S15

| ASV | PBC | ISA | MAS | ALX | Rank | n | rho | p | Phylum | Lowest taxonomy |
|---|---|---|---|---|---|---|---|---|---|---|
| 424 | 0.72 | 0.96 | 0.021 | 0.67 | 32 | 32 | 0.55 | 0.000 | *Pseudomonadota* | *f Sphingomonadaceae* |
| 661 | 0.73 | 0.99 | 0.017 | 0.53 | 38 | 30 | 0.62 | 0.000 | *Pseudomonadota* | *f Burkholderiaceae C* |
| 1339 | 0.64 | 0.99 | 0.015 | 0.92 | 39 | 21 | 0.61 | 0.000 | *Bacteroidota* | *g Parapedobacter* |
| 146 | 0.73 | 0.95 | 0.024 | NA | 59 | 35 | 0.50 | 0.002 | *Pseudomonadota* | *g Sphingopyxis* |
| 308 | 0.67 | 0.94 | 0.020 | 0.52 | 67 | 33 | 0.55 | 0.000 | *Pseudomonadota* | *f Xanthomonadaceae* |
| 1513 | 0.73 | 0.96 | 0.011 | 0.53 | 80 | 26 | 0.65 | 0.000 | *Pseudomonadota* | *s Sinorhizobium fredii* |
| 840 | 0.57 | 0.95 | 0.014 | 0.67 | 85 | 29 | 0.54 | 0.001 | *Pseudomonadota* | *s Croceibacterium sp001897135* |
| 184 | 0.54 | 0.98 | 0.026 | NA | 87 | 32 | 0.47 | 0.003 | *Bacteroidota* | *f Flavobacteriaceae* |
| 1432 | 0.57 | 0.92 | 0.016 | 0.84 | 94 | 17 | 0.50 | 0.002 | *Planctomycetota* | *f Planctomycetaceae* |
| 500 | 0.58 | 0.95 | 0.018 | NA | 96 | 34 | 0.47 | 0.003 | *Pseudomonadota* | *s Nitrosomonas nitrosa* |
| 58 | 0.53 | 0.94 | 0.041 | 0.52 | 100 | 37 | 0.45 | 0.005 | *Pseudomonadota* | *s Luteimonas D sp000472505* |
| 567 | 0.71 | 0.90 | 0.012 | 0.55 | 124 | 32 | 0.48 | 0.003 | *Bacteroidota* | *s Flavipsychrobacter sp020636495* |
| 384 | 0.60 | 0.91 | 0.015 | NA | 129 | 32 | 0.33 | 0.043 | *Sumerlaeota* | s JAHLLQ01 sp020444065 |
| 124 | 0.55 | 0.90 | 0.025 | NA | 138 | 34 | 0.39 | 0.017 | *Bacteroidota* | s UBA2336 sp002425185 |
| 3130 | 0.61 | 0.97 | 0.007 | NA | 145 | 24 | 0.52 | 0.001 | *Planctomycetota* | *c Planctomycetia* |
| 2100 | 0.63 | 0.96 | 0.007 | NA | 146 | 24 | 0.46 | 0.004 | *Chloroflexota* | *o Thermomicrobiales* |
| 1830 | 0.80 | 0.90 | 0.007 | 0.74 | 148 | 34 | 0.54 | 0.001 | *Gemmatimonadota* | *f UBA6960* |
| 252 | 0.51 | 0.93 | 0.017 | NA | 149 | 28 | 0.31 | 0.059 | *Bacteroidota* | *s Edaphocola sp019739295* |
| 2767 | 0.59 | 0.97 | 0.007 | NA | 151 | 22 | 0.49 | 0.002 | *Pseudomonadota* | *s Luteimonas D colneyensis* |
| 90 | 0.56 | 0.89 | 0.021 | NA | 151 | 36 | 0.36 | 0.027 | *Pseudomonadota* | *s Arenimonas fontis* |
| 538 | 0.55 | 0.92 | 0.011 | NA | 154 | 31 | 0.28 | 0.099 | *Pseudomonadota* | *f Steroidobacteraceae* |
| 632 | 0.62 | 0.89 | 0.013 | 0.49 | 156 | 32 | 0.41 | 0.012 | *Myxococcota* | *g Enhygromyxa* |
| 3408 | 0.71 | 0.92 | 0.007 | 0.48 | 157 | 22 | 0.49 | 0.002 | *Pseudomonadota* | *c Gammaproteobacteria* |
| 65 | 0.49 | 0.92 | 0.022 | NA | 158 | 33 | 0.41 | 0.012 | *Bacteroidota* | *s Flavobacterium marinum* |
| 1230 | 0.64 | 0.89 | 0.009 | 0.63 | 159 | 33 | 0.45 | 0.006 | *Gemmatimonadota* | *f UBA6960* |
| 749 | 0.41 | 0.98 | 0.019 | 0.54 | 160 | 26 | 0.36 | 0.029 | *Acidobacteriota* | *f SCN-69-37* |
| 455 | 0.63 | 0.85 | 0.014 | 0.81 | 162 | 33 | 0.36 | 0.030 | *Planctomycetota* | *f Planctomycetaceae* |
| 568 | 0.61 | 0.88 | 0.011 | 0.57 | 163 | 33 | 0.49 | 0.002 | *Planctomycetota* | *g Lacipirellula* |
| 1390 | 0.65 | 0.91 | 0.008 | NA | 166 | 30 | 0.42 | 0.011 | *Planctomycetota* | *f Lacipirellulaceae* |
| 702 | 0.58 | 0.90 | 0.011 | NA | 167 | 32 | 0.31 | 0.060 | *Bacteroidota* | *f Cyclobacteriaceae* |
| 1370 | 0.53 | 0.94 | 0.009 | NA | 168 | 30 | 0.48 | 0.003 | *Actinomycetota* | *s VFJN01 sp009694375* |
| 388 | 0.47 | 0.93 | 0.020 | NA | 170 | 26 | 0.44 | 0.007 | *Pseudomonadota* | s SYSU-D60014 sp003576685 |
| 569 | 0.60 | 0.87 | 0.011 | 0.60 | 173 | 33 | 0.38 | 0.021 | *Gemmatimonadota* | *c Gemmatimonadetes* |
| 457 | 0.39 | 0.98 | 0.020 | 0.49 | 176 | 31 | 0.44 | 0.007 | *Bacteroidota* | *g JJ008* |
| 2820 | 0.65 | 0.95 | 0.005 | NA | 177 | 23 | 0.51 | 0.001 | *Pseudomonadota* | *g Devosia* |
| 519 | 0.51 | 0.92 | 0.012 | NA | 177 | 29 | 0.33 | 0.047 | *Bacteroidota* | *f Cyclobacteriaceae* |
| 523 | 0.51 | 0.90 | 0.015 | 0.50 | 178 | 32 | 0.43 | 0.008 | *Bacteroidota* | *f Cyclobacteriaceae* |
| 2096 | 0.47 | 0.92 | 0.012 | 0.61 | 181 | 18 | 0.55 | 0.000 | *Planctomycetota* | *f Planctomycetaceae* |
| 849 | 0.53 | 0.92 | 0.009 | NA | 181 | 28 | 0.42 | 0.009 | *Pseudomonadota* | *g Luteimonas D* |
| 1527 | 0.56 | 0.93 | 0.007 | NA | 183 | 27 | 0.48 | 0.003 | *Pseudomonadota* | *f Sphingomonadaceae* |
| 1578 | 0.49 | 0.97 | 0.009 | NA | 184 | 24 | 0.40 | 0.015 | *Pseudomonadota* | *f Sphingomonadaceae* |
| 618 | 0.52 | 0.90 | 0.013 | NA | 187 | 32 | 0.39 | 0.018 | *Planctomycetota* | *f Lacipirellulaceae* |

*Continued on next page*

| ASV | PBC | ISA | MAS | ALX | Rank | n | rho | p | Phylum | Lowest taxonomy |
|---|---|---|---|---|---|---|---|---|---|---|
| 320 | 0.52 | 0.88 | 0.017 | NA | 191 | 35 | 0.33 | 0.044 | *Actinomycetota* | *s SZUA-217 sp004356805* |
| 452 | 0.40 | 0.95 | 0.017 | NA | 196 | 36 | 0.45 | 0.005 | *Actinomycetota* | *g Streptomyces* |
| 561 | 0.30 | 0.99 | 0.016 | NA | 197 | 29 | 0.48 | 0.003 | *Bacteroidota* | *g Flavobacterium* |
| 689 | 0.54 | 0.91 | 0.008 | NA | 199 | 30 | 0.42 | 0.009 | *Chloroflexota* | *f Caldilineaceae* |
| 1034 | 0.30 | 0.99 | 0.014 | 0.42 | 202 | 26 | 0.57 | 0.000 | *Actinomycetota* | *s VFJN01 sp009694375* |
| 467 | 0.31 | 0.99 | 0.014 | NA | 202 | 28 | 0.52 | 0.001 | *Bacteroidota* | *s Algoriphagus terrigena* |
| 1762 | 0.55 | 0.91 | 0.007 | NA | 204 | 24 | 0.41 | 0.011 | *Actinomycetota* | *f Cellulomonadaceae* |
| 2427 | 0.53 | 0.94 | 0.006 | NA | 205 | 27 | 0.37 | 0.024 | *Hydrogenedentota* | *f SLHB01* |
| 438 | 0.45 | 0.93 | 0.013 | NA | 205 | 35 | 0.31 | 0.060 | *Pseudomonadota* | *s Paracoccus kondratievae* |
| 411 | 0.52 | 0.87 | 0.012 | 0.59 | 208 | 33 | 0.31 | 0.060 | *Pseudomonadota* | *c Gammaproteobacteria* |
| 808 | 0.47 | 0.92 | 0.011 | NA | 209 | 26 | 0.31 | 0.061 | *Gemmatimonadota* | *f Gemmatimonadaceae* |
| 159 | 0.51 | 0.92 | 0.008 | NA | 211 | 26 | 0.32 | 0.055 | *Pseudomonadota* | *o Pseudomonadales* |
| 645 | 0.49 | 0.92 | 0.009 | NA | 212 | 30 | 0.36 | 0.028 | *Pseudomonadota* | *s ZC4RG30 sp003242275* |
| 799 | 0.41 | 0.99 | 0.009 | NA | 215 | 21 | 0.41 | 0.012 | *Bacteroidota* | *g Flavobacterium* |
| 721 | 0.45 | 0.93 | 0.011 | NA | 215 | 28 | 0.29 | 0.084 | *Bacteroidota* | *g PHOS-HE28* |
| 1498 | 0.59 | 0.90 | 0.006 | NA | 217 | 31 | 0.44 | 0.006 | *Pseudomonadota* | *s Methylosinus sp000685825* |
| 1301 | 0.42 | 0.98 | 0.009 | NA | 217 | 30 | 0.44 | 0.006 | *Bacteroidota* | *c Bacteroidia* |
| 752 | 0.38 | 0.91 | 0.014 | 0.65 | 219 | 21 | 0.45 | 0.005 | *Bacteroidota* | *s Membranicola marinus A* |
| 154 | 0.53 | 0.85 | 0.016 | NA | 219 | 37 | 0.41 | 0.013 | *Pseudomonadota* | *f Sphingomonadaceae* |
| 327 | 0.41 | 0.97 | 0.010 | NA | 219 | 31 | 0.26 | 0.114 | *Pseudomonadota* | *s Pseudoxanthomonas suwonensis A* |
| 981 | 0.46 | 0.90 | 0.011 | 0.58 | 221 | 31 | 0.48 | 0.002 | *Pseudomonadota* | *o Rhizobiales* |
| 329 | 0.46 | 0.91 | 0.012 | NA | 221 | 36 | 0.28 | 0.098 | *Pseudomonadota* | *f Rhizobiaceae* |
| 543 | 0.53 | 0.86 | 0.013 | NA | 222 | 22 | 0.38 | 0.022 | *unclassified* | *unclassified* |
| 292 | 0.37 | 0.95 | 0.013 | NA | 222 | 31 | 0.32 | 0.051 | *Pseudomonadota* | *o Pseudomonadales* |
| 465 | 0.41 | 0.94 | 0.011 | NA | 223 | 28 | 0.32 | 0.054 | *Pseudomonadota* | *f Methylophilaceae* |
| 2552 | 0.43 | 0.94 | 0.009 | 0.49 | 227 | 19 | 0.54 | 0.001 | *Bacteroidota* | *c Bacteroidia* |
| 974 | 0.66 | 0.85 | 0.008 | NA | 231 | 29 | 0.44 | 0.007 | *Chloroflexota* | *s WHTK01 sp009377745* |
| 232 | 0.41 | 0.90 | 0.017 | NA | 231 | 33 | 0.22 | 0.186 | *OLB16* | *f SURF-12* |
| 4289 | 0.61 | 0.91 | 0.004 | NA | 235 | 25 | 0.37 | 0.023 | *Chloroflexota* | *g W-Chloroflexi-9* |
| 3177 | 0.60 | 0.88 | 0.006 | NA | 236 | 22 | 0.42 | 0.010 | *Deinococcota* | *g JAAYYF01* |
| 2377 | 0.64 | 0.89 | 0.005 | NA | 236 | 30 | 0.42 | 0.010 | *Pseudomonadota* | *f Ferrovibrionaceae* |
| 3314 | 0.58 | 0.90 | 0.005 | NA | 237 | 17 | 0.48 | 0.003 | *Bacteroidota* | *g PHOS-HE28* |
| 418 | 0.47 | 0.91 | 0.009 | NA | 237 | 28 | 0.23 | 0.170 | *Pseudomonadota* | *o Pseudomonadales* |
| 780 | 0.43 | 0.92 | 0.010 | NA | 239 | 20 | 0.46 | 0.004 | *Bacteroidota* | *s Flavobacterium marinum* |
| 2887 | 0.50 | 0.90 | 0.008 | NA | 241 | 20 | 0.44 | 0.007 | *Gemmatimonadota* | *o Longimicrobiales* |
| 280 | 0.52 | 0.81 | 0.019 | NA | 241 | 20 | 0.35 | 0.033 | *Bacteroidota* | *s Sphingobacterium sp900163865* |
| 406 | 0.50 | 0.88 | 0.010 | NA | 245 | 34 | 0.27 | 0.107 | *Pseudomonadota* | *s Pseudorhizobium flavum* |
| 1734 | 0.49 | 0.90 | 0.008 | NA | 247 | 26 | 0.42 | 0.009 | *Actinomycetota* | *f Ilumatobacteraceae* |
| 2741 | 0.69 | 0.87 | 0.005 | NA | 249 | 31 | 0.46 | 0.005 | *Actinomycetota* | *f Ilumatobacteraceae* |
| 4916 | 0.54 | 0.85 | 0.007 | 0.60 | 254 | 13 | 0.48 | 0.003 | *Planctomycetota* | *f Planctomycetaceae* |
| 867 | 0.50 | 0.91 | 0.006 | NA | 254 | 21 | 0.43 | 0.007 | *Sumerlaeota* | *f Sumerlaeaceae* |
| 2031 | 0.38 | 0.96 | 0.008 | NA | 254 | 26 | 0.40 | 0.014 | *Sumerlaeota* | *s JAIBAT01 sp019695055* |
| 2488 | 0.59 | 0.88 | 0.005 | NA | 255 | 21 | 0.41 | 0.011 | *Actinomycetota* | *f Ilumatobacteraceae* |
| 737 | 0.40 | 0.88 | 0.017 | NA | 260 | 19 | 0.45 | 0.006 | *Actinomycetota* | *g Streptomyces* |
| 3041 | 0.45 | 0.94 | 0.006 | NA | 261 | 24 | 0.38 | 0.019 | *Verrucomicrobiota* | *s Synoicihabitans sp023386835* |

*Continued on next page*

| ASV | PBC | ISA | MAS | ALX | Rank | n | rho | p | Phylum | Lowest taxonomy |
|---|---|---|---|---|---|---|---|---|---|---|
| 998 | 0.53 | 0.88 | 0.006 | NA | 263 | 33 | 0.25 | 0.142 | *Acidobacteriota* | f SCN-69-37 |
| 1058 | 0.55 | 0.87 | 0.006 | NA | 265 | 34 | 0.40 | 0.014 | *Actinomycetota* | *f Ilumatobacteraceae* |
| 1927 | 0.50 | 0.89 | 0.007 | NA | 266 | 28 | 0.43 | 0.009 | *Pseudomonadota* | *s Brevundimonas basaltis* |
| 3139 | 0.59 | 0.87 | 0.005 | NA | 267 | 17 | 0.38 | 0.022 | *Pseudomonadota* | *o Pseudomonadales* |
| 453 | 0.52 | 0.85 | 0.009 | NA | 268 | 30 | 0.35 | 0.034 | *Planctomycetota* | *s JAGQOG01 sp020427695* |
| 1594 | 0.45 | 0.91 | 0.007 | NA | 270 | 24 | 0.42 | 0.010 | *Bacteroidota* | *s Flavipsychrobacter sp020636495* |
| 3416 | 0.49 | 0.90 | 0.006 | NA | 272 | 24 | 0.44 | 0.007 | *Actinomycetota* | *o Acidimicrobiales* |
| 1938 | 0.51 | 0.89 | 0.006 | NA | 274 | 20 | 0.43 | 0.007 | *Pseudomonadota* | *f Rhodobacteraceae* |
| 161 | 0.47 | 0.91 | 0.006 | NA | 274 | 32 | 0.18 | 0.280 | *Pseudomonadota* | *s Halopseudomonas formosensis* |
| 778 | 0.41 | 0.91 | 0.008 | NA | 276 | 26 | 0.34 | 0.041 | *Pseudomonadota* | *g Cellvibrio* |
| 19318 | 0.56 | 0.87 | 0.005 | NA | 276 | 32 | 0.31 | 0.066 | *Chloroflexota* | *c Anaerolineae* |
| 414 | 0.56 | 0.83 | 0.007 | NA | 277 | 35 | 0.26 | 0.118 | *Pseudomonadota* | *c Alphaproteobacteria* |
| 1583 | 0.41 | 0.87 | 0.010 | 0.61 | 281 | 15 | 0.41 | 0.011 | *Bacteroidota* | *g Parapedobacter* |
| 784 | 0.40 | 0.89 | 0.011 | NA | 281 | 22 | 0.30 | 0.071 | *Myxococcota* | *f Polyangiaceae* |
| 1749 | 0.45 | 0.90 | 0.007 | NA | 282 | 30 | 0.40 | 0.015 | *Actinomycetota* | *g Streptomyces* |
| 797 | 0.44 | 0.90 | 0.007 | NA | 287 | 23 | 0.33 | 0.049 | *Pseudomonadota* | *f Burkholderiaceae C* |
| 1085 | 0.69 | 0.82 | 0.005 | NA | 287 | 35 | 0.29 | 0.082 | *Planctomycetota* | *f Planctomycetaceae* |
| 667 | 0.42 | 0.88 | 0.010 | NA | 288 | 34 | 0.37 | 0.023 | *Pseudomonadota* | *f Burkholderiaceae A* |
| 3060 | 0.52 | 0.88 | 0.005 | NA | 289 | 20 | 0.38 | 0.021 | *Pseudomonadota* | *g Novosphingobium* |
| 17109 | 0.53 | 0.87 | 0.005 | NA | 291 | 31 | 0.23 | 0.180 | *unclassified* | *unclassified* |
| 1144 | 0.37 | 0.86 | 0.014 | NA | 303 | 20 | 0.43 | 0.008 | *Bacteroidota* | *f Cyclobacteriaceae* |
| 2258 | 0.56 | 0.79 | 0.006 | NA | 304 | 22 | 0.36 | 0.030 | *Gemmatimonadota* | *f UBA6960* |
| 766 | 0.48 | 0.86 | 0.007 | NA | 309 | 35 | 0.41 | 0.013 | *Pseudomonadota* | *f Sphingomonadaceae* |
| 572 | 0.47 | 0.79 | 0.011 | NA | 310 | 13 | 0.41 | 0.012 | *Actinomycetota* | *g Glycomyces* |
| 9006 | 0.53 | 0.85 | 0.005 | NA | 311 | 26 | 0.25 | 0.136 | unclassified | unclassified |
| 2074 | 0.52 | 0.86 | 0.005 | NA | 312 | 29 | 0.35 | 0.034 | *Chloroflexota* | *g W-Chloroflexi-9* |
| 253 | 0.28 | 0.94 | NA | 0.74 | 319 | 21 | 0.55 | 0.000 | *Actinomycetota* | *g Promicromonospora* |
| 864 | 0.44 | 0.79 | 0.010 | 0.57 | 321 | 10 | 0.44 | 0.007 | *Pseudomonadota* | *s Advenella incenata* |
| 1823 | 0.51 | 0.84 | 0.006 | NA | 321 | 23 | 0.28 | 0.098 | *Pseudomonadota* | *o Pseudomonadales* |
| 1966 | 0.53 | 0.86 | 0.004 | NA | 322 | 29 | 0.33 | 0.043 | *Chloroflexota* | *f Tepidiformaceae* |
| 2769 | 0.40 | 0.84 | 0.009 | 0.60 | 324 | 14 | 0.46 | 0.004 | *Planctomycetota* | *f Planctomycetaceae* |
| 171 | 0.34 | 0.96 | NA | 0.50 | 325 | 31 | 0.39 | 0.017 | *Bacteroidota* | *f Cyclobacteriaceae* |
| 3127 | 0.54 | 0.85 | 0.004 | NA | 326 | 30 | 0.45 | 0.005 | *Chloroflexota* | *f Thermomicrobiaceae* |
| 26612 | 0.48 | 0.87 | 0.005 | NA | 329 | 21 | 0.30 | 0.073 | *Bacteroidota* | *f Saprospiraceae* |
| 1157 | 0.42 | 0.88 | 0.006 | NA | 331 | 27 | 0.41 | 0.011 | *Pseudomonadota* | *f Rhizobiaceae* |
| 1254 | 0.49 | 0.86 | 0.005 | NA | 333 | 32 | 0.25 | 0.132 | *Pseudomonadota* | *f Xanthobacteraceae* |
| 3518 | 0.50 | 0.87 | 0.004 | NA | 336 | 28 | 0.43 | 0.007 | *Chloroflexota* | *s JAIBBD01 sp019694855* |
| 1491 | 0.54 | 0.83 | 0.004 | NA | 338 | 35 | 0.32 | 0.053 | *Bacillota* | *c Bacilli* |
| 2310 | 0.51 | 0.87 | 0.003 | NA | 340 | 31 | 0.31 | 0.059 | *Actinomycetota* | *f Microtrichaceae* |
| 2260 | 0.46 | 0.81 | 0.008 | NA | 342 | 15 | 0.38 | 0.021 | *Pseudomonadota* | *s Pusillimonas D thiosulfatoxidans* |
| 1563 | 0.48 | 0.79 | 0.007 | NA | 342 | 18 | 0.38 | 0.022 | *Pseudomonadota* | *g Sphingobium* |
| 1806 | 0.43 | 0.88 | 0.005 | NA | 343 | 30 | 0.32 | 0.053 | *Pseudomonadota* | *f Beijerinckiaceae* |
| 4088 | 0.55 | 0.83 | 0.003 | NA | 344 | 31 | 0.29 | 0.079 | *Chloroflexota* | s SHYQ01 sp903917615 |
| 1137 | 0.43 | 0.79 | 0.009 | NA | 346 | 12 | 0.42 | 0.010 | *Actinomycetota* | *g Glycomyces* |
| 1484 | NA | 0.86 | 0.008 | 0.50 | 349 | 29 | 0.48 | 0.002 | *Actinomycetota* | *s VFJN01 sp009694375* |

*Continued on next page*

| ASV | PBC | ISA | MAS | ALX | Rank | n | rho | p | Phylum | Lowest taxonomy |
|---|---|---|---|---|---|---|---|---|---|---|
| 2470 | 0.46 | 0.88 | 0.004 | NA | 350 | 31 | 0.28 | 0.098 | *Pseudomonadota* | *f SG8-39* |
| 1391 | 0.52 | 0.83 | 0.004 | NA | 353 | 26 | 0.24 | 0.152 | *Pseudomonadota* | *f Methylophilaceae* |
| 4977 | 0.48 | 0.86 | 0.004 | NA | 360 | 18 | 0.34 | 0.037 | *Myxococcota* | *g Enhygromyxa* |
| 1232 | 0.48 | 0.83 | 0.005 | NA | 361 | 32 | 0.16 | 0.342 | *Acidobacteriota* | f UBA2999 |
| 587 | 0.48 | 0.82 | 0.005 | NA | 367 | 33 | 0.17 | 0.309 | *Pseudomonadota* | *s Ferrovibrio terrae* |
| 1488 | 0.39 | 0.86 | 0.006 | NA | 369 | 21 | 0.40 | 0.014 | *Bacteroidota* | *f Cyclobacteriaceae* |
| 1856 | 0.50 | 0.81 | 0.004 | NA | 377 | 19 | 0.33 | 0.046 | *Pseudomonadota* | *f Sphingomonadaceae* |
| 5670 | 0.48 | 0.84 | 0.004 | NA | 377 | 25 | 0.17 | 0.328 | *Myxococcota* | s MED-G138 sp019637095 |
| 2600 | 0.42 | 0.82 | 0.006 | NA | 381 | 22 | 0.31 | 0.058 | *Verrucomicrobiota* | *s Synoicihabitans sp023386835* |
| 1223 | 0.44 | 0.83 | 0.005 | NA | 384 | 20 | 0.22 | 0.194 | *Pseudomonadota* | *f Burkholderiaceae C* |
| 2047 | 0.35 | 0.85 | 0.006 | NA | 385 | 19 | 0.36 | 0.026 | *Verrucomicrobiota* | *g Cephaloticoccus* |
| 3775 | 0.46 | 0.78 | 0.005 | NA | 391 | 20 | 0.32 | 0.055 | *Chloroflexota* | *g W-Chloroflexi-9* |
| 3145 | 0.49 | 0.76 | 0.004 | NA | 392 | 15 | 0.38 | 0.022 | *Pseudomonadota* | *f Steroidobacteraceae* |
| 2202 | 0.36 | 0.84 | 0.005 | NA | 408 | 22 | 0.34 | 0.039 | *Planctomycetota* | s JADLFV01 sp020849625 |

Table S18: **Bacterial ASVs indicative of the nine most suppressive composts in the cucumber-*R. solani* system.** For abbreviations see Table S15

| ASV | PBC | ISA | MAS | ALX | Rank | n | rho | p | Phylum | Lowest taxonomy |
|---|---|---|---|---|---|---|---|---|---|---|
| 243 | 0.65 | 0.92 | 0.016 | NA | 28 | 37 | 0.56 | 0.000 | *Actinomycetota* | *s Mycobacterium hassiacum* |
| 434 | 0.56 | 0.93 | 0.013 | NA | 43 | 33 | 0.34 | 0.037 | *Pseudomonadota* | *g Rhodomicrobium* |
| 423 | 0.62 | 0.90 | 0.012 | NA | 56 | 36 | 0.40 | 0.013 | *Planctomycetota* | *s UBA2421 sp002343075* |
| 144 | 0.50 | 0.92 | 0.022 | NA | 58 | 35 | 0.24 | 0.146 | *Pseudomonadota* | *f Steroidobacteraceae* |
| 178 | 0.50 | 0.93 | 0.014 | NA | 64 | 34 | 0.51 | 0.001 | *Actinomycetota* | *g Nonomuraea* |
| 13 | 0.48 | 0.92 | 0.089 | 0.53 | 68 | 37 | 0.45 | 0.006 | *Gemmatimonadota* | *o Longimicrobiales* |
| 2784 | 0.74 | 0.94 | 0.005 | NA | 73 | 30 | 0.57 | 0.000 | *Actinomycetota* | *o Acidimicrobiales* |
| 803 | 0.49 | 0.95 | 0.012 | NA | 73 | 30 | 0.45 | 0.005 | *Chloroflexota* | *f UBA6265* |
| 394 | 0.47 | 0.98 | 0.019 | NA | 74 | 34 | 0.45 | 0.005 | *Pseudomonadota* | *o Rhizobiales* |
| 921 | 0.51 | 0.91 | 0.012 | NA | 75 | 31 | 0.37 | 0.022 | *unclassified* | *unclassified* |
| 1101 | 0.53 | 0.92 | 0.008 | NA | 76 | 35 | 0.33 | 0.049 | *Pseudomonadota* | *f Steroidobacteraceae* |
| 114 | 0.65 | 0.84 | 0.018 | NA | 77 | 37 | 0.42 | 0.010 | *Gemmatimonadota* | *c Gemmatimonadetes* |
| 379 | 0.51 | 0.89 | 0.014 | NA | 77 | 35 | 0.34 | 0.039 | *Pseudomonadota* | f HTCC2089 |
| 45 | 0.43 | 1.00 | 0.061 | NA | 80 | 36 | 0.56 | 0.000 | *Gemmatimonadota* | *o Longimicrobiales* |
| 483 | 0.63 | 0.87 | 0.011 | NA | 80 | 35 | 0.50 | 0.001 | *Actinomycetota* | *f Ilumatobacteraceae* |
| 595 | 0.58 | 0.86 | 0.012 | NA | 83 | 26 | 0.37 | 0.024 | *Actinomycetota* | *f Micromonosporaceae* |
| 609 | 0.51 | 0.92 | 0.008 | NA | 83 | 32 | 0.35 | 0.036 | *Actinomycetota* | *g Actinocorallia* |
| 642 | 0.53 | 0.88 | 0.012 | NA | 84 | 33 | 0.42 | 0.009 | *Pseudomonadota* | *s ZC4RG20 sp017577365* |
| 301 | 0.45 | 0.92 | 0.024 | NA | 85 | 31 | 0.41 | 0.011 | *Actinomycetota* | *c Actinomycetia* |
| 10 | 0.52 | 0.85 | 0.054 | NA | 86 | 37 | 0.52 | 0.001 | *Chloroflexota* | *s Sphaerobacter thermophilus* |
| 346 | 0.41 | 0.93 | 0.024 | NA | 88 | 34 | 0.55 | 0.000 | *Pseudomonadota* | *c Gammaproteobacteria* |
| 736 | 0.49 | 0.92 | 0.010 | NA | 88 | 34 | 0.37 | 0.022 | *Pseudomonadota* | *f SG8-39* |
| 4 | 0.51 | 0.85 | 0.065 | NA | 88 | 37 | 0.20 | 0.242 | *Chloroflexota* | *s Aggregatilinea lenta* |
| 1327 | 0.49 | 0.90 | 0.013 | NA | 89 | 31 | 0.37 | 0.022 | *unclassified* | *unclassified* |
| 518 | 0.52 | 0.89 | 0.009 | NA | 92 | 35 | 0.31 | 0.061 | *Deinococcota* | *g JAAYYF01* |
| 8464 | 0.60 | 0.91 | 0.005 | NA | 93 | 32 | 0.59 | 0.000 | *Chloroflexota* | *s Sphaerobacter thermophilus* |
| 298 | 0.47 | 0.90 | 0.019 | NA | 94 | 29 | 0.52 | 0.001 | *Bacteroidota* | *s Anseongella ginsenosidimutans* |
| 3220 | 0.74 | 0.89 | 0.005 | NA | 95 | 32 | 0.58 | 0.000 | *Actinomycetota* | *c Acidimicrobiia* |
| 277 | 0.51 | 0.86 | 0.014 | NA | 95 | 35 | 0.38 | 0.021 | *Acidobacteriota* | *g CADEFD01* |
| 398 | 0.46 | 0.91 | 0.017 | NA | 96 | 35 | 0.53 | 0.001 | *Actinomycetota* | *f ZC4RG35* |
| 891 | 0.49 | 0.89 | 0.012 | NA | 96 | 31 | 0.42 | 0.009 | *Planctomycetota* | *f Lacipirellulaceae* |
| 1249 | 0.51 | 0.89 | 0.008 | NA | 100 | 31 | 0.36 | 0.027 | *Pseudomonadota* | *g Hyphomicrobium C* |
| 1313 | 0.59 | 0.85 | 0.009 | NA | 102 | 27 | 0.42 | 0.009 | *Pseudomonadota* | *g Terrihabitans* |
| 1013 | 0.49 | 0.91 | 0.008 | NA | 103 | 34 | 0.49 | 0.002 | *Pseudomonadota* | *c Alphaproteobacteria* |
| 368 | 0.48 | 0.88 | 0.016 | 0.56 | 103 | 36 | 0.37 | 0.025 | *Bacillota G* | *g Capillibacterium* |
| 1372 | 0.52 | 0.88 | 0.008 | NA | 103 | 28 | 0.36 | 0.027 | *unclassified* | *unclassified* |
| 788 | 0.48 | 0.93 | 0.008 | NA | 104 | 32 | 0.46 | 0.005 | *Bacteroidota* | *f Cyclobacteriaceae* |
| 102 | 0.56 | 0.83 | 0.012 | NA | 109 | 37 | 0.45 | 0.005 | *Actinomycetota* | *s Thermomonospora curvata* |
| 125 | 0.49 | 0.84 | 0.020 | NA | 114 | 37 | 0.40 | 0.014 | *Actinomycetota* | *s ZC4RG17 sp017577575* |
| 537 | 0.45 | 0.92 | 0.009 | NA | 116 | 34 | 0.28 | 0.095 | *Chloroflexota* | *f Anaerolineaceae* |
| 1416 | 0.55 | 0.87 | 0.006 | NA | 117 | 32 | 0.49 | 0.002 | *Actinomycetota* | *f Miltoncostaeaceae* |
| 658 | 0.49 | 0.88 | 0.008 | NA | 119 | 31 | 0.46 | 0.004 | *Actinomycetota* | *g Nonomuraea* |

*Continued on next page*

| ASV | PBC | ISA | MAS | ALX | Rank | n | rho | p | Phylum | Lowest taxonomy |
|---|---|---|---|---|---|---|---|---|---|---|
| 900 | 0.52 | 0.79 | 0.014 | NA | 123 | 23 | 0.31 | 0.058 | *Pseudomonadota* | *g Ga0077530* |
| 105 | 0.55 | 0.81 | 0.010 | NA | 127 | 37 | 0.44 | 0.007 | *Actinomycetota* | *s Thermomonospora curvata* |
| 2729 | 0.55 | 0.86 | 0.005 | NA | 128 | 32 | 0.46 | 0.004 | *Bacillota* | *s Calditerricola satsumensis* |
| 1469 | 0.49 | 0.88 | 0.007 | NA | 130 | 33 | 0.47 | 0.003 | *Planctomycetota* | *s UBA2421 sp002343075* |
| 285 | 0.53 | 0.82 | 0.010 | NA | 132 | 36 | 0.31 | 0.066 | *Chloroflexota* | *p Chloroflexota* |
| 3067 | 0.64 | 0.83 | 0.006 | NA | 134 | 34 | 0.32 | 0.057 | *Chloroflexota* | *s Aggregatilinea lenta* |
| 710 | 0.54 | 0.85 | 0.006 | NA | 134 | 35 | 0.22 | 0.192 | *Pseudomonadota* | *g Filomicrobium* |
| 3809 | 0.55 | 0.84 | 0.006 | NA | 137 | 19 | 0.46 | 0.005 | *Pseudomonadota* | *c Gammaproteobacteria* |
| 3223 | 0.49 | 0.87 | 0.007 | NA | 137 | 35 | 0.31 | 0.058 | *Chloroflexota* | *s Sphaerobacter thermophilus* |
| 794 | 0.45 | 0.89 | 0.008 | NA | 137 | 34 | 0.21 | 0.221 | *Pseudomonadota* | *s Povalibacter uvarum* |
| 1446 | 0.47 | 0.91 | 0.006 | NA | 138 | 33 | 0.51 | 0.001 | *Actinomycetota* | *o Euzebyales* |
| 2055 | 0.50 | 0.88 | 0.005 | NA | 140 | 34 | 0.47 | 0.003 | *Chloroflexota* | *f Thermomicrobiaceae* |
| 791 | 0.44 | 0.89 | 0.008 | NA | 140 | 27 | 0.38 | 0.020 | *Actinomycetota* | *g Streptomyces* |
| 23055 | 0.58 | 0.84 | 0.005 | NA | 140 | 31 | 0.38 | 0.021 | *Chloroflexota* | *s Aggregatilinea lenta* |
| 21324 | 0.60 | 0.85 | 0.003 | NA | 141 | 33 | 0.32 | 0.053 | *Chloroflexota* | *c Anaerolineae* |
| 2403 | 0.49 | 0.86 | 0.007 | NA | 141 | 35 | 0.18 | 0.284 | *Pseudomonadota* | *f Steroidobacteraceae* |
| 269 | 0.46 | 0.86 | 0.011 | NA | 142 | 36 | 0.38 | 0.022 | *Pseudomonadota* | *o Rhizobiales* |
| 749 | 0.38 | 0.86 | 0.015 | NA | 143 | 26 | 0.57 | 0.000 | *Acidobacteriota* | *f SCN-69-37* |
| 2887 | 0.60 | 0.83 | 0.005 | NA | 144 | 20 | 0.59 | 0.000 | *Gemmatimonadota* | *o Longimicrobiales* |
| 1370 | 0.49 | 0.87 | 0.006 | NA | 144 | 30 | 0.49 | 0.002 | *Actinomycetota* | *s VFJN01 sp009694375* |
| 4478 | 0.46 | 0.86 | 0.009 | NA | 147 | 29 | 0.40 | 0.015 | *Gemmatimonadota* | *c Gemmatimonadetes* |
| 2570 | 0.55 | 0.85 | 0.003 | NA | 149 | 37 | 0.47 | 0.003 | *Bacillota A* | *s Caldicoprobacter algeriensis* |
| 2130 | 0.43 | 0.90 | 0.007 | NA | 149 | 29 | 0.41 | 0.013 | *Pseudomonadota* | *c Alphaproteobacteria* |
| 1362 | 0.54 | 0.83 | 0.005 | NA | 157 | 34 | 0.36 | 0.029 | *Actinomycetota* | *o Acidimicrobiales* |
| 21550 | 0.49 | 0.84 | 0.007 | NA | 157 | 35 | 0.24 | 0.150 | *Chloroflexota* | *c Anaerolineae* |
| 523 | 0.40 | 0.84 | 0.013 | NA | 162 | 32 | 0.52 | 0.001 | *Bacteroidota* | *f Cyclobacteriaceae* |
| 3249 | 0.49 | 0.85 | 0.005 | NA | 163 | 32 | 0.31 | 0.059 | *Bacillota A* | *c Clostridia* |
| 3191 | 0.54 | 0.77 | 0.005 | NA | 167 | 15 | 0.49 | 0.002 | *Pseudomonadota* | *s Sinorhizobium fredii* |
| 1024 | 0.41 | 0.88 | 0.006 | NA | 169 | 29 | 0.22 | 0.188 | *Myxococcota* | *f GCA-2862545* |
| 694 | 0.41 | 0.74 | 0.016 | NA | 170 | 17 | 0.53 | 0.001 | *Planctomycetota* | *g Tautonia* |
| 1067 | 0.48 | 0.84 | 0.007 | NA | 170 | 26 | 0.51 | 0.001 | *Actinomycetota* | *g Nonomuraea* |
| 1870 | 0.46 | 0.86 | 0.005 | NA | 176 | 25 | 0.41 | 0.012 | *Pseudomonadota* | *f Sphingomonadaceae* |
| 1594 | 0.44 | 0.85 | 0.007 | NA | 177 | 24 | 0.42 | 0.010 | *Bacteroidota* | *s Flavipsychrobacter sp020636495* |
| 4759 | 0.50 | 0.84 | 0.004 | NA | 178 | 35 | 0.39 | 0.017 | *Actinomycetota* | *s ZC4RG17 sp017577575* |
| 3791 | 0.51 | 0.82 | 0.004 | NA | 182 | 21 | 0.25 | 0.138 | *Bacillota G* | *g Capillibacterium* |
| 2106 | 0.47 | 0.73 | 0.008 | NA | 183 | 15 | 0.36 | 0.028 | *Pseudomonadota* | *s Palsa-892 sp022844205* |
| 23950 | 0.51 | 0.82 | 0.003 | NA | 184 | 29 | 0.33 | 0.047 | *Chloroflexota* | *s Aggregatilinea lenta* |
| 25535 | 0.40 | 0.79 | 0.008 | NA | 197 | 23 | 0.49 | 0.002 | *unclassified* | *unclassified* |
